# Detecting branching rate heterogeneity with tree balance statistics in lineage tracing trees

**DOI:** 10.1101/2024.06.27.601073

**Authors:** Yingnan Gao, Alison F. Feder

## Abstract

Understanding variation in cellular growth rates among cells in tumors is crucial for predicting cancer progression and interpreting tumor-derived genetic data. Advances in lineage tracing technologies enable the reconstruction of high-resolution, single-cell phylogenies of cancer cell populations, but methods to detect cellular growth rate differences on these phylogenies remain limited. Tree balance statistics offer a way forward, but it is unknown if and how these statistics are distorted when applied to phylogenetic reconstructions built from lineage tracing data, and if these distortions limit the utility of tree balance statistics to distinguish between evolutionary scenarios characterized by variable or homogeneous cellular growth rates. Here, we examined two tree balance statistics, *J*^1^ and the Sackin index, and benchmarked their performance in distinguishing lineage tracing trees derived from populations with and without variable cellular growth rates. We found that when tumor population sizes and lineage tracing editing rates are approximately known and in favorable ranges, *J*^1^ detects departures from homogenous growth rates just as well on lineage tracing trees as on true genealogical trees, while the Sackin index loses most of its power even under the most favorable conditions. We applied our *J*^1^-based test to data derived from cancer lineage tracing experiments and found widespread signals of growth rate heterogeneity in murine autochthonous lung cancers, and lung and PDAC xenograft experiments in mice. Our results demonstrate the potential and challenges of tree balance statistics in analyzing growth dynamics in lineage tracing data.

**Significance statement:** Although tree balance statistics are increasingly applied to examine tumor growth dynamics in trees built from bulk sequencing data, their application in tumor lineage tracing studies remains limited. In this study, we benchmarked *J*^1^ and the Sackin index’s potential to distinguish between tumor populations with and without varying growth rates among concurrent lineages using trees reconstructed from lineage tracing experiments. We describe conditions under which *J*^1^ can detect deviations from homogenous growth rates in lineage tracing trees, and find that the Sackin index is not well-suited to this objective. Our findings provide a first look at the complications associated with using tree balance statistics on trees reconstructed from lineage tracing data, and provide a practical guide for the application of these balance statistics in future lineage tracing studies.

## Introduction

Variation in cancer cell division rates drive the expansion of specific cell populations within tumors and determine the genetic patterns observed in cancer sequencing efforts. The extent and timing of cancer growth rate heterogeneity during tumor development is an active area of study, as identifying genetic and transcriptomic changes among tumor cells that lead to faster growth can help pinpoint drivers of cancer malignancy and suggest pathways for targeted therapies. Existing approaches to identify growth rate heterogeneity include staining for differential division markers in different parts of a tumor (Reeves et al. 2018; Zheng et al. 2020; Gaglia et al. 2022) and quantifying the differential expression of hallmark genes across tumor cells or subregions (Fennell et al. 2022; Househam et al. 2022; Martínez-Ruiz et al. 2023). In-depth cancer genome sequencing has further permitted phylogenetic (Lewinsohn et al. 2023; Salehi et al. 2023) and population genetic analyses (Bozic et al. 2010; Williams et al. 2018) to detect growth rate differences from naturally occurring genetic variation during tumor progression. Nearly all tree-based analyses focus on the relationships between major cancer subpopulations or “clones”, and there are limited opportunities to reconstruct higher resolution genealogical relationships (e.g., at the single-cell level) from bulk sequencing data, because the relationships between individual cells cannot be reconstructed. While single-cell DNA sequencing is expanding in a cancer context, these studies can normally only profile a relatively small number of cells (Bian et al. 2018; Baslan et al. 2020; Borgsmüller et al. 2023). The lack of high-resolution genealogical relationships limits the inference of growth dynamics in tumor populations at fine scales, obstructing the identification of genetic and transcriptomic changes that confer increased fitness to tumor cells from single-cell sequencing data.

Recent advances in CRISPR-Cas9-based lineage tracing technologies (Liu et al. 2021; Choi et al. 2022; Lin et al. 2023) provide new opportunities to understand the lineage relationships of cancer populations *in vivo* at much finer resolution without relying on naturally occurring mutations (Simeonov et al. 2021; Yang et al. 2022; Jones et al. 2023). Lineage tracing systems can be artificially introduced into tumor or embryonic cell lines to contain specific genomic regions (target sites) engineered to rapidly mutate via CRISPR-based targeting once initiated. Each site can acquire a single, irreversible insertion or deletion mutation (i.e., an edit) that is passed down during cell division, and multiple target sites can be arranged into barcodes. When mice containing these engineered cells develop autochthonous tumors after induction (Yang et al. 2022) or when engineered tumor cells engraft after injection into a host (Simeonov et al. 2021; Quinn et al. 2021), their barcodes gradually mutate over time as they divide and expand. At the end of an experiment, descendent cancer cells can be recovered from their host, and these barcodes can then be read out from individual cells through single-cell RNA sequencing to reconstruct lineage relationships among the recovered cells.

In lineage tracing experiments, the genealogical relationships among cancer cells can hypothetically be reconstructed down to the single-cell level. In a fully resolved single-cell phylogenetic tree, a tip (or leaf) represents an extant cell that is recovered and sequenced, and an internal node, where an edge branches into two separate edges, represents the cell division through which the latest common ancestor cell of a group of cells divided into two daughter cells. If certain cells divide more rapidly than others, this can generate asymmetric branching topologies in which faster growing cells leave more descendants than slower growing ones. While lineage-tracing experiments have yielded insights into cancer plasticity (Eyler et al. 2020; Simeonov et al. 2021; Yang et al. 2022; Schiffman et al. 2024), metastasis (Quinn et al. 2021; Zhang et al. 2021; Serio et al. 2024) and progression (Yang et al. 2022), the branching structures of lineage tracing trees have not yet been widely exploited to identify heterogeneous growth processes in tumors.

Methods to examine differential growth rates from tree-like data have a long history in evolutionary biology (Kersting et al. 2025), and have increasingly gained traction in the analysis of cancer genetic data. For example, tree balance statistics that describe the degree to which clades divide into subclades of equal size (Shao & Sokal 1990) are increasingly applied to clonal cancer trees (Schwarz et al. 2014; Werner et al. 2017; Jiang & Tomlinson 2020; Scott et al. 2020; Lynch et al. 2022; Salehi et al. 2023; Liu et al. 2024). Other examples include multi-state speciation and extinction models used in macroevolution (Maddison et al. 2007; FitzJohn 2012) that have recently been adapted to look at growth rate differences in hepatocellular carcinomas (Lewinsohn et al. 2023), and branch length-based phylogenetic tree statistics devised for studying viral fitness (Neher et al. 2014) that have been applied to evaluate the differential fitness among lineages in a tumor population (Prillo et al. 2023). However, the application of the above methods to single-cell trees derived from lineage tracing trees has been much more limited, due to underlying data complexities including barcode saturation and a lack of well-established clock models. While there are important advances in modeling more complex clocks to incorporate branch lengths (Feng et al. 2021; Seidel & Stadler 2022; Prillo et al. 2023; Zwaans et al. 2025) and in designing new lineage tracing systems with better resolution (Liu et al. 2021; Choi et al. 2022; Lin et al. 2023), opportunities remain to quantify heterogeneous growth within cancer cell populations through applying tree balance statistics to existing tree topological data. To exploit these opportunities, we require a greater understanding on how tree balance statistics behave on single-cell trees reconstructed from lineage tracing data.

In this paper, we investigated if two tree balance statistics, *J*^1^ and the Sackin index, can be used to detect growth rate heterogeneity in single-cell trees reconstructed from lineage tracing data. We first found that tree reconstruction from lineage tracing data can introduce a number of complications into tree topologies compared to the underlying genealogical truth, including increased tree balance and widespread polytomies that affect the node-to-tip ratio. Despite these complications, the *J*^1^ statistic can detect departures from growth rate homogeneity in lineage tracing trees under multiple models of cancer evolution. Due to its sensitivity to the node-to-tip ratio, the Sackin index cannot. We further investigated *J*^1^’s robustness under parameter misspecification, and found that its performance depends on estimating the lineage tracing editing rate accurately. Finally, we applied *J*^1^ to three existing lineage tracing datasets in tumors, and found widespread heterogeneous growth rates *in vivo*.

## Results

### Branching rate heterogeneity reduces tree balance

We first examined the behavior of two common tree balance statistics in summarizing single-cell tumor phylogenies simulated under heterogeneous growth rates and subject to imperfect tree reconstruction based on lineage tracing. In these simulations, we modeled cell division as a pure birth process in which cells divide at rate λ. At cell division, two new birth processes are generated. This process can be equivalently represented by a single-cell tree in which a cell division is represented by a branching of edges (i.e., an internal node), and branch lengths represent the time between cell divisions or sampling. We modelled cellular growth rate heterogeneity as variation of branching rates among concurrent lineages in the tree.

We examined two models of how branching rates can change along a tree and compared them to a pure birth Yule model in which all cells have the same division rate over time (i.e., an equal branching rate or EBR model) The continuous rate heterogeneity (CRH) model assumes gradual changes of branching rate over time that reflect, for example, frequent but small fitness changes to gene expression. Briefly, the log-transformed branching rate changes over time following Brownian diffusion, and at a split, the two descendant cells inherit the branching rate of their parent. The discrete rate heterogeneity (DRH) model assumes sudden changes of branching rates at split events that reflect large fitness effects by driver mutations. Briefly, right before a split, the branching rate has a chance to either increase or decrease fitness, and the two descendant cells inherit the branching rate of their parent at the split. The strength of branching rate heterogeneity for both the CRH and DRH models is reported below as a rescaled metric of the variance of lineage birth rate per unit time. More details of these models and this parameterization can be found in the Materials and Methods.

Using the three models above, we simulated completely resolved trees each representing the true genealogical history of a full cancer cell lineage. To make these trees more comparable to those derived from lineage tracing experiments, we subsampled living cells uniformly without replacement to represent the experimental processes of tumor excision, cellular dissociation, and cell sorting for a subset of cancer cells to undergo single-cell sequencing. We grew all cancer lineages to 6250 living cells, subsampled 50, 250, or 1250 of those cells, and examined the tree structures that retained only direct ancestors of these subsampled cells.

To understand baseline tree balance attributes of these branching rate heterogeneity models without the added imprecision of tree reconstruction from lineage tracing data, we first computed tree balance on these subsampled trees that otherwise faithfully capture the true genealogical history. Under both CRH and DRH models, branching rates between sister clades diverge over time. Clades with higher branching rates expand faster than those with lower branching rates and CRH and DRH tree topologies often appear imbalanced compared to EBR trees (**Figure 1A**). We quantified this signal using two measures of tree balance, *J*^1^ and the Sackin index. Briefly, *J*^1^ is the average Shannon equitability (normalized information entropy) of the distribution of sister clade sizes at each node, with higher values indicating greater tree balance (Lemant et al. 2022). The Sackin index is the sum of the number of ancestral nodes across all tips, with greater values indicating lesser tree balance (Sackin 1972; Sokal 1983; Shao & Sokal 1990). Both statistics recover tree imbalance resulting from branching rate heterogeneity, and capture that more extreme heterogeneity leads to greater imbalance and greater variance in tree balance between replicates simulated under the same conditions (**Figure 1B**).

Given the divergence in both *J*^1^ and the Sackin index between trees with and without rate heterogeneity, we asked if these statistics could be used as the basis of statistical tests for the presence of rate heterogeneity in lineage tracing trees. Briefly, we calculated the probability that an EBR tree of the same tree size and sampling characteristics as the focal tree has a more imbalanced *J*^1^ statistic or Sackin index than the focal tree, and rejected the null hypothesis of EBR if that probability was smaller than the desired significance level (**Figure 1C**). Because the exact distributions of *J*^1^ and the Sackin index under EBR have not been resolved analytically, we established their null distributions empirically by simulating EBR genealogical trees and calculating the tree balance statistics on the simulated trees. Tests based on both statistics yielded higher power at higher strengths of branching rate heterogeneity, and test power exceeded 0.99 with a branching rate heterogeneity strength of 1.0 in the CRH model and 0.624 in the DRH model (**Figure 1D**). We found that the power of both tree statistics decreased as the tree size (i.e., sampling effort) decreased from 1250 tips (**Figure S1**). Both statistics yielded a type I error rate matching the significance level used in the tests (α = 0. 05, **Figure 1D** and **Figure S1**).

**Figure 1.**
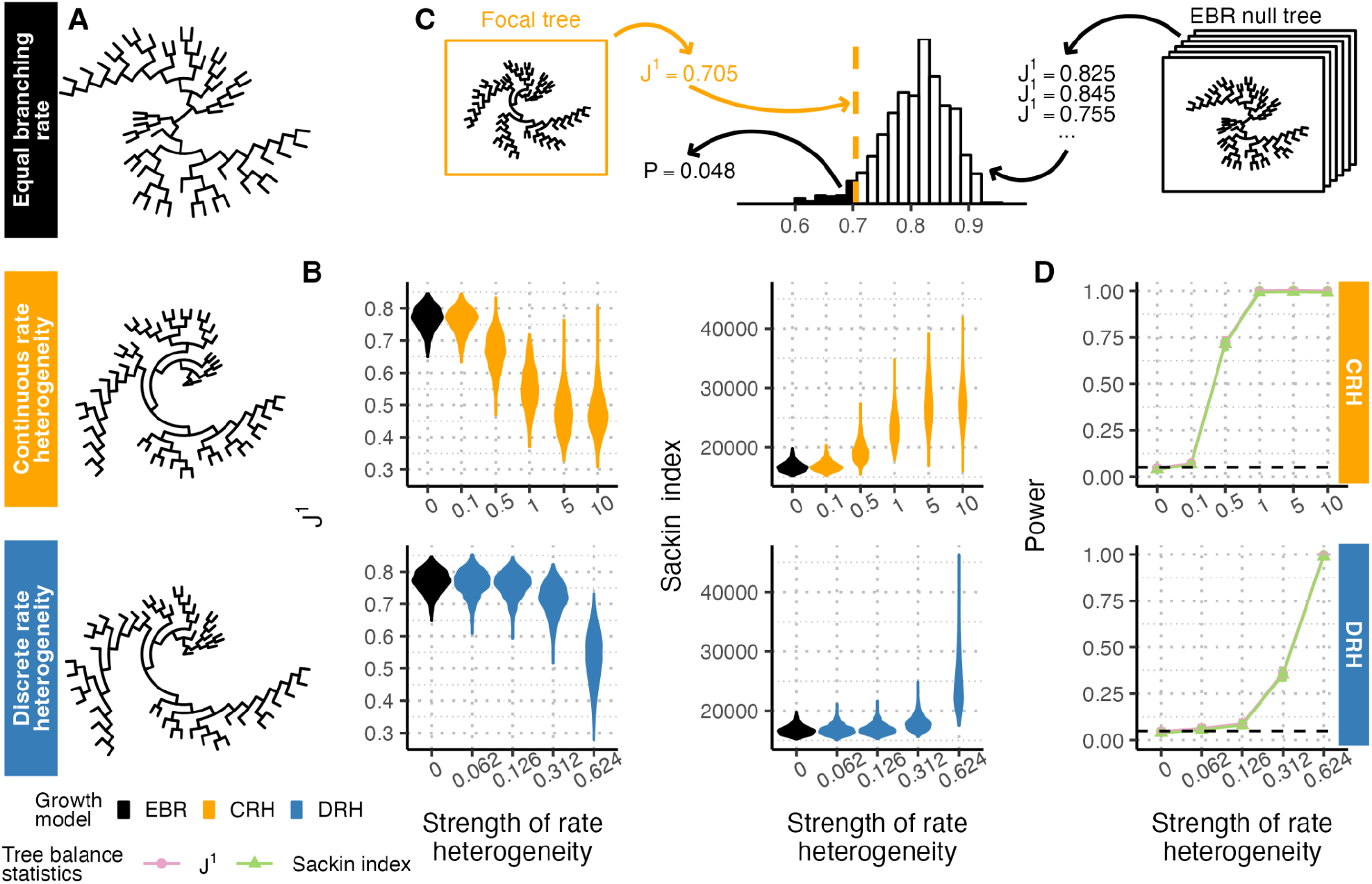
Branching rate heterogeneity affects tree balance as measured by *J*^1^ and the Sackin index. **(A)** Genealogical trees simulated without rate heterogeneity (equal branching rate, or EBR, black) and with rate heterogeneity show visually distinct balance properties. Models of continuous rate heterogeneity (CRH, yellow) and discrete rate heterogeneity (DRH, blue) used a rate heterogeneity strength of 1.0 and 0.624, respectively. Trees are subsampled to 50 tips. **(B)** Distributions of *J*^1^ (left) and the Sackin index (right) for genealogical trees simulated with different strengths of branching rate heterogeneity for CRH (top, rate strengths ϵ {0, 0. 1, 0. 5, 1, 5, 10}) and DRH (bottom, rate strengths ϵ {0, 0. 062, 0. 126, 0. 312, 0. 624}) models. Each distribution summarizes over 1000 replicates. **(C)** Schematic of a statistical approach to compare a focal tree’s balance to an empirical distribution of tree balance under equal branching rates. **(D)** Tests based on *J*^1^ (pink) and the Sackin index (green) are powered to detect departures from EBR in genealogical trees. Solid lines show test power, black dashed lines show the significance level (α = 0. 05), and error bars show 95% confidence intervals from 1000 replicates. Throughout the figure, simulated trees are size 6250 downsampled to 1250 tips unless otherwise noted.

### Lineage tracing trees appear more balanced than their genealogical truths

The fully-resolved single-cell genealogical trees analyzed in the previous section are rarely available for empirical cell populations, and must be inferred (i.e., from lineage tracing barcodes). We can reconstruct the single-cell phylogeny of a cell population from the observed edit states among extant cells using methods suitable to the discrete and irreversible nature of barcode edits (Jones et al. 2020; Simeonov et al. 2021; Yang et al. 2022; Prillo et al. 2023). While tree inference can always yield incorrect lineage relationships between tips, the lineage tracing procedure itself can add additional complications to tree reconstruction. For example, insufficient barcode diversity due to limited barcode length or too low an editing rate can lead to unresolved polytomies, indicating that we cannot fully resolve the genealogical relationships between cells. Both lineage tracing-specific and general tree reconstruction challenges can distort tree topology in lineage tracing trees and undermine the utility of tree balance statistics to detect growth rate heterogeneity as described in the previous section.

To first evaluate the impact of these complications on tree balance, we simulated the reconstruction of lineage tracing trees from the previously simulated true genealogical trees using Cassiopeia (Jones et al. 2020). Briefly, each lineage on the tree (representing an ancestral cell) carries an array of target sites that are edited according to exponential waiting times with mean *t* _*edit*_. Extant cells at the tips of the tree inherit all barcode edits that occur on branches connecting them to the founder cell. Lineage tracing trees are then reconstructed from barcodes of extant cells using Cassiopeia’s MaxCut solver (**Figure 2A**). We examined barcode editing rates (µ_*edit*_ = 1/*t*_*edit*_) that ranged from 0.01 to 1.00, representing an average of 0.09 to 1.00 edits/site, respectively.

We found that different editing rates strongly affected the properties (e.g., node-to-tip ratios) and topologies of lineage tracing trees derived from the same underlying genealogical tree (**Figure 2B-D**). Trees experiencing very fast editing rates relative to their tree length (µ_*edit*_ = 1) captured relationships among the first few cell divisions, but lost resolution as cellular barcodes became fully saturated. As a result, reconstructed lineage trees had widespread star subtrees (defined here as a clade with > 2 tree tips among which each tip is 1 edge away from the clade root) and star subtree-like structures (defined here as a clade with > 2 tree tips among which some but not all tips are 2 edges away from the clade root and the other tips are 1 edge away, examples highlighted in **Figure 2B**). On the other extreme, trees experiencing very slow editing rates relative to their tree length (µ_*edit*_ = 0. 01) also could not fully resolve the relationships between all cells because mutations did not occur at every division. These slow editing rates also led to widespread polytomies (**Figure 2B** and **C**). In both cases, the polytomies induced by these extreme editing rates reduced the node-to-tip ratio from the binary expectation (**Figure 2D**), which affects tree balance statistics that covary with this property (Shao & Sokal 1990). Notably, we found that trees with rate heterogeneity experienced greater reductions in the node-to-tip ratio than EBR trees, especially at lower editing rates. This effect likely emerged because rapidly expanding clades generated by rate heterogeneity formed larger polytomies than observed under equal branching rates. Choosing an editing rate approximately reciprocal to the tree length (here, µ _*edit*_ = 0. 1) minimized these two challenges, reducing polytomies in reconstructed trees (**Figure 2B** and **C**) and better preserving the true node-to-tip ratio than the more extreme editing rates (**Figure 2D**). We observed consistent patterns of these distortions across tree sizes (**Figure S2**).

A separate challenge in these tree reconstructions was that the MaxCut solver reconstructed trees with more balanced roots than those in the true genealogies, which could hide evidence of growth rate heterogeneity. We examined the sister clade sizes at the root of each tree by calculating the log-transformed ratio of the larger clade size over the smaller one. A balanced tree has a log-transformed ratio close to 0 while an imbalanced tree yields a higher value. We found that reconstructed lineage tracing trees have ratios smaller than those of true genealogical trees (indicating greater tree balance), regardless of the lineage tracing editing rate, the underlying growth model, and the tree size (**Figures 2E** and **S2**). These incorrectly reconstructed roots suggest tree building under lineage tracing could mask signatures of the underlying cellular growth dynamics.

We next evaluated the distortion of *J*^1^ and the Sackin index in the reconstructed lineage tracing trees. Specifically, we examined their distributions across different editing rates and divergence in their values between genealogical and lineage tracing trees (**Figure 2F**). Consistent with the other tree topology metrics, both *J*^1^ and the Sackin index peaked and had the smallest variance when the editing rate was approximately reciprocal to the tree length (µ_*edit*_ = 0. 1). While the editing rate affected *J*^1^ non-monotonically, it had similar effects on the trees with and without rate heterogeneity (colored and black violin plots respectively, **Figure 2F**). In contrast, lineage tracing decreased the Sackin index to a greater extent among trees with rate heterogeneity than those without, likely resulting from the Sackin index’s sensitivity to the node-to-tip ratio. Stated another way, the Sackin index captures that trees with rate heterogeneity appear more balanced than those without rate heterogeneity under lineage tracing, in opposition to its findings on the true genealogical trees. This suggests that using higher Sackin indices to detect underlying rate heterogeneity may yield underpowered (or even backwards) results in the context of lineage tracing (see below). We found consistent distortion of *J*^1^ and the Sackin index’s distributions by lineage tracing reconstruction across tree sizes (**Figure S3**).

**Figure 2.**
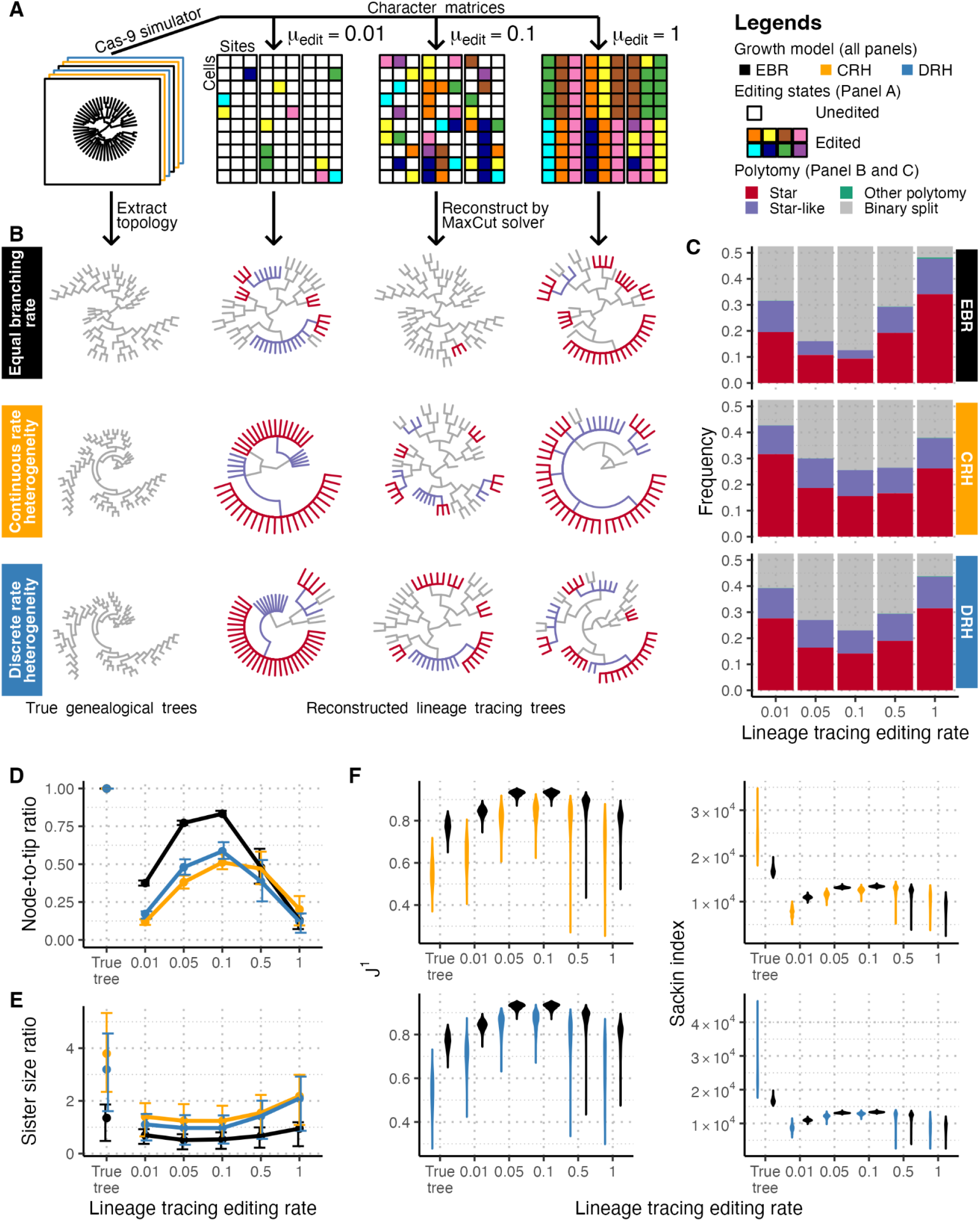
Lineage tracing reconstruction distorts tree topologies and balance. **(A)** To transform simulated genealogical trees into lineage tracing trees, we used the Cas-9 simulator in Cassiopeia to simulate barcode editing, and the MaxCut algorithm to reconstruct tree topologies. Larger rectangles represent editing cassettes composed of individual target sites as squares. Target sites are depicted as white when unedited and colored when edited. **(B)** Reconstructed lineage tracing trees under all three models (rows, EBR, CRH, DRH) have distorted topologies dependent on underlying edit rate (columns, µ_*edit*_ ϵ {0. 01, 0. 1, 1. 0}). Star and star-like polytomies are marked in red and purple, respectively. Trees shown are size 6250 and downsampled to 50 tips for visualization purposes. Reconstructed trees across editing rates (µ_*edit*_ ϵ {0. 01, 0. 05, 0. 1, 0. 5, 1. 0}) and rate heterogeneity models show widespread star andstar-like polytomies **(C)**, decreased node-to-tip ratios **(D)**, and sister clade size ratios **(E)** when compared to true genealogical trees (far left). Error bars represent interquartile ranges. **(F)** Topological distortions also affect *J*^1^ (left) and the Sackin index (right) for both CRH and DRH. Throughout the figure, CRH and DRH trees have a rate heterogeneity strength of 1.0 and 0.624, respectively, and trees are simulated up to size 6250 and downsampled to 1250 tips unless otherwise noted. Each frequency, ratio or distribution summarizes over 1000 replicates.

### *J*^1^ can detect deviations from equal branching rate in reconstructed lineage tracing trees with known parameters

As trees reconstructed from lineage tracing data show distorted tree balance compared to true genealogical trees, we next asked if *J*^1^ and the Sackin index could still detect departures from EBR in formal statistical tests using reconstructed lineage tracing trees. To evaluate their performances, we repeated the formal statistical tests described in the previous section (**Figure 1C**), but with modifications to address the distortion by lineage tracing reconstruction. Specifically, we adjusted the null distributions of *J*^1^ and the Sackin index by repeating lineage tracing simulation and tree reconstruction among the equal branching rate growth processes forming the null. If lineage tracing distortions in the focal tree are reproduced in the EBR null trees through lineage tracing simulation, the contrast between the summary statistics of the focal tree and its adjusted null distribution will only leave the signal of tree balance from the underlying growth model.

For this modified statistical test, we considered first the ideal situation in which the true lineage tracing editing rate and the true population size are known or accurately estimated. Under these conditions and an intermediate editing rate (µ _*edit*_ = 0. 1), *J*^1^ ‘s performance on reconstructed lineage trees was nearly identical to its performance on true genealogical trees (**Figure 3A**), with only a slight loss of power at intermediately strong branching rate heterogeneity. In contrast, the Sackin index had virtually no power to detect deviations from EBR. This loss of power comes from excess distortion of the Sackin index under the combination of lineage tracing reconstruction and rate heterogeneity as described in the previous section. As that distortion diminished when the lineage tracing editing rate increased (**Figure 2D**), we observed a partial restoration in the power of the Sackin index-based tests (**Figure 3B**). However, at higher editing rates, tests based on either statistic had decreased power compared to the genealogical tree baselines (**Figure 3B**), likely due to barcode saturation resulting in high tree reconstruction uncertainty. As visualized in **Figure 2F**, trees both with and without rate heterogeneity have highly overlapping distributions with wide variances for both *J*^1^ and the Sackin index at high editing rates, explaining the limited power. Both *J*^1^ and the Sackin index maintained a type I error rate matching the significance level used in the tests (α = 0. 05, **Figure 3B**). Because of the Sackin index-based test’s overall inability to detect deviations from EBR under even ideal reconstructed lineage tracing conditions, we focus only on the *J*^1^-based test in the analyses that follow.

**Figure 3.**
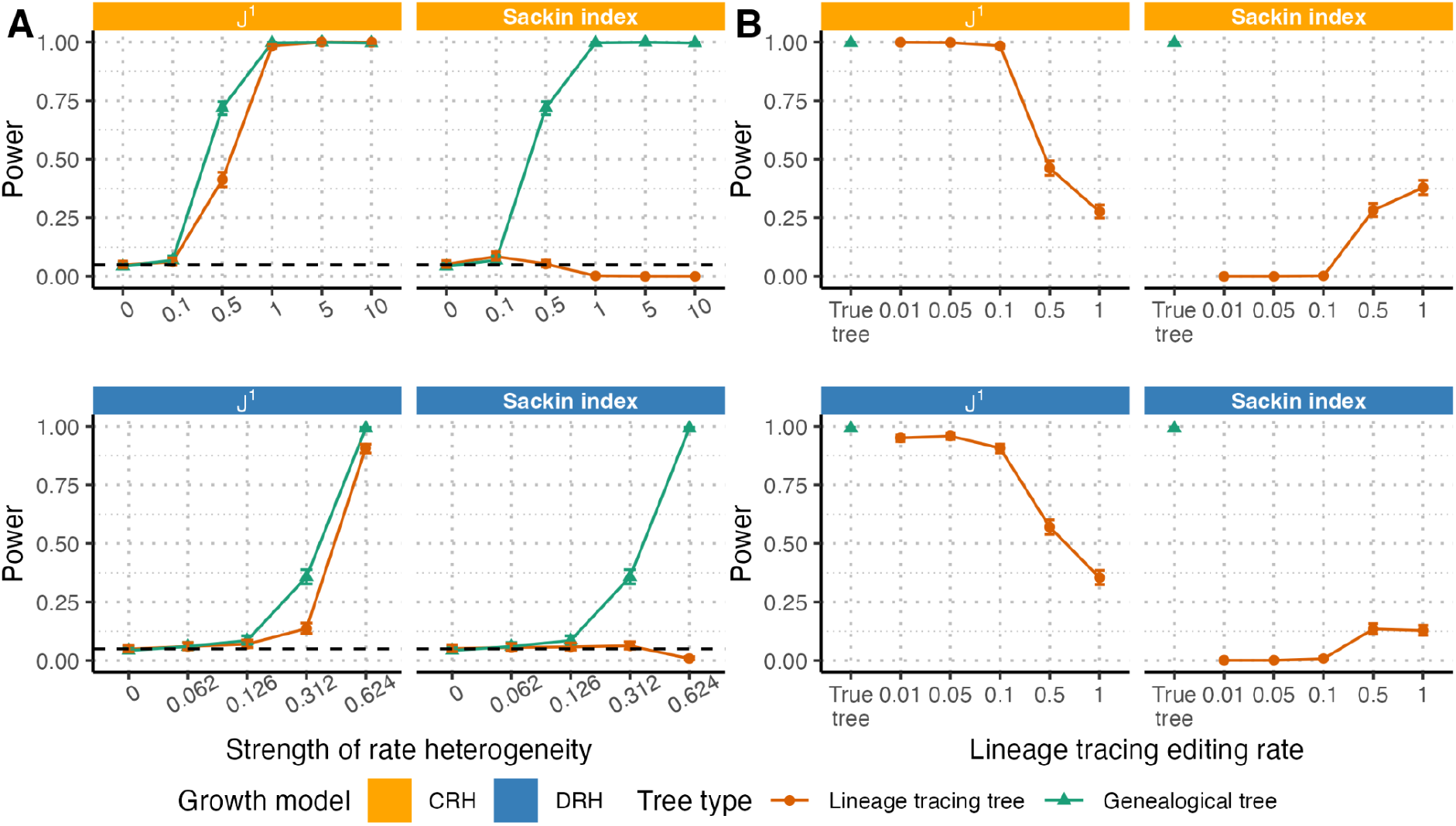
Power of *J*^1^ and Sackin index-based test for branching rate heterogeneity on trees derived from lineage tracing data. For both CRH (top) and DRH (bottom), we evaluated the power of the test illustrated in **Figure 1C** to detect branching rate heterogeneity using *J*^1^ (left) and the Sackin index (right) on reconstructed lineage tracing trees rather than genealogical trees. **(A)** For editing rate µ_*edit*_ = 0. 1, we foundthat while *J*^1^-based tests retained nearly all of their power to detect rate heterogeneity on lineage tracing trees as compared to genealogical trees across different strengths of rate heterogeneity ({0. 1, 0. 5, 1, 5, 10}for CRH and {0. 062, 0. 126, 0. 312, 0. 624} for DRH), Sackin index-based tests did not. **(B)** On trees with strong rate heterogeneity (1.0 and 0.624, for CRH and DRH, respectively) and variable editing rates (µ _*edit*_ ϵ {0. 01, 0. 05, 0. 1, 0. 5, 1}), higher lineage tracing editing rates lowered *J*^1^ test power compared to the test performed on the true trees (left). For the Sackin index, test power was 0 for low editing rates, but increased marginally for higher editing rates. Throughout these figures, solid lines indicate test power, dashed black lines indicate the significance level (α = 0. 05), and error bars indicate 95% confidence intervals from 1000 replicates. All trees are simulated up to size 6250 and downsampled to 1250 tips.

We evaluated the power of *J*^1^-based tests across different tree sizes, and found that sampling fewer tips (i.e., a lower sampling proportion) decreased test power moderately but did not affect the type I error rate (**Figure S4**).

### Parameter misspecification compromises *J*^1^’s power and type I error rate in lineage tracing trees

Because *J*^1^ can effectively distinguish lineage tracing trees with and without branching rate heterogeneity when the editing rate and sampling intensity are known, we next examined *J*^1^’s robustness when those parameters are misspecified. Specifically, parameter misspecification could be important in altering the EBR null distributions against which a focal tree is compared. We therefore repeated the *J*^1^-based statistical test of the previous section, but used null distributions of *J*^1^ calculated from trees with lineage tracing editing rates or sampling intensities that did not match the focal tree.

We first examined the performance of the *J*^1^-based test when the lineage tracing editing rate was misspecified and found that *J*^1^’s power depended sensitively on accurate estimation of the barcode editing rate. When the true editing rate was in an optimal range to capture cellular division dynamics as described above (here, µ_*edit*_ = 0. 1), over- and underestimation of the editing rate decreased both test power and the correctly calibrated type I error rate (**Figure S5**). When the true editing rate was very high or very low (µ_*edit*_ = 0. 01 or µ_*edit*_ = 1), challenges emerging from misspecification were more substantial. The non-monotonicity of theunderlying *J*^1^ EBR null distributions with respect to the editing rate (**Figure 2F**) resulted in non-monotonic power and false positive rates depending on the degree of editing rate misspecification. In particular, false positives rose dramatically when overestimating the low editing rate (µ_*edit*_ = 0. 01) and underestimating the high editing rate (µ_*edit*_ = 1). We therefore conclude that while editing rate misspecification for intermediate editing rates (µ_*edit*_ = 0. 1) can make *J*^1^-based tests for rate heterogeneity moderately more conservative, such tests are inappropriate on highly under- or oversaturated trees unless the editing rates are known.

Examining misspecification on other parameters, we found that the *J*^1^-based test was less sensitive to incorrect estimation of the population size (i.e., sampling intensity) for µ_*edit*_ = 0. 1 and µ_*edit*_ = 1, although it became somewhat more conservative when the population size was underestimated at µ_*edit*_ = 0. 1 (**Figure S5**). In contrast, the slowest editing rate examined (µ_*edit*_ = 0. 01) displayed an evident inflation in the power and false positive rate when its population size was overestimated. This sensitivity to misspecified population size at low editing rates likely derives from differential barcode saturation in larger versus smaller trees causing divergent distributions of *J*^1^ (**Figure S6**). When editing rates are higher, a greater range of tree sizes reach equivalent barcode saturation and therefore have more comparable distributions of *J*^1^. We therefore conclude that robust estimation of the true population size and sampling proportion is less critical than that of the true editing rate, except when the true editing rate is low enough to record insufficient information about cellular relationships.

### Branching rate heterogeneity can lead to underestimation of the editing rate based on character matrix saturation

The sensitivity of the *J*^1^-based test to misspecification of the lineage tracing editing rate underscored the need to accurately estimate lineage tracing editing rates for empirical applications. Ideally, the true editing rate per time unit or per division can be determined experimentally, but this is not always possible. We evaluated a naive strategy to estimate the barcode editing rate in which we matched the observed distribution of barcode saturation (i.e., the proportion of sites that were edited per barcode) in empirical character matrices to simulated character matrices from EBR trees with a range of editing rates, and selected the rate that minimized the differences in barcode saturation between empirical and simulated data.

We first validated that this approach could recover known editing rates in simulated data from EBR trees and those with branching rate heterogeneity. We found that in EBR trees, this approach yielded accurate estimates for editing rates < 0. 5 (**Table S1**), and largely unbiased estimates with moderate error such that most editing rate estimates were no more than 0.1 units away from their true values for editing rates ≥ 0. 5. In trees with strong branching rate heterogeneity (i.e., a strength of 1.0 for CRH and 0.624 for DRH), this naive method underestimated the editing rate for editing rates ≥ 0. 05 for CRH trees and ≥ 0. 1 for DRH trees (**Table S1**).

This effect emerged because populations with rate heterogeneity grew faster than those without, and thus reached their terminal sizes sooner, and consequently yielded fewer recorded edits per barcode. Because underestimation of the editing rate leads to minorly reduced power but does not affect the type I error rate (**Figure S5**), we concluded that using empirically-estimated barcode editing rates was appropriate for µ_*edit*_ ≈ 0. 1.

### Widespread departures from equal branching rate growth in empirical lineage tracing data

We next sought to use this *J*^1^-based test to look for evidence of growth rate heterogeneity in three cancer lineage tracing datasets in lung cancer (Quinn et al. 2021; Yang et al. 2022) and pancreatic ductal adenocarcinoma or PDAC (Simeonov et al. 2021): 1) Yang et al. designed chimeric mice with inducible lung cancer development and lineage tracing via lentiviral vectors, and harvested 73 autochthonous lung tumors 5-6 months after cancer initiation. 2) Quinn et al. implanted human lung cancer cells engineered for lineage tracing into the lungs of mice 4 days after lineage tracing initiation through lentiviral infection, and harvested 148 descendant xenograft lung tumor clonal lineages from the primary injection site 53-80 days after implantation. 3) Simeonov et al. implanted a murine PDAC cell line engineered for lineage tracing into mice via orthotopic injection, induced lineage tracing after 1 week and harvested tumor cells from the primary tumor and five other sites four weeks later. All three experiments used CRISPR-Cas-9-based targeting of ∼30 target barcode edit sites, although see Materials and Methods for more details. Of these tumors, we selected those primary tumors or tumor lineages with at least 100 tips, ultimately yielding 21 tumor trees from the autochthonous lung tumor dataset, 11 clone trees from the xenograft lung tumor dataset, and 5 subclonal trees from the PDAC dataset (**Table S2**, see Materials and Methods).

To obtain the parameters for generating *J*^1^’s EBR null distribution, we estimated the population size and the lineage tracing editing rate separately for each tree in the three datasets. We estimated the population size from the timespan of lineage tracing experiments (see Material and Methods), yielding a population size of 10^6^ for both sets of lung cancer trees and of 31250 for PDAC subclonal trees. We estimated the lineage tracing editing rate for each tree as described in the previous section, yielding rates ranging from 0.2 to 0.4 for autochthonous lung cancer trees, from 0.05 to 0.2 for xenograft lung cancer clone trees, and from 0.2 to 1.0 for PDAC subclonal trees (**Table S2**). Notably, three PDAC tumors had highly saturated barcodes and correspondingly high estimated editing rates (M1_Clone_1_N100, M1_Clone_2_N16, M1_Clone_3_N0). Due to the significant challenges associated with testing for rate heterogeneity under high editing rates as described above, we excluded these three PDAC tumor subclones from our analysis.

Using these estimated parameters, we tested each lineage tracing tree for divergence from an EBR null and found widespread growth rate heterogeneity at the tumor level. Among the autochthonous lung cancer tumors, 19 out of the 21 lung trees showed a significant departure from EBR (**Figure 4**, top 3 rows). We compared these results to Yang et al’s initial analysis of these trees in which they looked for specific subclades undergoing subclonal expansion using the sister clade sizes (Griffiths & Tavaré 1998; Speidel et al. 2019). While our results were largely concordant with Yang et al’s, our scan identified two tumors with a significant departure from EBR that did not have a subclonal expansion identified (3746_NT_T2, 3435_NT_T3). This suggests that our *J*^1^-based test can pick up subtler rate heterogeneity effects that may act on multiple sections of the tree simultaneously rather than only detecting large subclonal expansions restricted to a single clade, although this finding could also be due to differences in the tree reconstruction algorithms used in the two analyses. We also discovered a lack of significant departure from EBR for one tumor identified as having a single large clonal expansion in Yang et al (3703_NT_T2). Examination of this tree revealed a flat topology dominated by one large polytomy, which we demonstrate can occur regularly under models of EBR growth and slow rates of lineage editing (**Figure 2**).

We also found widespread growth rate heterogeneity in xenograft lung tumors at the clonal level, with 9 out of 11 clonal trees showing a significant departure from EBR (**Figure 4**, bottom 2 rows). Quinn et al. did not directly test for growth rate heterogeneity within their trees because the primary aim of the study was characterizing metastatic behavior, but these results are consistent with the autochthonous lung cancer model. We further examined if rate heterogeneity in the primary tumor (as measured by *J*^1^) was associated with metastatic potential (as measured by the rate of outgoing metastatic events, or TreeMetRate in Quinn et al. 2021) and found no correlation between them (Pearson’s ρ = 0. 061, *P* = 0. 858) among the 11 tumors analyzed. Finding little correlation between rate heterogeneity and metastatic behavior is not necessarily surprising because proliferative and metastatic behavior can exploit different or opposed regulatory pathways, and we did not examine their relationships on a cell-by-cell basis.

**Figure 4.**
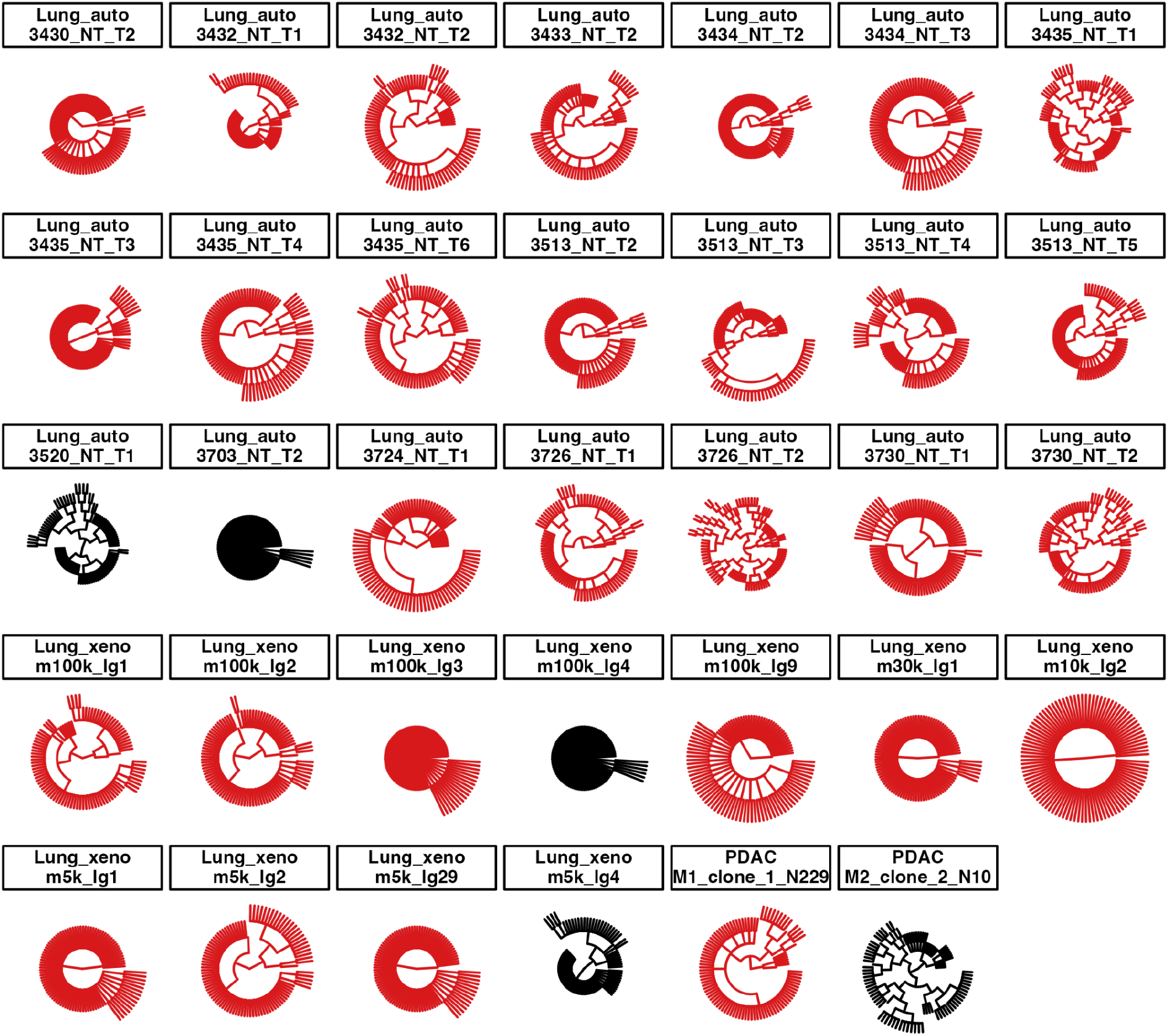
Branching rate heterogeneity is widespread in three *in vivo* lineage tracing datasets. We plot reconstructed tree topologies from three *in vivo* lineage tracing studies of cancer cell lines or autochthonous lung cancers. Trees with a significant departure from EBR are displayed in red, while those without are displayed in black. All trees are uniformly trimmed down to 100 tips for display, but full size trees can be found in **Figure S7**.

We could examine only two PDAC primary tumor trees, and 1/2 showed a significant departure from EBR (M1_Clone_1_N229, **Figure 4**, last 2 panels). The limited number of tumor subclones available with appropriate size and editing characteristics necessary for our test makes it difficult to generalize how widespread growth rate heterogeneity is in this dataset or in PDAC xenografts more generally, but these data do not directly contradict widespread rate heterogeneity in the context of tumor lineage tracing experiments as in the two lung cancer datasets.

## Discussion

Tree-based analyses of cancer genomic data have already yielded extensive insights into cancer evolutionary dynamics (Gerlinger et al. 2012; Schwartz & Schäffer 2017; Casasent et al. 2018; Alves et al. 2019; Lewinsohn et al. 2023). Borrowing from a rich literature preceding mass cancer sequencing (Kersting et al. 2025), tree balance statistics have been applied to suggest departures from homogeneous growth in bulk cancer data (Schwarz et al. 2014; Werner et al. 2017; Chkhaidze et al. 2019; Scott et al. 2020; Noble et al. 2022; Salehi et al. 2023; Liu et al. 2024). However, the application of these tree balance statistics to lineage tracing trees has been limited due to an incomplete understanding of how tree reconstruction from lineage tracing cassettes alters tree balance.

In this paper, we examined tree balance in reconstructed lineage tracing trees through two such statistics, *J*^1^ and the Sackin index, and found multiple challenges with using tree balance in this setting. First, simply building these trees can create excess signatures of tree balance that can hide branching rate heterogeneity. When we used the tree reconstruction algorithm MaxCut from Cassiopeia (Sevillya et al. 2016; Jones et al. 2020) on simulated data, we found that tree roots are reconstructed to be significantly more balanced than their genealogical truth. Future work should examine the degree to which this problem persists with other reconstruction algorithms. Second, because lineage tracing trees have a variety of topological complications resulting from editing rates that are too fast or slow compared to their division dynamics, tree statistics that are highly impacted by polytomies may give misleading evidence of balance even when computed on the same underlying growth process. Further, if these trees are reconstructed from highly saturated barcodes that only cover a small number of early divisions, trees both with and without rate heterogeneity can look highly similar. These factors become challenges when trying to use these statistics as the basis for tests for rate heterogeneity as we explore here. However, we demonstrate that if both the tree reconstruction and editing process can be incorporated into null distributions of neutral growth, signatures of branching rate heterogeneity can still be seen through these lineage tracing-related distortions.

In particular, we noted that *J*^1^ was significantly more robust and consistent when applied to trees reconstructed from lineage tracing data than the more established and widely used Sackin index. This robustness likely relates to *J*^1^’s “universality”, or the degree to which a statistic’s extrema are independent from the underlying tree’s node-to-tip ratio (Lemant et al. 2022). Unlike *J*^1^, the Sackin index is highly sensitive to the node-to-tip ratio and thus examining the same underlying growth process as captured through either high or low editing rates can lead to significantly divergent statistic values.

We note several caveats with our analysis. 1) We established all our growth models on a pure birth process, and assume that rate heterogeneity emerges from variations in birth rates. However, fitness divergence may also arise from variations in death rates, and via more complex modes than the two considered here. Generalizing our growth models to birth-death processes will allow the simulation of cell populations that are more comparable to those observed in empirical data, although we anticipate that many of the challenges associated with lineage tracing trees will persist. 2) Although tree balance statistics are fast and easy to calculate, many different underlying forces can lead to similar statistic values, which limits inferential methods based on tree statistics alone. In our study, the *J*^1^-based test can only detect deviations from EBR, but cannot distinguish if these deviations arise from CRH or DRH, or if their basis is genetic, transcriptional, environmental or a combination thereof. They also cannot reliably quantify the strength of rate heterogeneity.

Likelihood-based methods are likely to be better powered for disentangling these forces and quantifying their strengths. 3) We did not explore how the population size, associated with the amount of time that selection acts, affects the strength of branching rate heterogeneity we can detect in *J*^1^-based tests. Generating a detailed map of the detection limit of rate heterogeneity given tumor population sizes will aid in the interpretation of test outcomes, especially negative ones. 4) We find that *J*^1^-based tests have greatly reduced power when the barcodes are highly saturated, preventing their application to certain empirical datasets. This caveat limits the applicability of *J* ^1^-based tests to only experiments with well-calibrated lineage tracing systems.

In this study, we focused our analyses on eukaryotic CRISPR-Cas9 lineage tracing systems whose sites are independently edited. There are more lineage tracing systems available for investigating population growth dynamics, including systems applicable to bacterial cells (Sheth et al. 2017), those with molecular mechanisms other than CRISPR-Cas9 editing (Pei et al. 2017; Chen et al. 2022), and those with alternative editing behaviours (Choi et al. 2022). Despite these differences, we anticipate that tree-based analyses using lineage tracing data from these systems may face similar challenges to those encountered in this study. We recommend using system-specific simulation to benchmark tree-based analyses before applying them to the empirical data. From the perspective of tree balance, we also encourage the development of tree balance statistics that are inherently robust to lineage tracing to maximize the power to learn the population growth dynamics from lineage tracing data.

## Material and Methods

### Simulating genealogical trees

We simulated genealogical trees of growing tumor populations under three different birth/branching rate models: equal birth/branching rate (EBR), continuous rate heterogeneity (CRH) and discrete rate heterogeneity (DRH). All models employed a pure birth (i.e., Yule) model in which cell divisions occur according to a Poisson process with a rate λ. We used scale-free units of time that could be empirically matched to specific data settings. For our EBR simulations, we used a constant rate of λ = 1 division per unit time for all branches.

For DRH, we modeled the rate of change at cell divisions following Prillo et al. 2023. Specifically, the first lineage possesses a division rate of λ = 1 and each daughter can acquire a mutation at cell division with probability *p*. Mutations have a 0.1 probability of increasing the branching rate by a factor of 2.014 (i.e., λ_*daughter*_ = 2. 014 * λ_*parent*_) and a 0.9 probability of decreasing the branching rate by 7.5% (i.e., λ_*daughter*_ = 0. 925 * λ _*parent*_). We generated DRH trees with four different mutation probabilities *p* ϵ {0. 007, 0. 029, 0. 179, 0. 714}. For the EBR and DRH models, we simulated tumor populations using the birth-death simulator in Cassiopeia (Jones et al. 2020).

For CRH, we modeled that a branch’s λ changes continuously over time rather than only at cell division events following Harvey & Rabosky 2018. Specifically, we modeled an underlying continuous trait *x* for each lineage that evolves via Brownian motion with an arbitrary diffusion coefficient of 0.1. To maintain the DRH model behavior in which the relative increase in the branching rate based on a consistent change in *x* does not change over time, we combined the Brownian change model of Harvey & Rabosky 2018 with an exponential transformation of branching rates as suggested by Kopperud et al. 2023. Specifically, the cellular division rate λ of a lineage is given by an exponential transformation of *x* and a scaling coefficient *c* (λ = *e*^*cx*^). Consistent with the EBR and DRH models, the initial lineage has λ = 1 (i.e., *x* = 0) and we generated CRH trees with five different scaling coefficients *c* ϵ {0. 1, 0. 5, 1. 0, 5. 0, 10}. For the CRH model, we simulated tumor populations using the R package ‘diversitree’ (FitzJohn 2012).

In order to roughly compare the results of rate heterogeneity between the DRH and CRH models, we report both models’ strength of rate heterogeneity as a rescaling of the variance of the log-transformed birth rate normalized by time. If the scaling coefficient, *c*, is larger (under CRH) or if the probability of mutation at a division event, *p*, is greater (under DRH), the birth rate changes more rapidly along a single lineage over time, potentially permitting faster evolutionary rates. We summarize these dynamics by computing the strength of branching rate heterogeneity, *s*, as 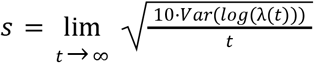, where λ(*t*) is the birth rate of a single lineage at time *t*. Note, this summary of the process is different from the traveling wave dynamics described in Neher & Hallatschek 2013, in which the fitness variance *within* a population over time equilibrates, as the most fit lineages give rise to more fit children. Here, we consider the possible paths of the birth rate itself along a single cellular lineage, and not the realized birth rates among cells in a population. This is for the purpose of summarizing DRH and CRH on the same scales only, and simulated cellular populations display self-reinforcing birth rate dynamics similar to Neher & Hallatschek 2013. In the case of CRH, *s* is the scaling factor *c* directly, as, where *x*(*t*) is Brownian diffusion *Var*(*log*(λ(*t*))) = *Var*(*cx*(*t*)) = *c*^2^ *Var*(*x*(*t*)) = 0. 1*c*^2^ *t* where *x*(*t*) is Brownian diffusion with a diffusion coefficient of 0.1. Thus, for CRH, we report *s* ϵ {0. 1, 0. 5, 1, 5, 10}. In the case of DRH,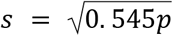 which can briefly be understood as follows: the log birth rate at time *t, log*(λ(*t*)), can be decomposed into the sum of a process that increases by 0.311 on average every *pt* generations (i.e., generations with a mutation) and a process that increases or decreases by 0.389 on average every *pt* generations with probabilities 0.1 and 0.9, respectively. Only this second process contributes to the variance of λ(*t*) across independent lineages, which increases by the single step variance of a biased random walk, 4 · 0. 1 · 0. 9 · 0. 389^2^ = 0. 0545 every *pt* generations. We simulated *p* ϵ {0. 007, 0. 029, 0. 179, 0. 714} to produce *s* ϵ {0. 062, 0. 126, 0. 312, 0. 624}. The EBR model always has *s* = 0.

For all models, we simulated 1000 full genealogical trees up to 6250 tips for each model parameter. To make the genealogical trees more comparable to the lineage tracing trees generated from experimental data, we uniformly subsampled the full genealogical trees. Specifically, we subsampled a full tree down to a size of 50, 250 and 1250 using the uniform tree sampler in Cassiopeia, preserving the root-to-tip distance before and after subsampling. Unless otherwise specified, we refer to these subsampled genealogical trees as genealogical trees in this study.

### Simulating lineage tracing trees

To evaluate the extent to which tree balance is distorted in trees reconstructed from lineage tracing data, we simulated the processes of barcode editing over time on true genealogies and then tree reconstruction from the simulated character matrices. First, we used Cassiopeia’s Cas-9 lineage tracing simulator to simulate the process of barcode editing on a known tree to produce character matrices that could be used in tree reconstruction (Jones et al. 2020). Under this Cas-9 simulator, each target site is independently edited over time with exponential waiting times between edits. Following the design of Yang et al.’s lineage tracing system, we used 10 cassettes of size 3 (i.e., 10 arrays each with 3 target sites) in each cell and 50 possible states for each site. We examined 5 different editing rates (defined as the reciprocal of expected waiting time until an editing event): 0.01, 0.05, 0.10, 0.50 and 1.00. We disabled Cassiopeia’s default Cas-9 resection behavior to avoid excessive missing states, but kept all other parameters at their Cassiopeia defaults. Second, we reconstructed tree topologies (i.e., trees without branch lengths) from the simulated character matrices using the MaxCut solver (Sevillya et al. 2016) in Cassiopeia as Prillo et al. 2023 recently suggested it had optimal performance to accurately reconstruct tree topologies. Unless otherwise stated, we refer to these reconstructed tree topologies as lineage tracing trees in our study.

### Tree balance statistics

In this study, we examined two statistics to quantify tree balance or imbalance in genealogical and lineage tracing trees. First, we examined the Sackin index (Sackin 1972; Sokal 1983; Shao & Sokal 1990), due to its use in non-lineage tracing tumor phylogenetic studies. The Sackin index calculates the sum of the number of ancestors (internal nodes) across all tips in a tree of size *n*:

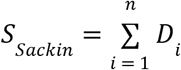

where *i* is the index of tips and *D*_*i*_ is the number of ancestors of the *i*^th^ tip. The Sackin index is compatible with trees with polytomies, and among trees of the same size, a higher Sackin index indicates a less balanced tree.

Second, we examined the *J*^1^ tree balance statistic (Lemant et al. 2022), which calculates the clade-size weighted average Shannon equitability of daughter clade sizes across all *k* internal nodes. In the context of the single-cell genealogical and lineage tracing trees, it is computed as:

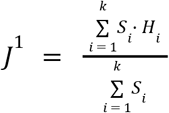

where *S*_*i*_ is the number of tips descending from the focal node *i* and *H* is the Shannon equitability of the daughter clade sizes at the focal node *i*:

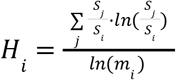

In the above formula, *j* is the index of internal nodes immediately descending from the focal node *i*, and *m*_*i*_ is the outdegree of the focal node *i*. Among trees of the same size, a higher *J*^1^ indicates a more balanced tree, with a perfectly balanced tree having *J*^1^ = 1.

### Establishing null distributions for tree balance statistics

For a genealogical or lineage tracing tree of interest, we tested for the presence of branching rate heterogeneity by calculating the probability that an EBR with a comparable number of tips, population size and lineage tracing conditions is at least as imbalanced as the focal tree. We rejected the null hypothesis of EBR if this probability was smaller than a given significance level. To calculate that probability, we established empirical null distributions by computing the statistic value on 1000 simulated EBR trees of the same type (genealogical or lineage tracing) as the focal tree. EBR nulls were calculated as described above in the **Simulating lineage trees** and **Simulating genealogical trees** sections. To analyze model misspecification and for better fits to the empirical data, we simulated a wider range of underlying true population sizes (1250, 6250, 31250, and 10^6^, subsampled down to sizes of 50, 250 and 1250) and editing rates (0.01, 0.05, 0.1, 0.2, … 1.0) than for the focal trees. We simulated 1000 genealogical trees for each set of parameters, yielding a total of 12,000 EBR genealogical trees and 144,000 EBR lineage tracing trees (4 population sizes, 3 tree sizes, 12 editing rates, and 1000 trees per condition). We calculated *J*^1^ and the Sackin index from these trees to empirically establish their null distributions.

### Benchmarking tree balance statistics in the test against EBR

To evaluate the performance of the two tree balance statistic tests, we benchmarked their power and type I error rate. The power is the probability that the test detects a significant departure from EBR if the focal tree was generated under a model of branching rate heterogeneity. The type I error rate is the probability of detecting a significant departure from EBR if the focal tree was generated under a model of equal branching rates. For power and type I error calculations on genealogical trees, we conducted the statistical test using an empirical null distribution derived from genealogical trees under the same population size and number of tips as the focal tree.

When the focal tree was a lineage tracing tree, we considered first the ideal situation in which the true parameters (i.e., the population size and the lineage tracing editing rate) were known. We conducted the statistical test using a null distribution derived from the lineage tracing trees under the same parameters as the focal tree, and estimated the power and type I error rate. To further test the robustness of tree balance statistics against parameter misspecification, we repeated the above benchmarking with incorrect null distributions derived from lineage tracing trees with different parameters from the focal tree.

### Estimating lineage tracing editing rate from character matrices

To approximately estimate the editing rate from observed cellular barcode matrices when the cellular population size is known but the editing rate is unknown, we calculate the editing rate that produces barcode saturation most comparable to observed data. Specifically, for each cell in the focal character matrix, we calculate the proportion of edited target sites as the number of edited sites divided by the number of non-missing sites (referred to as editing proportion hereafter). We compared the distribution of editing proportions in the focal character matrix to editing proportion distributions in 1000 character matrices derived from size-matched EBR trees with each of 11 different editing rates (0.05, 0.1, 02, 0.3, … 1.0) via the Kolmogorov–Smirnov test. For each editing rate, we calculated the median Kolmogorov-Smirnov D statistic across the 1000 simulated comparisons. We selected the editing rate that minimized the median D statistic, and chose the smaller editing rate in the event of ties.

We benchmarked this approach by re-estimating lineage tracing editing rates in a subset of trees with known rates. We performed the above procedure on 100 lineage tracing trees generated under EBR, CRH and DRH with a tree size of 1250, population size of 6250, and lineage tracing editing rates of 0.01, 0.05, 0.1, 0.5 and 1. For CRH and DRH trees, we examined trees with branching rate heterogeneity strength of 1.0 and 0.624, respectively. We calculated the estimation bias as the mean difference between the true and estimated rates, and the error as the mean absolute difference between the true and estimated rates.

### Empirical data

We applied the *J*^1^-based statistical test to three sets of published tumor lineage tracing data: a murine autochthonous lung cancer dataset (Yang et al. 2022), a murine xenograft lung cancer dataset (Quinn et al. 2021), and a murine xenograft pancreatic ductal adenocarcinoma (PDAC) dataset (Simeonov et al. 2021). We used experimental details surrounding these datasets to establish parameters for tree and lineage tracing simulation in Cassiopeia necessary for performing the test (Jones et al. 2020).

Yang et al produced chimeric mice in which a subset of murine cells carried an inducible lineage tracing system with 4-10 integrated cassettes with 3 target sites each. Exposure of these cells to a lentivirus expressing Cre recombinase both activated conditional oncogenic alleles (initiating tumor development) and induced the expression of Cas9 and associated guide RNAs (initiating lineage tracing via the accumulation of indels in target sites). Five to six months after tumor and lineage tracing initiation, 29 primary tumors were harvested from 11 mice and underwent single-cell RNA sequencing to determine barcode identity. We included 21 in our analysis based on Yang et al’s filtering criteria (tumors that had at least 100 cells, at least 10% of cells reporting a unique set of editing states, and at least 20% of the target sites not having an editing state that was shared by more than 98% of the cells), and analyzed the associated barcode character matrices as described below.

Quinn et al introduced a similar lineage tracing system in a human lung adenocarcinoma cell line (A549) in which engineered cells had 10 integrated cassettes with 3 target sites each. Lineage tracing was initiated *ex vivo* by a lentiviral vector expressing guide RNAs and cells were allowed to expand for four days before implantation into one of four mice (5k, 10k, 30k or 100k injected cells/mouse). Tumorous tissue was collected 53-80 days post-implantation, dissociated, and fluorescence-sorted to retain only tumor cells for single-cell RNA sequencing. We analyzed only cells derived from the initial sites of injection to focus on dynamics in the primary tumors as opposed to metastatic lineages. In total, we examined the character matrices from 11 tumor clone samples with at least 100 cells.

Simeonov et al introduced the macsGESTALT (Simeonov et al. 2021) lineage tracing system into a murine pancreatic ductal carcinoma cell line (6419c5), in which each cell carried between 1 and 13 cassettes of 5 target sites each. Note, on average only 60%-70% of these cassettes were recovered in the descendant cells during sequencing. Two mice were injected with ∼30k cells each and lineage tracing was initiated by doxycycline 1 week after injection. Cells were allowed to expand for four additional weeks before the primary tumor and macroscopic metastases from multiple organs of each mouse were excised. Cells were dissociated and underwent single-cell RNA sequencing to determine barcode identity. We selected clades descending directly from the root of the published clonal trees and filtered out all cells not isolated from primary tumors. In total, we examined the character matrices from five clades with at least 100 cells each, although three of those clades were later excluded due to a high degree of barcode saturation.

From the character matrices generated from the three studies, we used the MaxCut solver in Cassiopeia to infer tree topologies, and computed barcode saturation for inferring the probable editing rate (**Table S2**).

### Testing empirical data against the EBR hypothesis

The lineage tracing editing rates and underlying lineage population sizes were not directly reported in any of the empirical datasets described above. We therefore approximated the population sizes from the experimental designs of the three studies. Under a simple model of exponential growth with 2 day generation times, the three to five month growth periods of cancer cells from Quinn et al and Yang et al would yield cancer cell populations beyond our simulation ability, so we assumed lineage population sizes of 10^6^ for the two lung cancer datasets. The PDAC clones only grew for four weeks after lineage tracing initiation, yielding an approximate population size of 16384 cells under exponential growth. We assumed the closest population size of 31250 for PDAC subclones to employ the null distributions generated in test benchmarking. We note in both cases that our simulated benchmarking suggested that the null distributions used in our *J*^1^-based were relatively robust to population size misspecification. We estimated the lineage tracing editing rate for each tree separately using the method described in the previous section.

Ideally, each focal tree should use a null distribution of *J*^1^ derived from simulated EBR trees with the same number of tips. When empirical trees were small (i.e., ≤ 250 tips), we derived the empirical nulls from previously simulated EBR trees, in which populations were grown to size 31250 or 10^6^, downsampled to 250 tips, underwent lineage tracing simulation and tree reconstruction, and then were further uniformly downsampled to the size of the focal tree. Similarly, when empirical trees had between 251 and 1250 tips, we repeated the above procedures but the simulated EBR trees were first downsampled to 1250 tips before lineage tracing simulation. When empirical trees were large (i.e., > 1250 tips), this procedure was not possible without resimulating lineage tracing and tree reconstruction, so we instead downsampled the focal tree to 1250 tips and used the empirical null distribution of 1250 tip EBR trees directly. Apart from the derivation of the null distributions, the statistical test against EBR was done in the same way as described in previous sections.

## Supporting information

Supplementary Data and Scripts

Suppelmentary Tables

## Data Availability

Scripts used to reproduce the analysis in this study are publicly available at https://github.com/federlab/branching_rate_het.

## Acknowledgements

This study was supported by the National Institutes of Health [1DP2 CA280623-01].

## Supplemental Figures

**Figure S1.**
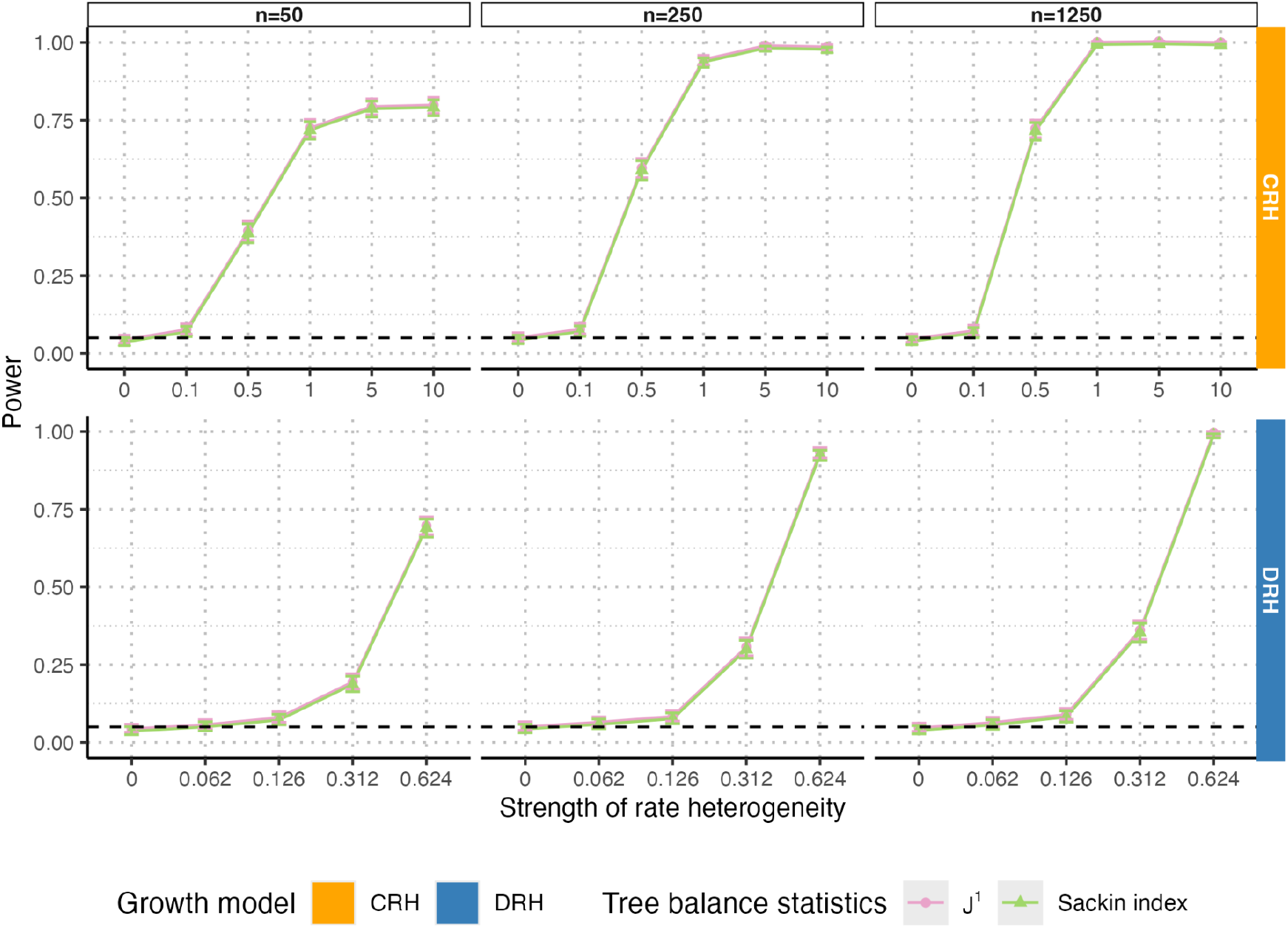
Tests for departures from equal branching rates based on *J*^1^ and the Sackin index have decreased power as the number of tree tips decreases. Power for tests based on *J*^1^ (pink) and the Sackin index (green) under different strengths of branching rate heterogeneity for both continuous rate heterogeneity (yellow, {0, 0. 1, 0. 5, 1, 5, 10}) and discrete rate heterogeneity (blue, {0, 0. 062, 0, 126, 0. 312, 0. 624}). Simulated trees are size 6250 downsampled to 50, 250 and 1250 tips (columns). Solid lines show test power, black dashed lines show the significance level (α = 0. 05), and error bars show 95% confidence intervals over 1000 replicates.

**Figure S2.**
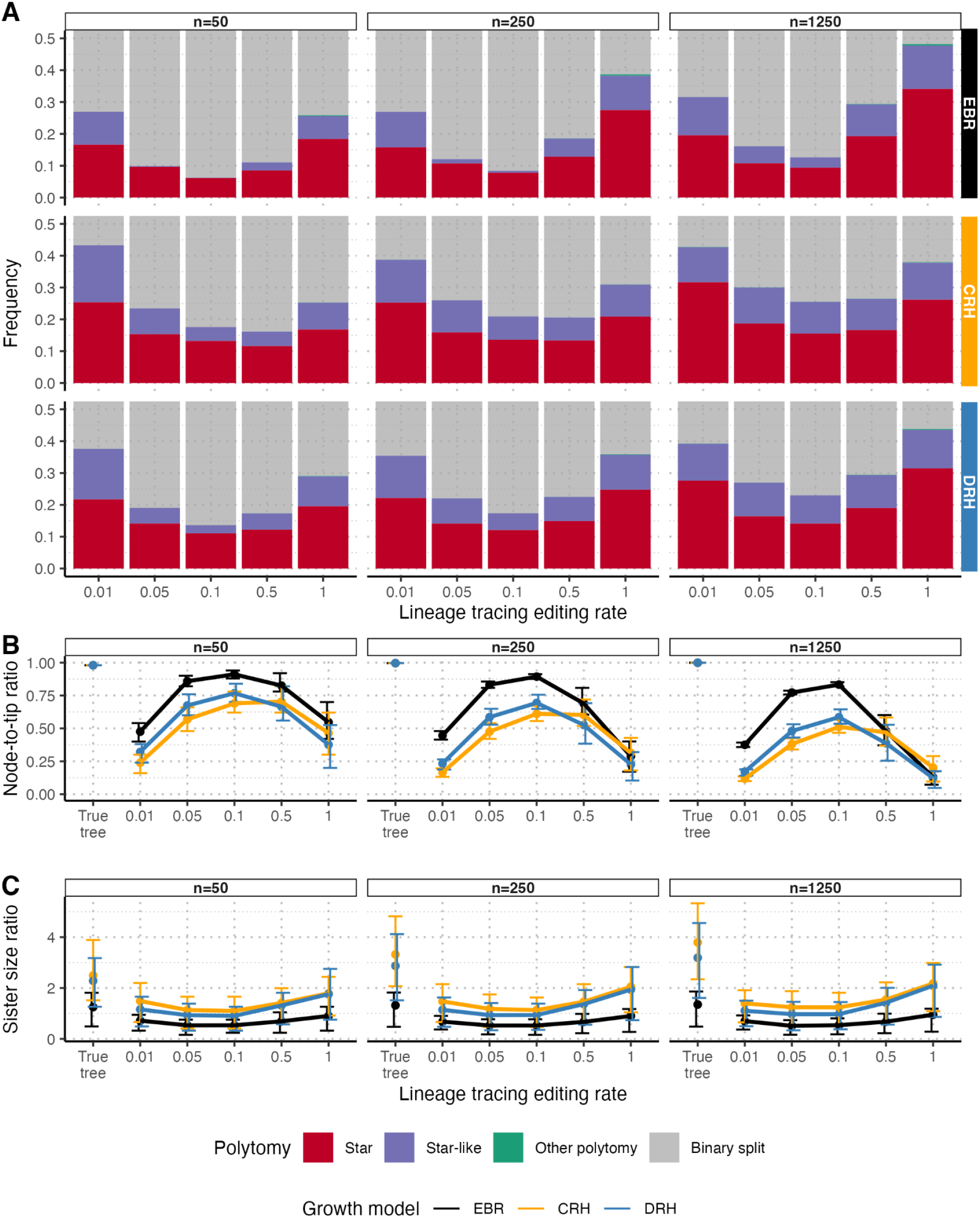
Lineage tracing tree topology distortion is consistent across tree sizes. Reconstructed trees across editing rates (µ_*edit*_ ϵ {0. 01, 0. 05, 0. 1, 0. 5, 1. 0}) and rate heterogeneity models have consistent distortions across tree sizes (*n*_*tips*_ ϵ {50, 250, 1250}) for the proportion of clades that are star or star-like **(A)**, the node-to-tip ratio **(B)**, and the difference in root sister clade size ratios **(C)**. Throughout the figure, CRH and DRH trees have rate heterogeneity strengths of 1.0 and 0.624, respectively, and trees are simulated up to 6250 tips and downsampled to 50, 250 or 1250 tips as noted. Error bars represent interquartile ranges. Each proportion or ratio summarizes over 1000 replicates.

**Figure S3.**
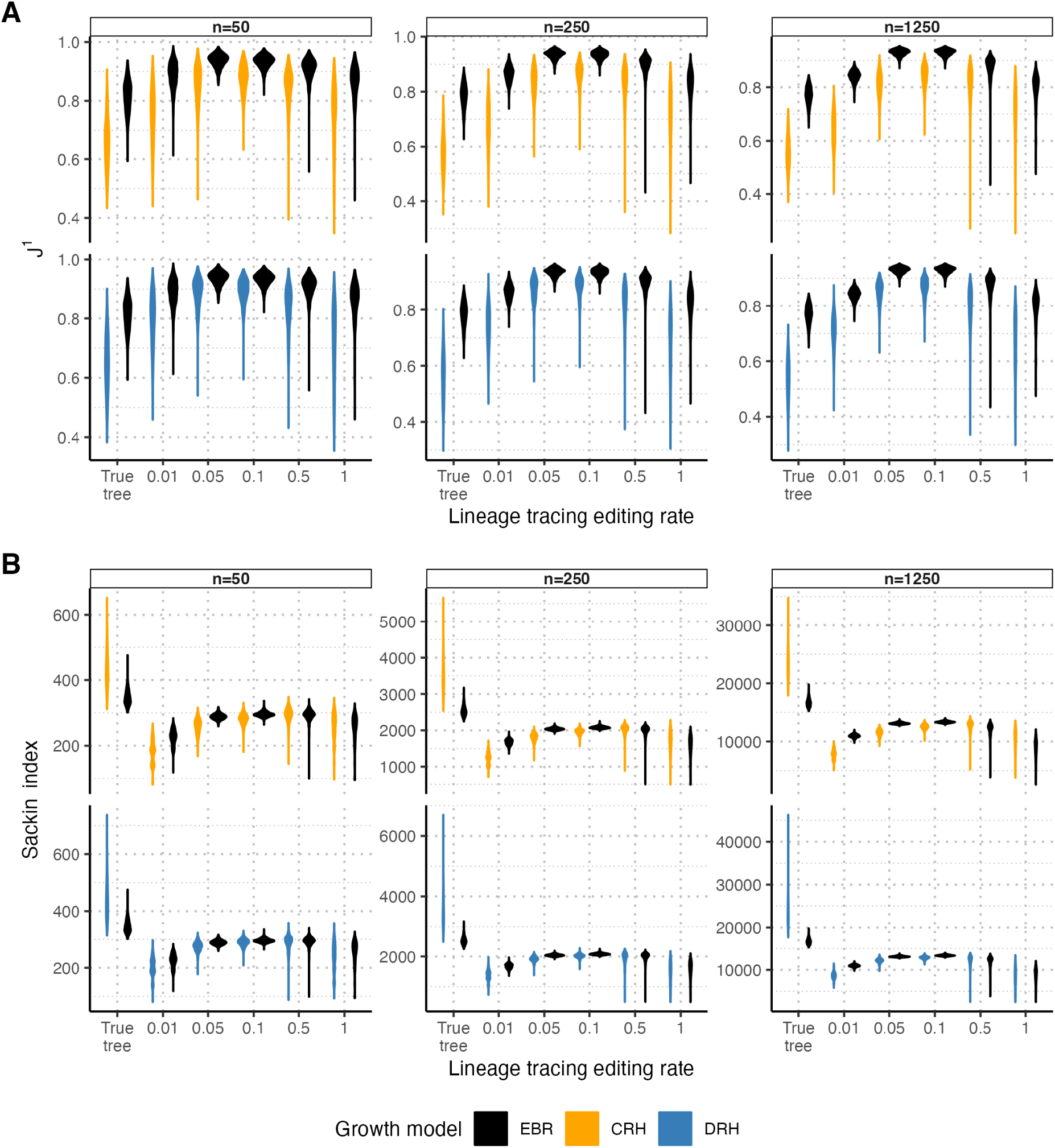
Distributions of *J*^1^ and the Sackin index under different growth models in trees with different numbers of tips. For both *J*^1^ **(A)** and the Sackin index **(B)**, lineage tracing trees across editing rates (µ_*edit*_ ϵ {0. 01, 0. 05, 0. 1, 0. 5, 1. 0}) have distorted tree balance statistics compared to true genealogical trees, but the magnitude of the distortion is relatively insensitive to the number of tree tips (*n*_*tips*_ ϵ {50, 250, 1250}). CRH and DRH trees have a rate heterogeneity strength of 1.0 and 0.624, respectively, and trees are simulated up to 6250 tips and downsampled to 50, 250 or 1250 tips as noted. Each distribution summarizes over 1000 replicates.

**Figure S4.**
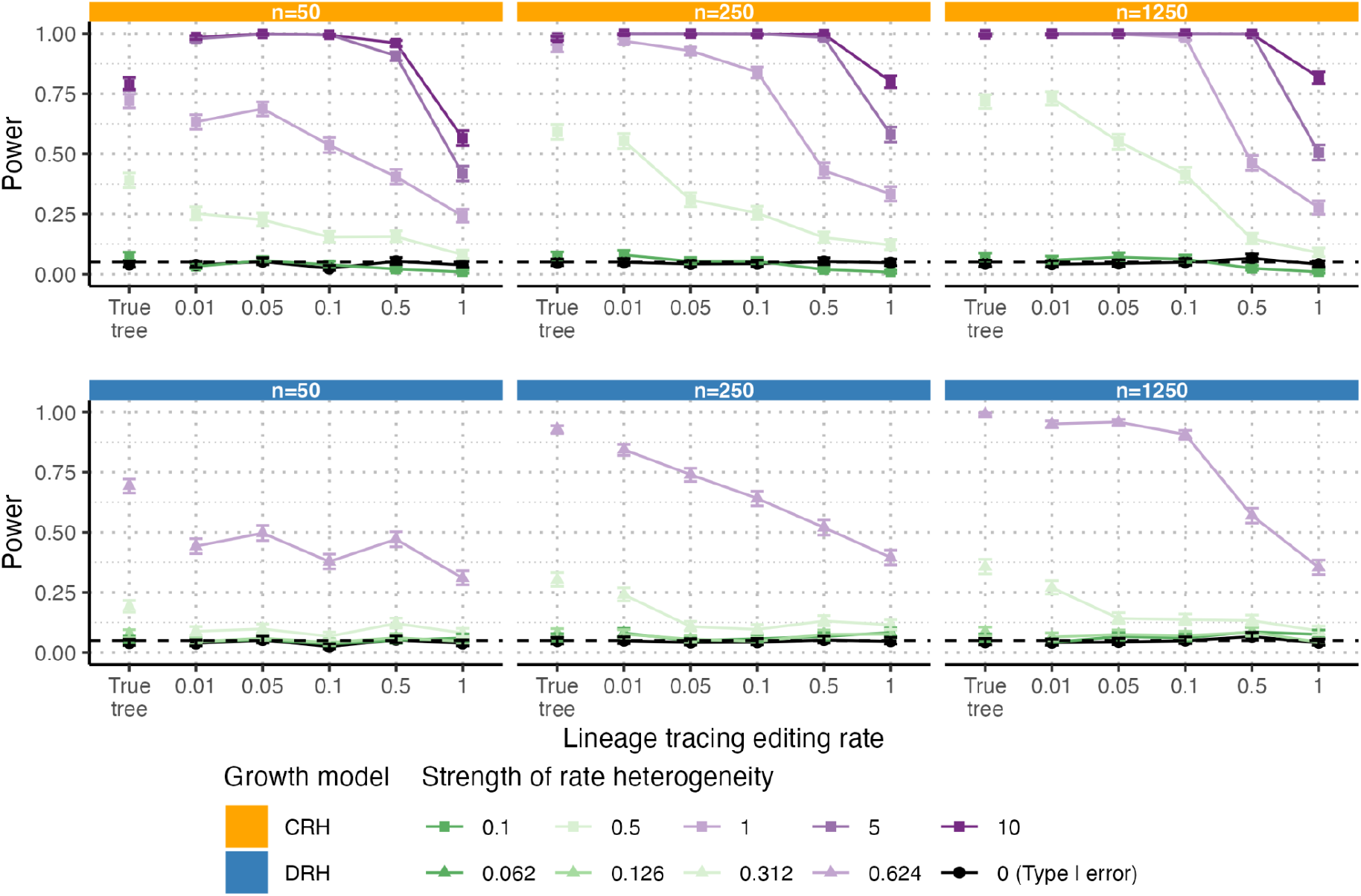
*J*^1^-based test power as a function of tree size and barcode editing rate. The power of the *J*^1^-based test for rate heterogeneity in lineage tracing trees decreases as the editing rate increases for both CRH (top, yellow) and DRH (bottom, blue), among trees with fewer tips (columns, see also **Figure S1**), and at lower strengths of branching rate heterogeneity (colors). Solid lines show test power, black dashed lines show the significance level (α = 0. 05), and error bars show 95% confidence intervals over 1000 replicates. Trees are simulated up to 6250 tips and downsampled to 50, 250 or 1250 tips as noted. The power of the *J*^1^-based tests in true genealogical trees is shown for comparison.

**Figure S5.**
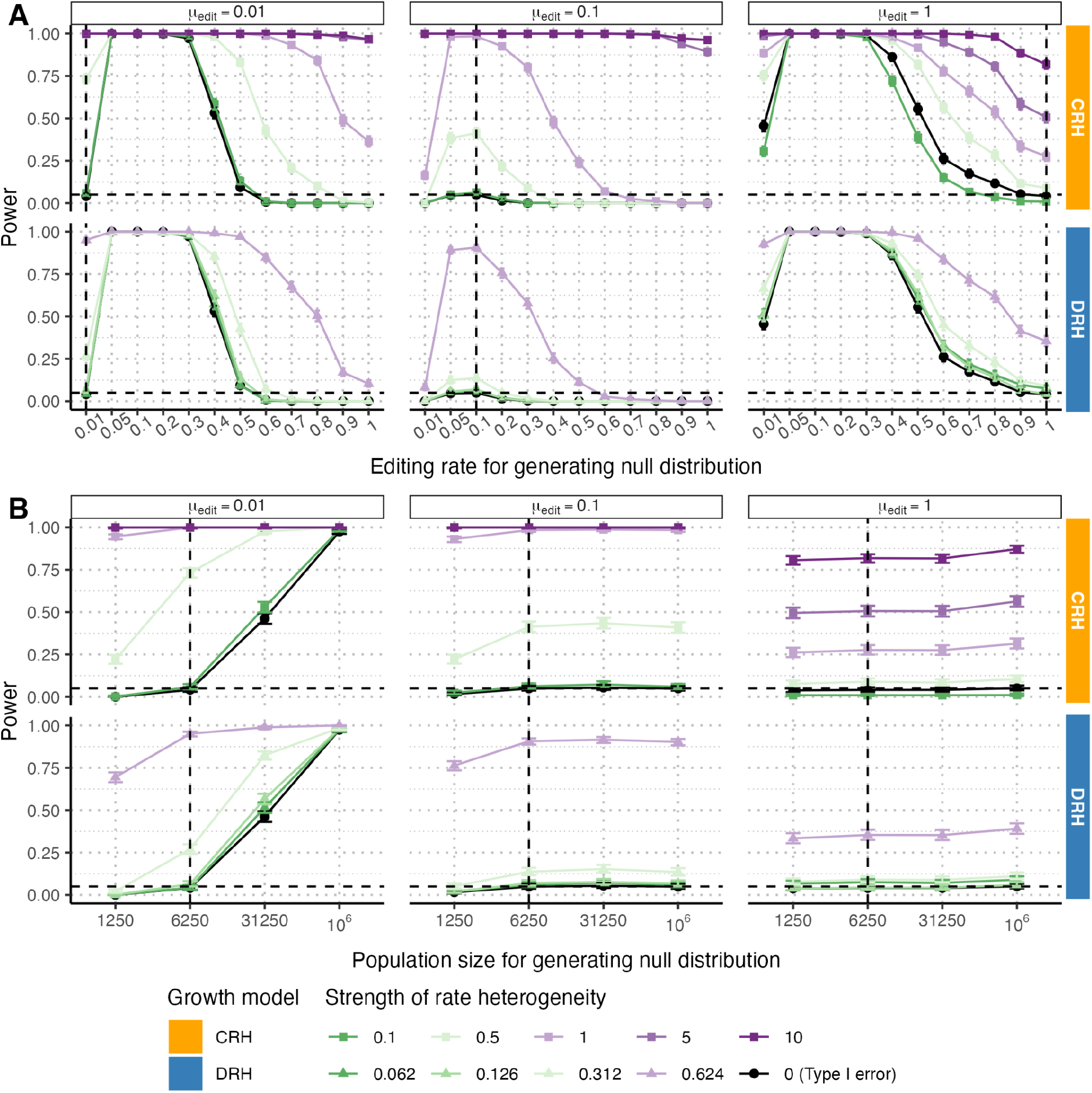
Robustness of *J*^1^’s performance under misspecified parameters for generating EBR null distributions. **(A)** *J*^1^’s power is sensitive to misspecification in the lineage tracing editing rate regardless of the true editing rate (µ_*edit*_ ϵ {0. 01, 0. 1, 1}, in columns and also marked with a vertical black dashed line). **(B)** *J*^1^ ‘s power is sensitive to misspecification in the population size only when the lineage tracing editing rate is low (µ_*edit*_ = 0. 01). EBR null trees are simulated from total population sizes of *n* _*tips*_ ϵ {1250, 6250, 31250, 10_6_}, downsampled to 1250 tips and then used for the basis of comparison for focal trees simulated up to 6250 tips and downsampled to 1250 tips (marked by vertical black dashed line). Solid lines show test power, black horizontal dashed lines show the significance level (α = 0. 05), and error bars show 95% confidence intervals over 1000 replicates. Throughout the figure, trees are simulated up to 6250 tips and downsampled to 1250 tips unless otherwise specified.

**Figure S6.**
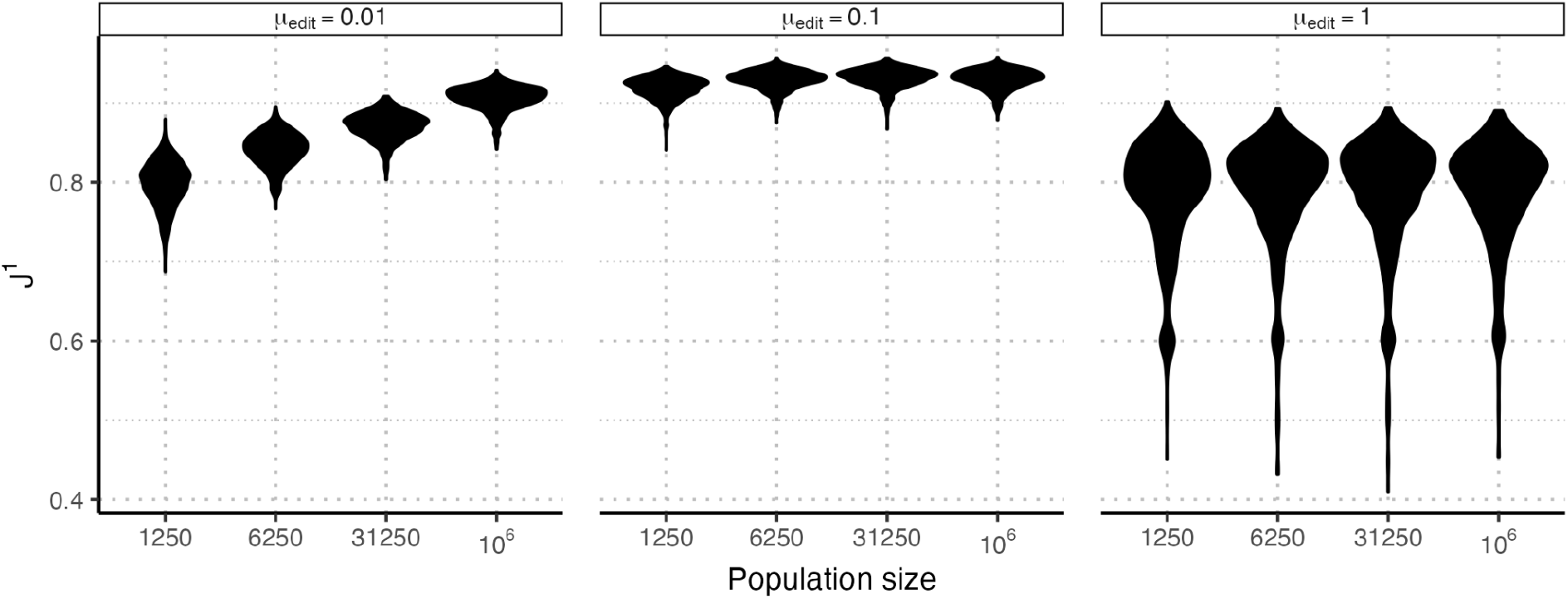
*J* ^1^ distributions for reconstructed EBR lineage trees. The distribution of *J* ^1^ in reconstructed EBR lineage tracing trees downsampled to 1250 tips from different population sizes (*n*_*tips*_ ϵ {1250, 6250, 31250, 10_6_}) and with different editing rates (µ_*edit*_ ϵ {0. 01, 0. 1, 1}, faceted panels). *J*^1^ isrobust to total population size at higher editing rates (µ_*edit*_ϵ {0. 1, 1}) but shows a population size dependence at lower editing rates (µ_*edit*_ = 0. 01), likely due to insufficient time for barcode saturation. Each distribution summarizes over 1000 replicates.

**Figure S7.**
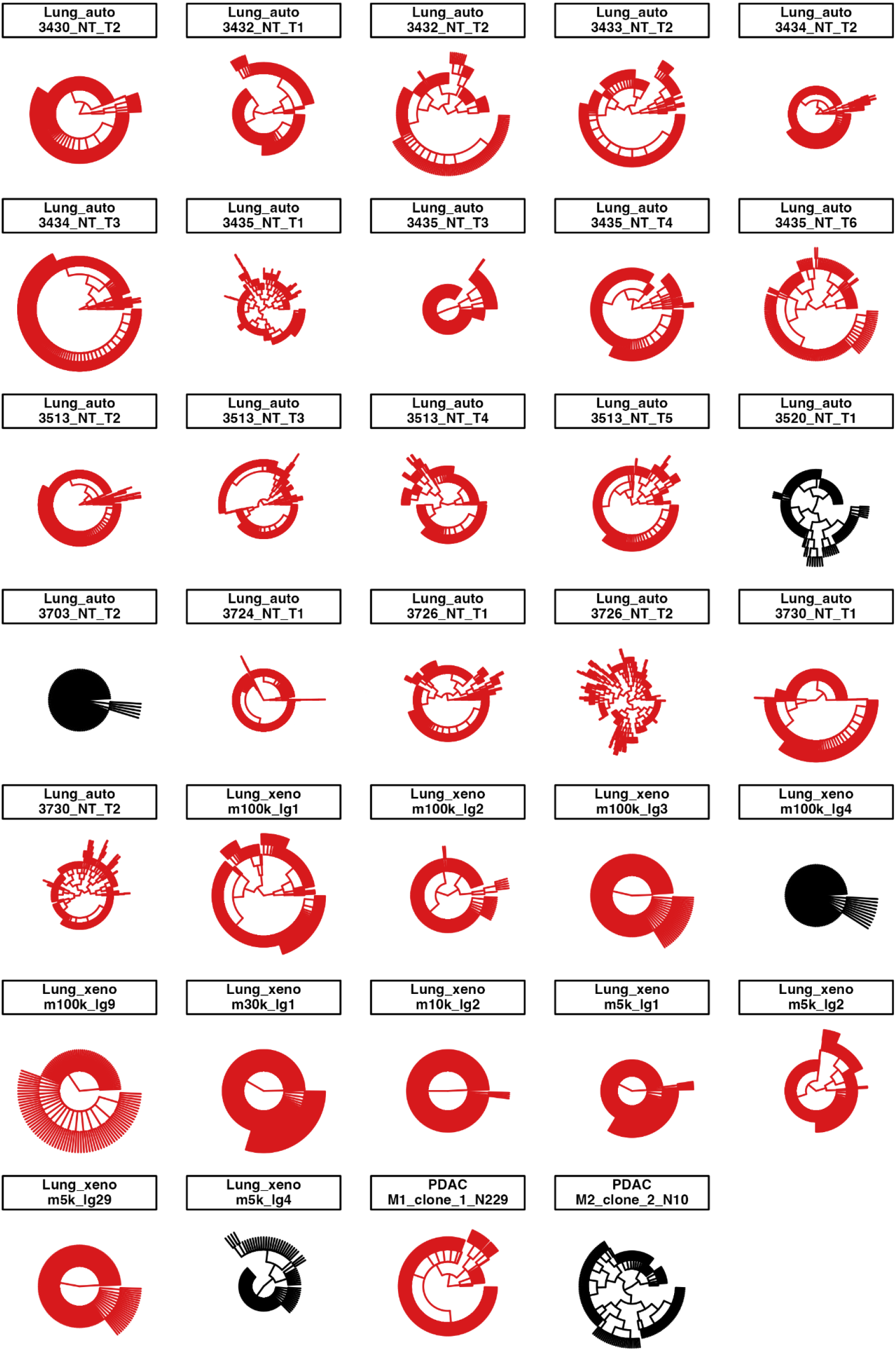
Branching rate heterogeneity is widespread in three *in vivo* lineage tracing datasets. We plot reconstructed tree topologies from three *in vivo* lineage tracing studies of cancer cell lines or autochthonous lung cancers. Trees with a significant departure from EBR are displayed in red, while those without are displayed in black. All trees are displayed in their full size.

## References

Alves JM, Prado-López S, Cameselle-Teijeiro JM, Posada D. 2019. Rapid evolution and biogeographic spread in a colorectal cancer. Nat. Commun. 10:5139. doi: 10.1038/s41467-019-12926-8.

Baslan T et al. 2020. Novel insights into breast cancer copy number genetic heterogeneity revealed by single-cell genome sequencing. eLife. 9:e51480. doi: 10.7554/eLife.51480.

Bian S et al. 2018. Single-cell multiomics sequencing and analyses of human colorectal cancer. Science. 362:1060–1063. doi: 10.1126/science.aao3791.

Borgsmüller N, Valecha M, Kuipers J, Beerenwinkel N, Posada D. 2023. Single-cell phylogenies reveal changes in the evolutionary rate within cancer and healthy tissues. Cell Genomics. 3:100380. doi: 10.1016/j.xgen.2023.100380.

Bozic I et al. 2010. Accumulation of driver and passenger mutations during tumor progression. Proc. Natl. Acad. Sci. 107:18545–18550. doi: 10.1073/pnas.1010978107.

Casasent AK et al. 2018. Multiclonal Invasion in Breast Tumors Identified by Topographic Single Cell Sequencing. Cell. 172:205-217.e12. doi: 10.1016/j.cell.2017.12.007.

Chen C, Liao Y, Peng G. 2022. Connecting past and present: single-cell lineage tracing. Protein Cell. 13:790–807. doi: 10.1007/s13238-022-00913-7.

Chkhaidze K et al. 2019. Spatially constrained tumour growth affects the patterns of clonal selection and neutral drift in cancer genomic data. PLOS Comput. Biol. 15:e1007243. doi: 10.1371/journal.pcbi.1007243.

Choi J et al. 2022. A time-resolved, multi-symbol molecular recorder via sequential genome editing. Nature. 608:98–107. doi: 10.1038/s41586-022-04922-8.

Eyler CE et al. 2020. Single-cell lineage analysis reveals genetic and epigenetic interplay in glioblastoma drug resistance. Genome Biol. 21:174. doi: 10.1186/s13059-020-02085-1.

Feng J et al. 2021. Estimation of cell lineage trees by maximum-likelihood phylogenetics. Ann. Appl. Stat. 15:343–362. doi: 10.1214/20-AOAS1400.

Fennell KA et al. 2022. Non-genetic determinants of malignant clonal fitness at single-cell resolution. Nature. 601:125–131. doi: 10.1038/s41586-021-04206-7.

FitzJohn RG. 2012. Diversitree: comparative phylogenetic analyses of diversification in R. Methods Ecol. Evol. 3:1084–1092. doi: 10.1111/j.2041-210X.2012.00234.x.

Gaglia G et al. 2022. Temporal and spatial topography of cell proliferation in cancer. Nat. Cell Biol. 24:316–326. doi: 10.1038/s41556-022-00860-9.

Gerlinger M et al. 2012. Intratumor Heterogeneity and Branched Evolution Revealed by Multiregion Sequencing. N. Engl. J. Med. 366:883–892. doi: 10.1056/NEJMoa1113205.

Griffiths RC, Tavaré S. 1998. The age of a mutation in a general coalescent tree. Commun. Stat. Stoch. Models. 14:273–295. doi: 10.1080/15326349808807471.

Harvey MG, Rabosky DL. 2018. Continuous traits and speciation rates: Alternatives to state-dependent diversification models. Methods Ecol. Evol. 9:984–993. doi: 10.1111/2041-210X.12949.

Househam J et al. 2022. Phenotypic plasticity and genetic control in colorectal cancer evolution. Nature. 611:744–753. doi: 10.1038/s41586-022-05311-x.

Jiang X, Tomlinson IPM. 2020. Why is cancer not more common? A changing microenvironment may help to explain why, and suggests strategies for anti-cancer therapy. Open Biol. 10:190297. doi: 10.1098/rsob.190297.

Jones MG et al. 2020. Inference of single-cell phylogenies from lineage tracing data using Cassiopeia. Genome Biol. 21:92. doi: 10.1186/s13059-020-02000-8.

Jones MG, Yang D, Weissman JS. 2023. New Tools for Lineage Tracing in Cancer In Vivo. Annu. Rev. Cancer Biol. 7:111–129. doi: 10.1146/annurev-cancerbio-061421-123301.

Kersting SJ, Wicke K, Fischer M. 2025. Tree balance in phylogenetic models. Philos. Trans. R. Soc. B Biol. Sci. 380:20230303. doi: 10.1098/rstb.2023.0303.

Kopperud BT, Magee AF, Höhna S. 2023. Rapidly changing speciation and extinction rates can be inferred in spite of nonidentifiability. Proc. Natl. Acad. Sci. 120:e2208851120. doi: 10.1073/pnas.2208851120.

Lemant J, Le Sueur C, Manojlović V, Noble R. 2022. Robust, Universal Tree Balance Indices. Syst. Biol. 71:1210–1224. doi: 10.1093/sysbio/syac027.

Lewinsohn MA, Bedford T, Müller NF, Feder AF. 2023. State-dependent evolutionary models reveal modes of solid tumour growth. Nat. Ecol. Evol. 7:581–596. doi: 10.1038/s41559-023-02000-4.

Lin D et al. 2023. Time-tagged ticker tapes for intracellular recordings. Nat. Biotechnol. 41:631–639. doi: 10.1038/s41587-022-01524-7.

Liu K et al. 2021. Mapping single-cell-resolution cell phylogeny reveals cell population dynamics during organ development. Nat. Methods. 18:1506–1514. doi: 10.1038/s41592-021-01325-x.

Liu X et al. 2024. Tumor phylogeography reveals block-shaped spatial heterogeneity and the mode of evolution in Hepatocellular Carcinoma. Nat. Commun. 15:3169. doi: 10.1038/s41467-024-47541-9.

Lynch AR, Arp NL, Zhou AS, Weaver BA, Burkard ME. 2022. Quantifying chromosomal instability from intratumoral karyotype diversity using agent-based modeling and Bayesian inference Marston, AL, Akhmanova, A, & Graham, TA, editors. eLife. 11:e69799. doi: 10.7554/eLife.69799.

Maddison WP, Midford PE, Otto SP. 2007. Estimating a Binary Character’s Effect on Speciation and Extinction. Syst. Biol. 56:701–710. doi: 10.1080/10635150701607033.

Martínez-Ruiz C et al. 2023. Genomic–transcriptomic evolution in lung cancer and metastasis. Nature. 616:543–552. doi: 10.1038/s41586-023-05706-4.

Neher RA, Hallatschek O. 2013. Genealogies of rapidly adapting populations. Proc. Natl. Acad. Sci. 110:437–442. doi: 10.1073/pnas.1213113110.

Neher RA, Russell CA, Shraiman BI. 2014. Predicting evolution from the shape of genealogical trees McVean, G, editor. eLife. 3:e03568. doi: 10.7554/eLife.03568.

Noble R et al. 2022. Spatial structure governs the mode of tumour evolution. Nat. Ecol. Evol. 6:207–217. doi: 10.1038/s41559-021-01615-9.

Pei W et al. 2017. Polylox barcoding reveals haematopoietic stem cell fates realized in vivo. Nature. 548:456–460. doi: 10.1038/nature23653.

Prillo S, Ravoor A, Yosef N, Song YS. 2023. ConvexML: Scalable and accurate inference of single-cell chronograms from CRISPR/Cas9 lineage tracing data. bioRxiv. 2023.12.03.569785. doi: 10.1101/2023.12.03.569785.

Quinn JJ et al. 2021. Single-cell lineages reveal the rates, routes, and drivers of metastasis in cancer xenografts. Science. 371:eabc1944. doi: 10.1126/science.abc1944.

Reeves MQ, Kandyba E, Harris S, Del Rosario R, Balmain A. 2018. Multicolour lineage tracing reveals clonal dynamics of squamous carcinoma evolution from initiation to metastasis. Nat. Cell Biol. 20:699–709. doi: 10.1038/s41556-018-0109-0.

Sackin MJ. 1972. “Good” and “Bad” Phenograms. Syst. Biol. 21:225–226. doi: 10.1093/sysbio/21.2.225.

Salehi S et al. 2023. Cancer phylogenetic tree inference at scale from 1000s of single cell genomes. Peer Community J. 3:e63. doi: 10.24072/pcjournal.292.

Schiffman JS et al. 2024. Defining heritability, plasticity, and transition dynamics of cellular phenotypes in somatic evolution. Nat. Genet. 56:2174–2184. doi: 10.1038/s41588-024-01920-6.

Schwartz R, Schäffer AA. 2017. The evolution of tumour phylogenetics: principles and practice. Nat. Rev. Genet. 18:213–229. doi: 10.1038/nrg.2016.170.

Schwarz RF et al. 2014. Phylogenetic Quantification of Intra-tumour Heterogeneity. PLOS Comput. Biol. 10:e1003535. doi: 10.1371/journal.pcbi.1003535.

Scott JG, Maini PK, Anderson ARA, Fletcher AG. 2020. Inferring Tumor Proliferative Organization from Phylogenetic Tree Measures in a Computational Model. Syst. Biol. 69:623–637. doi: 10.1093/sysbio/syz070.

Seidel S, Stadler T. 2022. TiDeTree: a Bayesian phylogenetic framework to estimate single-cell trees and population dynamic parameters from genetic lineage tracing data. Proc. R. Soc. B Biol. Sci. 289:20221844. doi: 10.1098/rspb.2022.1844.

Serio RN et al. 2024. Clonal Lineage Tracing with Somatic Delivery of Recordable Barcodes Reveals Migration Histories of Metastatic Prostate Cancer. Cancer Discov. 14:1990–2009. doi: 10.1158/2159-8290.CD-23-1332.

Sevillya G, Frenkel Z, Snir S. 2016. Triplet MaxCut: a new toolkit for rooted supertree Paradis, E, editor. Methods Ecol. Evol. 7:1359–1365. doi: 10.1111/2041-210X.12606.

Shao K-T, Sokal RR. 1990. Tree Balance. Syst. Biol. 39:266–276. doi: 10.2307/2992186.

Sheth RU, Yim SS, Wu FL, Wang HH. 2017. Multiplex recording of cellular events over time on CRISPR biological tape. Science. 358:1457–1461. doi: 10.1126/science.aao0958.

Simeonov KP et al. 2021. Single-cell lineage tracing of metastatic cancer reveals selection of hybrid EMT states. Cancer Cell. 39:1150-1162.e9. doi: 10.1016/j.ccell.2021.05.005.

Sokal RR. 1983. A Phylogenetic Analysis of the Caminalcules. I. the Data Base. Syst. Biol. 32:159–184. doi: 10.1093/sysbio/32.2.159.

Speidel L, Forest M, Shi S, Myers SR. 2019. A method for genome-wide genealogy estimation for thousands of samples. Nat. Genet. 51:1321–1329. doi: 10.1038/s41588-019-0484-x.

Werner B, Traulsen A, Sottoriva A, Dingli D. 2017. Detecting truly clonal alterations from multi-region profiling of tumours. Sci. Rep. 7:44991. doi: 10.1038/srep44991.

Williams MJ et al. 2018. Quantification of subclonal selection in cancer from bulk sequencing data. Nat. Genet. 50:895–903. doi: 10.1038/s41588-018-0128-6.

Yang D et al. 2022. Lineage tracing reveals the phylodynamics, plasticity, and paths of tumor evolution. Cell. 185:1905-1923.e25. doi: 10.1016/j.cell.2022.04.015.

Zhang W et al. 2021. The bone microenvironment invigorates metastatic seeds for further dissemination. Cell. 184:2471-2486.e20. doi: 10.1016/j.cell.2021.03.011.

Zheng X et al. 2020. Spatial Density and Distribution of Tumor-Associated Macrophages Predict Survival in Non–Small Cell Lung Carcinoma. Cancer Res. 80:4414–4425. doi: 10.1158/0008-5472.CAN-20-0069.

Zwaans A, Seidel S, Manceau M, Stadler T. 2025. A Bayesian phylodynamic inference framework for single-cell CRISPR/Cas9 lineage tracing barcode data with dependent target sites. Philos. Trans. R. Soc. B Biol. Sci. 380:20230318. doi: 10.1098/rstb.2023.0318.

