## Supplementary Data and Scripts for "Detecting branching rate heterogeneity with tree balance statistics in lineage tracing trees": README.txt

### README.txt
# This text file documents the list of files in this folder that supports the
# manuscript titled 'Detecting branching rate heterogeneity with tree balance
# statistics in lineage tracing trees'.
# Authors: Yingnan Gao, Alison F. Feder
# DOI: https://doi.org/10.1101/2024.06.27.601073

### Empirical_data/
# This folder holds the raw and processed data files from empirical studies.
# Directly under this folder, the following files can be found:

# Empirical_data/empirical_metadata.txt
# This text file documents some basic information about the empirical trees like
# name, dataset, tree size and barcode size.

# Empirical_data/empirical_CM_stat.RDS
# This binary file contains the editing proportion of character matrices in empirical
# datasets. It is the output from the script 'Scripts/Analysis_test_EBR_in_empirical_data.R'

# Empirical_data/empirical_res.RDS
# This binary file contains the outcome of the test against EBR in trees from empirical
# datasets. It is the output from the script 'Scripts/Analysis_test_EBR_in_empirical_data.R'
# and the binary version of Table S2.

# There are two subfolders as well:

# Empirical_data/Lung_auto/
# This subfolder holds the character matrices (*_character_matrix.txt) and the
# reconstructed lineage tracing trees by Cassiopeia's MaxCut solver in Newick
# format (*__MaxCut.tre) for a murine autochthonous lung cancer dataset.
# The character matrices were those published by Yang et al
# (https://zenodo.org/records/5847462).

# Empirical_data/Lung_xeno/
# This subfolder holds the character matrices (*_character_matrix.txt) and the
# reconstructed lineage tracing trees by Cassiopeia's MaxCut solver in Newick
# format (*__MaxCut.tre) for a murine xenograft lung cancer dataset.
# The character matrices were those published by Quinn et al
# (GEO accession no. GSE161363).


# Empirical_data/PDAC/
# This subfolder holds the character matrices (*_character_matrix.txt) and the
# reconstructed lineage tracing trees by Cassiopeia's MaxCut solver in Newick
# format (*__MaxCut.tre) for a murine PDAC dataset.
# The character matrices were derived from those published by Simeonov et al
# (https://data.mendeley.com/datasets/t98pjcd7t6/1).

### Figures_and_Tables/
# This folder holds the ouput figures and tables from scripts in Scripts/ folder
# The following files can be found here:

# Figures_and_Tables/Supplementary_Tables.xlsx
# This Excel file is the aggregated supplementary tables, manually generated from
# Table_S*.txt files

# Figures_and_Tables/Table_S1.txt
# This text file is the output from Scripts/Analysis_benchmarking_LT_estimation.R

# Figures_and_Tables/Table_S2.txt
# This text file is the output from Scripts/Analysis_test_EBR_in_empirical_data.R

# All other .pdf and .png files are figures as outputs from scripts in Scripts/
# folder whose name corresponds to the figures in the manuscript.

### Scripts/
# This folder holds the scripts that are essential to reproduce the simulation 
# and analyses in the manuscript. These files can be grouped to several categories:

# Scripts/Python/
# This subfolder holds the .py scripts for generating lineage tracing trees via
# Cassiopeia (https://github.com/YosefLab/Cassiopeia). See reproduce_simulation.sh
# for more details.

# Scripts/Python/infer_LT_tree_from_CM.py
# This script uses Cassiopeia to reconstruct the lineage tracing tree from a character
# matrix in a text file using the MaxCut algorithm.

# Scripts/Python/simulate_genealogical_trees.py
# This script uses Cassiopeia to simulate genealogical trees of certain sizes and
# a growth model, and outputs them as text Newick files.

# Scripts/Python/simulate_lineage_tracing_trees.py
# This script uses Cassiopeia to simulate lineage tracing trees from genealogical
# trees by simulating character matrices and reconstruct trees from them using the
# MaxCut algorithm.

# Scripts/Rscripts/
# This subfolder holds the .R scripts for tree generation and other functions to
# be run in batch (e.g., on a HPC platform). See Scripts/reproduce_simulation.sh
# for more details. The two scripts not used for simulating trees are:

# Scripts/Rscripts/calculate_editing_proportion.R
# This script extract the barcode saturation status from the simulated character
# matrices. The aggregated output from this script is stored in Simulated_data/stats

# Scripts/Rscripts/infer_for_each_CM_file.R
# This script generates the empirical trees using the MaxCut algorithm in the folders
# in Empirical_data/ from the character matrix files.

# Scripts/R/
# This subfolder holds the .R files to be sourced by other R scripts or in
# RStudio. See the comments in each file for more details. 

# Scripts/R/Jq_tree_balance.R
# This file contains the functions to calculate J^1 and the Sackin index.

# Scripts/R/phylo_ops.R
# This file contains utility functions such as the wrapper function to apply a 
# function across a phylogeny.

# Scripts/R/test_of_constant_rate.R
# This file contains the function to apply the statistical test against EBR 
# using J^1 or the Sackin index.

# Scripts/Analysis_*.R
# These files are scripts that reproduces the analyses in the manuscript. They can
# be sourced from RStudio with the working directory set to the same directory as
# this README.txt.

# Scripts/Fig_*.R
# These files are scripts that reproduce the figures in the manuscript.
# See Scripts/reproduce_figures_and_tables.sh for more details.

# Scripts/reproduce_figures_and_tables.sh
# This is a tutorial to reproduce the figures and tables in the manuscript.

# Scripts/reproduce_simulation.sh
# This is a tutorial to reproduce the trees in the manuscript.

### Simulated_data/
# This folder holds the simulated genealogical and lineage tracing trees in text
# Newick files and the statistics derived from them in binary RDS files.

# Simulated_data/test_trees/
# This subfolder holds the Newick files for simulated trees that are used as the
# focal trees in the analyses.
# The naming conventions are:
# [$model]_S[$rate_heterogeneity]_N[$population_size]_n[$tree_size].nwk
# for true genealogical trees and
# [$model]_S[$rate_heterogeneity]_N[$population_size]_n[$tree_size]._[$num_cassettes]_[$num_site_per_cassette]_[$editing_rate].tre
# for lineage tracing trees

# Simulated_data/null_trees/
# This subfolder holds the Newick files for simulated EBR trees that are used to
# generate the null distribution of J^1 and the Sackin index.
# The naming conventions are:
# NULL_N[$population_size]_n[$tree_size].nwk for true genealogical trees and
# NULL_N[$population_size]_n[$tree_size]._[$num_cassettes]_[$num_site_per_cassette]_[$editing_rate].tre
# for lineage tracing trees

# Simulated_data/stats/
# This subfolder holds the binary RDS files containing intermediate results of the
# analyses. In this subfolder, the following files can be found:

# Simulated_data/stats/J1_Sackin_nulls.RDS
# This file contains the null distribution of J^1 and the Sackin index derived from
# the trees in Simulated_data/null_trees/

# Simulated_data/stats/J1_Sackin_data.RDS
# This file contains the raw test results from Analysis_test_EBR_with_tree_balance_statistics.R

# Simulated_data/stats/J1_Sackin_power.RDS
# This file contains the power and type I error statistics from Analysis_test_EBR_with_tree_balance_statistics.R

# Simulated_data/stats/J1_robustness_data.RDS
# This file contains the raw test results from Analysis_robustness_of_J1.R

# Simulated_data/stats/J1_robustness_power.RDS
# This file contains the power and type I error statistics from Analysis_robustness_of_J1.R

# Simulated_data/stats/CM_stat_combined.RDS
# This file contains the barcode saturation status of trees in Simulated_data/null_trees/

# Simulated_data/stats/CM_stat_test.RDS
# This file contains the barcode saturation status of a subset of trees in the
# Simulated_data/test_trees/ subfolder.

# Simulated_data/stats/benchmark_LT_est_from_CM.RDS
# This file contains the results from Analysis_benchmarking_LT_estimation.R

# Simulated_data/stats/LT_topology_and_examples.RDS
# This file contains the results from Fig_2_Effect_of_lineage_tracing_on_tree_balance.R
