## Supplementary Data and Scripts for "Detecting branching rate heterogeneity with tree balance statistics in lineage tracing trees": Analysis_benchmarking_LT_estimation.R

```
# load libraries and read in data
library(parallel)
null.CM.stat = readRDS("Simulated_data/stats/CM_stat_combined.RDS")$N6250$k30
test.CM.stat = readRDS("Simulated_data/stats/CM_stat_test.RDS")
source("Scripts/Fig_Color_Palettes.R")

# compare the barcode saturation status between test and null samples
bc.saturation.KS.compare = lapply(test.CM.stat,function(x){
  true.LT = c(0.01,0.05,seq(0.1,1,0.1))
  names(true.LT) = paste0("LT",c(0.01,0.05,seq(0.1,1,0.1)))
  ksD = mclapply(x,function(xx){
    lapply(xx,function(xxx){
      lapply(null.CM.stat,function(this.null){
        this.D = sapply(this.null,function(yy){
          ks.test(xxx,yy)$statistic
        })
        min.D = min(this.D)
        median.D = median(this.D)
        median.D.idx = which.min(abs(median.D-this.D))
        return(list(min=min.D,median=median.D,repr=median.D.idx))
      })
    })
  },mc.cores = 4)
  return(list(D=ksD,LT=true.LT))
})

# estimate the editing rate from comparisons via KS test 
estimated.rates = lapply(bc.saturation.KS.compare,function(x){
  this.stat = lapply(names(x$D),function(nn){
    # raw D from KS.test
    this.est = sapply(x$D[[nn]],function(xx){
      sapply(xx,function(xxx){
        xxx$median
      })
    })
    # best LT
    this.est = apply(this.est,2,function(xx){
      x$LT[rownames(this.est)[which.min(xx)]]
    })
  })
  names(this.stat) = names(x$D)
  return(list(est=this.stat,LT=x$LT))
})

# check the accuracy of estimated lineage tracing rates
estimated.rates.accuracy = lapply(estimated.rates,function(x){
  mean.diff = sapply(names(x$est),function(nn){
    xx = x$est[[nn]]
    this.stat = c(
      bias=mean(xx-x$LT[[nn]])/x$LT[[nn]],
      error=mean(abs(xx-x$LT[[nn]]))/x$LT[[nn]],
      freq.over=sum(xx>x$LT[[nn]])/length(xx),
      freq.under=sum(xx<x$LT[[nn]])/length(xx))
    return(this.stat)
  })
  return(mean.diff)
})
saveRDS(list(raw=bc.saturation.KS.compare,stat=estimated.rates.accuracy),
        "Simulated_data/stats/benchmark_LT_est_from_CM.RDS")

#
tableS1.df = do.call(rbind,lapply(names(model.color),function(mdl){
  x = estimated.rates.accuracy[[mdl]]
  data.frame(model=mdl,
             edit_rate=sapply(colnames(x),function(xx){
               as.numeric(substr(xx,3,10))
             }),
             bias=x["bias",],error=x["error",],
             freq.over=x["freq.over",],freq.under=x["freq.under",])
}))
#
write.table(tableS1.df,"Figures_and_Tables/Table_S1.txt",sep = "\t",
            quote = FALSE,row.names = FALSE,col.names = TRUE)
```
