## Supplementary Data and Scripts for "Detecting branching rate heterogeneity with tree balance statistics in lineage tracing trees": Analysis_compile_benchmarking_null.R

```
# compile empirical null for benchmarking
#
library(ape)
library(parallel)
out = lapply(list.files("Scripts/R/","R",
                        full.names = TRUE),source)
rm(out)

# make metadata
null.table = rbind(expand.grid(N=c(1250,6250,31250,1e6),n=c(50,250,1250),type="TGT",edit_rate=0),
                   expand.grid(N=c(1250,6250,31250,1e6),n=c(50,250,1250),type="LTT",edit_rate=c(0.01,0.05,seq(0.1,1.0,0.1))))
null.table = null.table[null.table$N>=null.table$n,]
null.table$file = do.call(mapply,c(list(FUN=function(N,n,type,edit_rate){
  if(type=="TGT"){
    sprintf("Simulated_data/null_trees/NULL_N%d_n%d.nwk",N,n)
  }else{
    sprintf("Simulated_data/null_trees/NULL_N%d_n%d._3_10_%03d.tre",N,n,round(edit_rate*100))
  }
}),as.list(null.table[,c("N","n","type","edit_rate")])))

# calculate null distribution
nulls = lapply(null.table$file,function(f){
  print(f)
  trees = read.tree(f)
  trees = lapply(trees,collapse.singles)
  this.null = list(
    J1=sapply(trees,calculate_Jq_tree_balance_metric),
    Sackin=sapply(trees,calculate_Sackin_index))
})

saveRDS(list(meta=null.table[,c("N","n","type","edit_rate")],null=nulls),
        "Simulated_data/stats/J1_Sackin_nulls.RDS")
```
