## Supplementary Data and Scripts for "Detecting branching rate heterogeneity with tree balance statistics in lineage tracing trees": Analysis_test_EBR_in_empirical_data.R

```
# test EBR for empirical data
# load packages and source scripts
library(ape)
library(parallel)
out = lapply(list.files("Scripts/R/","R",
                        full.names = TRUE),source)
rm(out)

# read in trees
empirical.meta = read.table("Empirical_data/empirical_metadata.txt",sep = "\t",header = TRUE)
rownames(empirical.meta) = empirical.meta$tree_id
empirical.trees = mapply(FUN = function(tree_id,group){
  read.tree(sprintf("Empirical_data/%s/%s__MaxCut.tre",group,tree_id))
},empirical.meta$tree_id,empirical.meta$group,SIMPLIFY = FALSE)

# calculate editing proportion from CM for empirical trees
if(!file.exists("Empirical_data/empirical_CM_stat.RDS")){
  empirical.CM = mapply(FUN = function(tree_id,group){
    read.table(sprintf("Empirical_data/%s/%s_character_matrix.txt",group,tree_id),
               sep="\t",header=TRUE,row.names = 1)
  },empirical.meta$tree_id,empirical.meta$group,SIMPLIFY = FALSE)
  empirical.CM.stat = lapply(names(empirical.CM),function(nn){
    df = empirical.CM[[nn]]
    df.stat = do.call(rbind,apply(df,1,function(x){
      if(is.numeric(x)){
        n.miss = sum(x<0)
        n.none = sum(x==0)
      }else{
        n.miss = sum(x=="-")
        n.none = sum(x=="0")
      }
      return(data.frame(n.miss=n.miss,n.none=n.none))
    },simplify = FALSE))
    df.stat$n.all = empirical.meta[nn,"barcode_length"]
    df.stat$pp = (df.stat$n.all-df.stat$n.miss-df.stat$n.none)/(df.stat$n.all-df.stat$n.miss)
    return(df.stat)
  })
  saveRDS(empirical.CM.stat,"Empirical_data/empirical_CM_stat.RDS")
}else{
  empirical.CM.stat = readRDS("Empirical_data/empirical_CM_stat.RDS")
}
null.CM.stat = readRDS("Simulated_data/stats/CM_stat_combined.RDS")

# estimate LT rate for each empirical tree
est.edit.rate.raw = lapply(1:dim(empirical.meta)[1],function(i){
  if(empirical.meta[i,"group"]=="PDAC"){
    this.CM = null.CM.stat$N31250$k30
  }else{
    this.CM = null.CM.stat$N1000000$k30
  }
  ksD = lapply(this.CM,function(x){
    this.D = sapply(x,function(xx){
      ks.test(empirical.CM.stat[[i]]$pp,xx)$statistic
    })
    min.D = min(this.D)
    median.D = median(this.D)
    median.D.idx = which.min(abs(median.D-this.D))
    return(list(D=this.D,median=median.D,min=min.D,repr=median.D.idx))
  })
  return(ksD)
})
est.edit.rate = sapply(est.edit.rate.raw,function(x){
  this.median = sapply(x,function(xx){
    xx$median
    })
  est.LT = names(this.median)[which.min(this.median)]
  return(as.numeric(substr(est.LT,3,nchar(est.LT))))
  })
empirical.meta$estimated_LT = est.edit.rate

# test EBR for each empirical tree
set.seed(20250108)
empirical.test = do.call(rbind,mclapply(names(empirical.trees),function(i){
  if(empirical.meta[i,"group"]=="PDAC"){
    this.N = "31250"
  }else{
    this.N = "1000000"
  }
  this.n = empirical.meta[i,"tree_size"]
  this.phy = di2multi(collapse.singles(empirical.trees[[i]]))
  if(this.n>1250){
    this.phy = keep.tip(this.phy,sample(this.phy$tip.label,1250))
    this.n = 1250
  }
  n.cat = 5^ceiling(log(this.n/50)/log(5))*50
  this.rate = empirical.meta[i,"estimated_LT"]
  this.f = sprintf("Simulated_data/null_trees/NULL_N%s_n%d._3_10_%03d.tre",this.N,n.cat,round(this.rate*100))
  print(c(empirical.meta[i,"tree_id"],this.f))
  this.ref = lapply(read.tree(this.f),collapse.singles)
  if(this.n!=n.cat){
    this.ref = lapply(this.ref,function(phy){
      keep.tip(phy,sample(phy$tip.label,this.n))
    })
  }
  this.J1 = sapply(this.ref,calculate_Jq_tree_balance_metric)
  this.test = 
    test_of_constant_branching_rate_by_summary_statistic(
      phy = this.phy,func = calculate_Jq_tree_balance_metric,null.phy = this.J1)
  res = cbind(empirical.meta[i,],
              data.frame(J1=this.test$statistic,p.value=this.test$p.value))
  return(res)
},mc.cores = 4))
# save test results
saveRDS(empirical.test,"Empirical_data/empirical_res.RDS")
write.table(empirical.test,"Figures_and_Tables/Table_S2.txt",sep="\t",
            col.names = TRUE,row.names = FALSE,quote = FALSE)
```
