## Supplementary Data and Scripts for "Detecting branching rate heterogeneity with tree balance statistics in lineage tracing trees": Analysis_test_EBR_with_tree_balance_statistics.R

```
#
library(ape)
library(parallel)
library(stringr)
#
out = lapply(list.files("Scripts/R/",pattern = ".R",full.names = TRUE),source)
rm(out)

#
file.meta = data.frame(
  file=list.files("Simulated_data/test_trees/"))
file.meta$model = sapply(file.meta$file,substr,start=1,stop=3)
file.meta$type = sapply(file.meta$file,function(f){
  x=substr(f,start=nchar(f)-2,stop=nchar(f))
  return(c(nwk="TGT",tre="LTT")[x])
})
file.meta$LT = sapply(file.meta$file,function(f){
  x=as.numeric(str_extract(string = f,pattern = "(?<=_3_10_)[0-9]*"))/100
  if(is.na(x)) x = 0
  return(x)
})
file.meta$N = sapply(file.meta$file,function(f){
  as.numeric(str_extract(string = f,pattern = "(?<=N)[0-9]*"))
})
file.meta$n = sapply(file.meta$file,function(f){
  as.numeric(str_extract(string = f,pattern = "(?<=n)[0-9]*"))
})
file.meta$hetero = mapply(FUN = function(f,model){
  if(model=="EBR"){
    return(0)
  }else{
    c("01"=0.1,"05"=0.5,"1"=1,"5"=5,"10"=10,"062"=0.062,"126"=0.126,"312"=0.312,"624"=0.624)[
      str_extract(string = f,pattern = "(?<=S)[0-9]*")]
  }
},file.meta$file,file.meta$model)

#
empirical.nulls= readRDS("Simulated_data/stats/J1_Sackin_nulls.RDS")
empirical.nulls$key = do.call(
  mapply,c(list(FUN = function(...){paste0(sapply(list(...),as.character),collapse = "_")}),
           as.list(empirical.nulls$meta)))

#
J1_Sackin_res = do.call(rbind,mclapply(1:dim(file.meta)[1],function(i){
  print(i)
  phy = read.tree(paste0("Simulated_data/test_trees/",file.meta$file[i]))
  phy = lapply(phy,collapse.singles)
  this.idx = which(empirical.nulls$key==sprintf("%d_%d_%s_%s",file.meta$N[i],file.meta$n[i],file.meta$type[i],as.character(file.meta$LT[i])))
  res = do.call(rbind,lapply(phy,function(this.phy){
    J1 = test_of_constant_branching_rate_by_summary_statistic(
      phy = this.phy,null.phy = empirical.nulls$null[[this.idx]]$J1)
    Sackin = test_of_constant_branching_rate_by_summary_statistic(
      phy = this.phy,null.phy = empirical.nulls$null[[this.idx]]$Sackin,
      func = "calculate_Sackin_index",alternative = "greater")
    this.df = cbind(file.meta[i,-1],
                    data.frame(stat=c(J1$statistic,Sackin$statistic),
                               p.value=c(J1$p.value,Sackin$p.value),method=c("J1","Sackin")))
    return(this.df)
  }))
},mc.cores = 4))
saveRDS(J1_Sackin_res,"Simulated_data/stats/J1_Sackin_data.RDS")

J1_Sackin_stat = aggregate.data.frame(
  x = data.frame(power=J1_Sackin_res$p.value),
  by = J1_Sackin_res[,c("model","type","LT","N","n","hetero","method")],
  FUN = function(x){
    sum(x<=0.05,na.rm = TRUE)/length(x)
    })
saveRDS(J1_Sackin_stat,"Simulated_data/stats/J1_Sackin_power.RDS")
```
