## Supplementary Data and Scripts for "Detecting branching rate heterogeneity with tree balance statistics in lineage tracing trees": Fig_1_Signal_of_branching_rate_heterogeneity.R

```
# load packages
library(ape)
library(ggplot2)
library(ggtree)
library(ggpubr)
source("Scripts/Fig_Color_Palettes.R")

# read in example trees
file.table = data.frame(type="TGT",model=c("EBR","CRH","DRH"),edit_rate=0)
file.table$file = sapply(file.table$model,function(model){
  this.param = c(EBR="",CRH="_S1",DRH="_S624")[model]
  sprintf("Simulated_data/test_trees/%s%s_N6250_n50.nwk",model,this.param)
})
trees = lapply(file.table$file,function(f){
  collapse.singles(read.tree(f)[[1]])
})

# read in J1 and Sackin across models
J1_Sackin_res = readRDS("Simulated_data/stats/J1_Sackin_data.RDS")
J1_Sackin_stat = readRDS("Simulated_data/stats/J1_Sackin_power.RDS")

# pre-compute CI for all possible p^ in the experimental design
CI.data = do.call(rbind,lapply(0:1000,function(x){
  this.test = prop.test(x = x,n = 1000,p = 0.05)
  data.frame(phat=x/1000,pup=this.test$conf.int[2],plow=this.test$conf.int[1])
}))

# title, color, and adjustment settings
full.title = c(EBR="Equal branching\nrate",
               CRH="Continuous rate\nheterogeneity",
               DRH="Discrete rate\nheterogeneity")
method.power.adjust = c(J1=+0.004,Sackin=-0.004)

# tree shape plot
# ggtree objects for binary trees
binary.tree.plot = lapply(trees,function(this.phy){
  #remove branch lengths and normalize depth
  this.phy$edge.length = this.phy$edge.length*0+1
  maxT=max(node.depth.edgelength(this.phy))
  this.phy$edge.length = this.phy$edge.length/maxT
  # make ggtree object
  this.plot = 
    ggtree(tr = this.phy, layout = "fan",open.angle = 0)+
    theme_transparent()+
    theme(legend.position = "none",
          plot.margin = unit(c(0,0,0,0),units = "points"))
  })

# aggregate panel A compactly
fig1a = ggplot()+
  coord_cartesian(xlim=c(2.5,27.5),ylim = c(2.5,77.5),expand = FALSE)+
  annotation_custom(grob = cowplot::as_grob(binary.tree.plot[[1]]),
                    xmin = 0,xmax = 30,ymin = 50,ymax = 80)+
  annotation_custom(grob = cowplot::as_grob(binary.tree.plot[[2]]),
                    xmin = 0,xmax = 30,ymin = 25,ymax = 55)+
  annotation_custom(grob = cowplot::as_grob(binary.tree.plot[[3]]),
                    xmin = 0,xmax = 30,ymin = 0,ymax = 30)+
  theme_transparent()+
  theme(panel.background = element_blank(),axis.line = element_blank(),axis.ticks = element_blank(),
        axis.text = element_blank(),axis.title = element_blank(),
        legend.position = "none",plot.margin = unit(c(0,0,0,0),units = "points"))

# compile data for statistic distribution
fig1.data = J1_Sackin_res[J1_Sackin_res$n==1250&J1_Sackin_res$type=="TGT",]
fig1.data$group = paste0(fig1.data$model,"_",fig1.data$hetero)
fig1.data$hetero = factor(fig1.data$hetero,
                          levels = sort(unique(fig1.data$hetero)))
fig1.data$model = factor(fig1.data$model,levels = names(model.color))

# make sub-panels in panel B
fig1b.panels = lapply(c("J1","Sackin"),function(mm){
  this.data = do.call(rbind,lapply(c("CRH","DRH"),function(mdl){
    df= fig1.data[(fig1.data$method==mm)&(fig1.data$model%in%c(mdl,"EBR")),]
    df$facet_model = mdl
    return(df)
  }))
  this.plot = ggplot(data=this.data)+
    geom_violin(mapping=aes(x=hetero,y=stat,group=group,fill=model,color=model))+
    ggh4x::facet_grid2(rows = vars(facet_model), cols = vars(method),independent = "x",scales="free_x")+
    scale_color_manual(values=model.color,name="Growth\nmodel",drop=FALSE)+
    scale_fill_manual(values=model.color,name="Growth\nmodel",drop=FALSE)+
    xlab("Strength of rate\nheterogeneity")+ylab(parse(text=c(J1="J^1",Sackin="Sackin~~index")[mm]))+
    theme(legend.position = "bottom",
          axis.title = element_text(size=10),
          axis.text = element_text(size=8),
          axis.text.x = element_text(angle=30),
          legend.text = element_text(size = 8),
          legend.title = element_text(size=8,lineheight = 0.75),
          legend.key.height = unit(5,units="points"),
          legend.key.width = unit(5,units="points"),
          legend.title.position = "left",
          legend.background = element_rect(fill="transparent"),
          panel.background = element_blank(),
          panel.spacing.x = unit(4,units = "points"),
          strip.background = element_blank(),
          strip.text = element_blank(),
          panel.grid = element_line(color="grey",linetype = "dotted"),
          axis.line = element_line(),plot.margin = unit(c(4,8,4,8),units = "points"))
  return(this.plot)
})

# generate schematic for lots of EBR trees
box.df = data.frame(x=rev(seq(0,1,0.2)),y=rev(seq(0,1,0.2)))
null_tree.plot = ggplot(box.df)+
  geom_tile(mapping = aes(x=x,y=y),height=5,width=5,fill="white",color="black",linewidth=0.5)+
  annotation_custom(grob = cowplot::as_grob(binary.tree.plot[[1]]),xmin = -3,xmax = 3,ymin = -3,ymax = 3)+
  coord_cartesian(xlim = c(-2.6,3.6),ylim = c(-3,3.6))+
  theme(panel.background = element_blank(),axis.line = element_blank(),axis.ticks = element_blank(),
        axis.text = element_blank(),axis.title = element_blank(),
        legend.position = "none",plot.margin = unit(c(0,0,0,0),units = "points"))

# generate schematic for a focal tree
### This is a cherry-picked example with a p-value slightly below 0.05
example.tree = collapse.singles(read.tree("Simulated_data/test_trees/CRH_S1_N6250_n50.nwk")[[91]])
example.tree$edge.length = example.tree$edge.length*0+1
example.tree.plot = ggtree(tr = example.tree,open.angle = 0,layout = "fan")+
  theme_transparent()+
  theme(legend.position = "none",
        plot.margin = unit(c(0,0,0,0),units = "points"))
example.J1 = round(J1_Sackin_res$stat[106181]*2,2)/2
focal_tree.plot = ggplot()+
  annotate(geom = "tile",x=0,y=0,height=5,width=5,fill="white",color=model.color[2],linewidth=0.5)+
  annotation_custom(grob = cowplot::as_grob(example.tree.plot),xmin = -3.25,xmax = 3,ymin = -3,ymax = 3.25)+
  coord_cartesian(xlim = c(-2.6,3.6),ylim = c(-3,3.6))+
  theme(panel.background = element_blank(),axis.line = element_blank(),axis.ticks = element_blank(),
        axis.text = element_blank(),axis.title = element_blank(),
        legend.position = "none",plot.margin = unit(c(0,0,0,0),units = "points"))

# generate null distribution for J1
null_J1 = readRDS("Simulated_data/stats/J1_Sackin_nulls.RDS")$null[[2]]$J1
null.base = ggplot(data=data.frame(J1=round(null_J1*2,2)/2,outside=round(null_J1*2,2)/2<=example.J1))+
  geom_histogram(mapping = aes(x=J1,fill=outside),bins = 20,color="black")+
  geom_vline(xintercept = example.J1,color=model.color[2],linetype="dashed",linewidth=1)+
  scale_fill_manual(values=c("TRUE"="black","FALSE"="transparent"))+
  scale_x_continuous(breaks=c(0.6,0.7,0.8,0.9))+
  coord_cartesian(xlim=c(0.5,1),expand = FALSE)+
  xlab(" ")+
  theme(legend.position = "none",
        panel.background = element_blank(),
        panel.grid = element_blank(),
        axis.line.x = element_line(),
        axis.line.y = element_blank(),
        axis.ticks.y = element_blank(),
        axis.text.y = element_blank(),axis.text.x = element_text(size=8),
        axis.title.y = element_blank(),axis.title.x = element_text(size=10),
        plot.margin = unit(c(1,1,1,1),units = "points"))

# compile data for power and error
fig1.power.data = J1_Sackin_stat[J1_Sackin_stat$n==1250&J1_Sackin_stat$type=="TGT",]
fig1.power.data$upper = CI.data$pup[match(round(fig1.power.data$power,digits = 3),round(CI.data$phat,digits = 3))]
fig1.power.data$lower = CI.data$plow[match(round(fig1.power.data$power,digits = 3),round(CI.data$phat,digits = 3))]
fig1.power.data$power = fig1.power.data$power+method.power.adjust[fig1.power.data$method]
fig1.power.data$upper = fig1.power.data$upper+method.power.adjust[fig1.power.data$method]
fig1.power.data$lower = fig1.power.data$lower+method.power.adjust[fig1.power.data$method]
fig1.power.data = do.call(rbind,lapply(c("CRH","DRH"),function(mdl){
  df = fig1.power.data[fig1.power.data$model%in%c(mdl,"EBR"),]
  df$facet_model = mdl
  df$hetero = factor(df$hetero,levels=c(0,0.062,0.1,0.126,0.312,0.5,0.624,1,5,10))
  return(df)
}))

# make sub-panels for panel D
fig1d.panels = ggplot(fig1.power.data)+
  geom_line(mapping=aes(x=hetero,y=power,group=method,color=method))+
  geom_point(mapping=aes(x=hetero,y=power,group=method,color=method,shape=method))+
  geom_errorbar(mapping=aes(x=hetero,ymin=lower,ymax=upper,color=method,group=method),width=0.2)+
  geom_hline(yintercept=0.05,color="black",linetype="dashed")+
  ggh4x::facet_grid2(rows = vars(facet_model), cols = vars(type),independent = "x",scales="free_x")+
  scale_color_manual(name="Tree balance\nstatistics",values = tbs.color,labels=tbs.label)+
  scale_shape_discrete(name="Tree balance\nstatistics",labels=tbs.label)+
  coord_cartesian(ylim=c(0,1))+
  xlab("Strength of rate\nheterogeneity")+ylab("Power")+
  theme(legend.position = "bottom",
        axis.title = element_text(size=10),
        axis.text = element_text(size=8),
        axis.text.x = element_text(angle=30),
        legend.text = element_text(size = 8),
        legend.title = element_text(size=8,lineheight = 0.75),
        legend.title.position = "left",
        legend.background = element_rect(fill="transparent"),
        panel.background = element_blank(),
        panel.spacing.x = unit(4,units = "points"),
        strip.background = element_blank(),
        strip.text = element_blank(),
        panel.grid = element_line(color="grey",linetype = "dotted"),
        axis.line = element_line(),plot.margin = unit(c(4,8,4,8),units = "points"))

# align panel B and D
fig1bd = ggarrange(plotlist=c(fig1b.panels,list(fig1d.panels)),ncol=3,nrow=1,
                  align = "v",common.legend = TRUE,legend = "none")

# create the master canvas and add arrows and annotations
fig1 = ggplot()+
  coord_cartesian(xlim = c(-1,181),ylim = c(-1,131),expand = FALSE)+
  theme(panel.background = element_blank(),
        axis.line = element_blank(),axis.ticks = element_blank(),
        axis.text = element_blank(),axis.title = element_blank(),
        legend.position = "none",plot.margin = unit(c(0,0,0,0),units = "points"))
# add panels
fig1 = fig1+
  annotation_custom(grob = cowplot::as_grob(fig1bd),xmin = 35,xmax = 180,ymin = 0,ymax = 90)+
  annotation_custom(grob = cowplot::as_grob(fig1a),xmin = 2.5,xmax = 42.5,ymin = 12.5,ymax = 132.5)+
  annotation_custom(grob = cowplot::as_grob(focal_tree.plot),xmin = 55,xmax = 85,ymin = 100,ymax = 130)+
  annotation_custom(grob = cowplot::as_grob(null.base),xmin = 95,xmax = 135,ymin = 90,ymax = 125)+
  annotation_custom(grob = cowplot::as_grob(null_tree.plot),xmin = 150,xmax = 180,ymin = 97.5,ymax = 127.5)
# add growth model labels
fig1 = fig1+
  annotate(geom="tile", x = 2.75,y = 32.5,height=35,width=7,fill=model.color[3])+
  annotate(geom="tile", x = 2.75,y = 72.5,height=35,width=7,fill=model.color[2])+
  annotate(geom="tile", x = 2.75,y = 112.5,height=35,width=7,fill=model.color[1])+
  annotate(geom="text", x = 2.75,y = 32.5,label=full.title[3],angle=90,size=3,hjust=0.5,lineheight=0.75,color="white",fontface="bold")+
  annotate(geom="text", x = 2.75,y = 72.5,label=full.title[2],angle=90,size=3,hjust=0.5,lineheight=0.75,color="white",fontface="bold")+
  annotate(geom="text", x = 2.75,y = 112.5,label=full.title[1],angle=90,size=3,hjust=0.5,lineheight=0.75,color="white",fontface="bold")+
  annotate(geom="tile", x = 178,y = 34.5,height=30,width=4,fill=model.color[3])+
  annotate(geom="tile", x = 178,y = 73.5,height=30,width=4,fill=model.color[2])+
  annotate(geom="text", x = 178,y = 34.5,label=names(model.color)[3],angle=270,size=3,hjust=0.5,lineheight=0.75,color="white",fontface="bold")+
  annotate(geom="text", x = 178,y = 73.5,label=names(model.color)[2],angle=270,size=3,hjust=0.5,lineheight=0.75,color="white",fontface="bold")

# add panel C arrow annotations
arrows.df = data.frame(
  x=c(80,90,110,155,140),
  xend=c(90,111,90,140,130),
  y=c(125,117.5,102.5,125,108),
  yend=c(122.5,115,107.5,122.5,105),
  color=model.color[c("CRH","CRH","EBR","EBR","EBR")],
  curvature=c(-0.5,0.2,0.5,0.5,-0.5))
for(i in 1:dim(arrows.df)[1]){
  fig1 = fig1+
    annotate(geom="curve",x = arrows.df$x[i], xend = arrows.df$xend[i],y = arrows.df$y[i],
             yend = arrows.df$yend[i],color=arrows.df$color[i],curvature = arrows.df$curvature[i],
             linewidth=0.75,arrow = ggplot2::arrow(angle = 45,length = ggplot2::unit(4,units = "points")))
}
# add panel C text annotations
fig1 = fig1+
  annotate(geom="text", x = 69,y = 127.5,label="Focal tree",size=3,hjust=0.5,lineheight=0.75,color=model.color[2])+
  annotate(geom="text", x = 165,y = 128,label="EBR null tree",size=3,hjust=0.5,lineheight=0.75)+
  annotate(geom="text", x = 90,y = 120,label="J^1==0.705",parse=TRUE,size=3,hjust=0.5,lineheight=0.75,color=model.color[2])+
  annotate(geom="text", x = 140,y = 120,label="J^1==0.825",parse=TRUE,size=3,hjust=0.5,lineheight=0.75)+
  annotate(geom="text", x = 140,y = 117,label="J^1==0.845",parse=TRUE,size=3,hjust=0.5,lineheight=0.75)+
  annotate(geom="text", x = 140,y = 114,label="J^1==0.755",parse=TRUE,size=3,hjust=0.5,lineheight=0.75)+
  annotate(geom="text", x = 140,y = 111,label="...",size=3,hjust=0.5,lineheight=0.75)+
  annotate(geom="text", x = 90,y = 105,label="P==0.048",parse=TRUE,size=3,hjust=0.5,lineheight=0.75)

# add panel label
fig1 = fig1+
  annotate(geom="text", x = 9,y = 127.5,label="A",size=4,color="black",fontface="bold")+
  annotate(geom="text", x = 40,y = 90,label="B",size=4,color="black",fontface="bold")+
  annotate(geom="text", x = 55,y = 127.5,label="C",size=4,color="black",fontface="bold")+
  annotate(geom="text", x = 137,y = 90,label="D",size=4,color="black",fontface="bold")

# add panel legend
fig1 = fig1+
  annotation_custom(grob = cowplot::as_grob(get_legend(fig1b.panels[[1]])),xmin = 0,xmax = 50,ymin = 7,ymax = 13)+
  annotation_custom(grob = cowplot::as_grob(get_legend(fig1d.panels)),xmin = 1,xmax = 51,ymin = -1,ymax = 5)
# save to file
ggsave(plot = fig1,path = "Figures_and_Tables/",filename = "Figure_1.png",device = "png",
       height = 120,width = 185,scale = 1,units = "mm",dpi = 300)
ggsave(plot = fig1,path = "Figures_and_Tables/",filename = "Figure_1.pdf",device = "pdf",
       height = 120,width = 185,scale = 1,units = "mm",colormodel="cmyk")
```
