## Supplementary Data and Scripts for "Detecting branching rate heterogeneity with tree balance statistics in lineage tracing trees": Fig_2_Effect_of_lineage_tracing_on_tree_balance.R

```
# load packages
library(ape)
library(ggplot2)
library(ggtree)
library(ggpubr)
library(ggnewscale)
source("Scripts/R/phylo_ops.R")
source("Scripts/Fig_Color_Palettes.R")

# generate the metadata for trees to be used in this figure
file.table = rbind(
  expand.grid(type="TGT",model=c("EBR","CRH","DRH"),
              edit_rate=0,size=c(50,250,1250)),
  expand.grid(type="LTT",model=c("EBR","CRH","DRH"),
              edit_rate=c(0.01,0.05,0.10,0.50,1.00),size=c(50,250,1250))
)

# generate paths to each file
file.table$file = do.call(
  mapply,
  c(list(FUN=function(type,model,edit_rate,size){
    this.param = c(EBR="",CRH="_S1",DRH="_S624")[model]
    if(type=="TGT"){
      this.file = sprintf("Simulated_data/test_trees/%s%s_N6250_n%d.nwk",model,this.param,size)
    }else{
      this.file = sprintf("Simulated_data/test_trees/%s%s_N6250_n%d._3_10_%03d.tre",model,this.param,size,round(edit_rate*100))
    }
    return(this.file)
  }),
  as.list(file.table[,c("type","model","edit_rate","size")]))
)

# get example trees and tree statistics
### will take some time if generated from scratch
if(file.exists("Simulated_data/stats/LT_topology_and_examples.RDS")){
  tree.props=readRDS("Simulated_data/stats/LT_topology_and_examples.RDS")
}else{
  tree.props = lapply(1:dim(file.table)[1],function(j){
    f = file.table$file[j]
    x = read.tree(f)
    df = do.call(rbind,lapply(1:length(x),function(i){
      phy=collapse.singles(x[[i]])
      this.split = split(phy$edge[,2],phy$edge[,1])
      this.poly = do.call(rbind,lapply(names(this.split),function(nn){
        xx = this.split[[nn]]
        this.df = data.frame(model=file.table$model[j],edit_rate=file.table$edit_rate[j],
                             size=file.table$size[j],tree=i,m=length(xx),pendant=sum(xx<=Ntip(phy)))
        this.df$star = (this.df$m==this.df$pendant)&(this.df$m>2)
        this.df$starlike = !this.df$star&(this.df$pendant>=1)&(this.df$m>2)
        this.df$gpoly = (this.df$pendant<1)&(this.df$m>2)
        return(this.df)
      }))
      this.poly$type = do.call(what = mapply,args = c(list(FUN = function(...){
        paste0(as.numeric(c(...)),collapse="")
      },SIMPLIFY=TRUE),this.poly[,c("star","starlike","gpoly")]))
      return(this.poly)
    }))
    df = aggregate(1:dim(df)[1],by=df[,c("model","edit_rate","size","type")],FUN=length)
    df2 = do.call(rbind,lapply(1:length(x),function(i){
      phy = collapse.singles(x[[i]])
      root.clades = get_clade_size_for_each_node(phy)[phy$edge[phy$edge[,1]==(Ntip(phy)+1),2]]
      data.frame(model=file.table$model[j],edit_rate=file.table$edit_rate[j],size=file.table$size[j],
                 tree=i,node2tip=Nnode(phy)/Ntip(phy),root_balance=max(root.clades)/min(root.clades))
    }))
    example.tree=collapse.singles(x[[1]])
    return(list(example=example.tree,node=df,tree=df2))
  })
  saveRDS(tree.props,"Simulated_data/stats/LT_topology_and_examples.RDS")
}

# plot of simulated samples in the schematic
example.sample = ggtree(tr = tree.props[[1]]$example,layout = "fan",open.angle = 0)+
  theme_transparent()+
  theme(legend.position = "none",
        plot.margin = unit(c(0,0,0,0),units = "points"))
box.df = data.frame(x=rev(seq(0,1,0.2)),y=rev(seq(0,1,0.2)),model=rev(names(model.color)))
sample.plot = ggplot(box.df)+
  geom_tile(mapping = aes(x=x,y=y,color=model),height=5,width=5,fill="white",linewidth=0.5)+
  scale_color_manual(values=model.color)+
  annotation_custom(grob = cowplot::as_grob(example.sample),xmin = -2,xmax = 2,ymin = -2,ymax = 2)+
  coord_cartesian(xlim = c(-2.6,3.6),ylim = c(-3,3.6))+
  theme(panel.background = element_blank(),axis.line = element_blank(),axis.ticks = element_blank(),
        axis.text = element_blank(),axis.title = element_blank(),
        legend.position = "none",plot.margin = unit(c(0,0,0,0),units = "points"))

# example CM and their plots
set.seed(20250314)
example.CM = lapply(c(0.01,0.1,1),function(this.rate){
  res = phyloapply(x = list(barcode=numeric(9)),phy = tree.props[[1]]$example,
             func = function(x,l){
               wt = rexp(n = length(x[[1]]$barcode),rate = this.rate)
               mut = ceiling(runif(n=length(x[[1]]$barcode))*8)
               editable = x[[1]]$barcode==0
               x[[1]]$barcode[editable&(wt<l)] = mut[editable&(wt<l)]
               return(x[[1]])
             },
             use.branch.length = TRUE,use.self = FALSE,postorder = FALSE)
  res = res[as.vector(matrix(1:50,nrow = 10,ncol = 5,byrow = TRUE))]
  df = do.call(rbind,lapply(1:10,function(i){
    data.frame(raw=res[[i]]$barcode,state=res[[i]]$barcode,site=(1:9)+rep(c(0,0.25,0.5),each=3),tip=i)
    }))
  df$state = factor(df$state,0:8)
  return(df)
})

# generate CM plots
example.CM.plot = lapply(example.CM,function(df){
  ggplot(data = df)+
    geom_tile(mapping=aes(x=site,y=tip,fill=state),
              height=1,width=1,color="black",linewidth=0.5)+
    scale_fill_manual(values=CM.colors)+
    coord_cartesian(xlim = c(-1,11),ylim = c(-1,11),expand = FALSE)+
    theme(panel.background = element_blank(),axis.line = element_blank(),axis.ticks = element_blank(),
          axis.text = element_blank(),axis.title = element_blank(),
          legend.position = "none",plot.margin = unit(c(0,0,0,0),units = "points"))
})

# make a legend for the CM plots
legend.CM.plot = ggplot(
  data = data.frame(state=factor(0:8,levels=0:8),x=c(1,rep(1:4,2)),y=c(3.5,rep(1:2,each=4)))
  )+
  geom_tile(mapping=aes(x=x,y=y,fill=state),
            height=0.9,width=0.9,color="black",linewidth=0.5)+
  scale_fill_manual(values=CM.colors)+
  coord_cartesian(xlim = c(0,5),ylim = c(0,5),expand = FALSE)+
  theme(panel.background = element_blank(),axis.line = element_blank(),axis.ticks = element_blank(),
        axis.text = element_blank(),axis.title = element_blank(),
        legend.position = "none",plot.margin = unit(c(0,0,0,0),units = "points"))

# arrange panel A
fig2a = ggarrange(plotlist = c(list(sample.plot),example.CM.plot),
                  ncol = 4,nrow = 1,align = "hv",
                  legend = "none",common.legend = TRUE)

# tree shape plots
### these trees have transparent backgrounds so they can be placed on top of
### other plots
selected.trees = (file.table$size==50)&(file.table$edit_rate%in%c(0,0.01,0.1,1))
tree.plot = lapply(tree.props[selected.trees],function(x){
  this.phy = x$example
  #remove branch lengths and normalize depth
  this.phy$edge.length = this.phy$edge.length*0+1
  maxT=max(node.depth.edgelength(this.phy))
  this.phy$edge.length = this.phy$edge.length/maxT
  # make ggtree object
  this.split = split(this.phy$edge[,2],this.phy$edge[,1])
  this.poly = do.call(rbind,lapply(names(this.split),function(nn){
    xx = this.split[[nn]]
    this.df = data.frame(node=as.numeric(nn),m=length(xx),pendant=sum(xx<=Ntip(this.phy)))
    this.df$star = (this.df$m==this.df$pendant)&(this.df$m>2)
    this.df$starlike = !this.df$star&(this.df$pendant>=1)&(this.df$m>2)
    return(this.df)
  }))
  this.plot = 
    ggtree(tr = this.phy,mapping = aes(color=polytomy),layout = "fan",open.angle = 0)+
    scale_color_manual(name="Polytomy",values = tree.highlight.color,labels=tree.structure.label)+
    theme_transparent()+theme(legend.position = "none")
  this.plot$data$polytomy = "000"
  this.plot$data$polytomy[this.plot$data$parent%in%this.poly$node[this.poly$star]] = "100"
  this.plot$data$polytomy[this.plot$data$parent%in%this.poly$node[this.poly$starlike]] = "010"
  return(this.plot)
})
names(tree.plot) = paste0(file.table$model,"_",file.table$edit_rate)[selected.trees]

# arrange them in a matrix compactly (panel B)
fig2b.df = data.frame(
  xmin=rep(c(0,25,50,75),each=3),
  xmax=rep(c(30,55,80,105),each=3),
  ymin=rep(c(50,25,0),4),
  ymax=rep(c(80,55,30),4),row.names = names(tree.plot))
fig2b = ggplot()+
  coord_cartesian(xlim=c(2.5,102.5),ylim = c(2.5,77.5),expand = FALSE)+
  theme(panel.background = element_blank(),axis.line = element_blank(),axis.ticks = element_blank(),
        axis.text = element_blank(),axis.title = element_blank(),
        legend.position = "none",plot.margin = unit(c(0,0,0,0),units = "points"))
for(nn in names(tree.plot)){
  fig2b = fig2b+annotation_custom(
    grob = cowplot::as_grob(tree.plot[[nn]]),
    xmin = fig2b.df[nn,"xmin"],xmax = fig2b.df[nn,"xmax"],
    ymin = fig2b.df[nn,"ymin"],ymax = fig2b.df[nn,"ymax"])
}
rm(nn)

# generate the panels for polytomy compositions
polytomy.counts = do.call(rbind,lapply(tree.props,function(x){
  x$node$freq = x$node$x/sum(x$node$x)
  return(x$node)
}))
polytomy.counts$edit_rate = factor(polytomy.counts$edit_rate,c(0,0.01,0.05,0.10,0.50,1.00))

# generate panel C in the main figure
fig2c = ggplot(polytomy.counts[(polytomy.counts$edit_rate!=0)&(polytomy.counts$size==1250),])+
  geom_bar(mapping=aes(x=edit_rate,y=freq,fill=type,alpha=type),stat = "identity")+
  facet_wrap(facets = vars(model), nrow = 3,ncol = 1,strip.position = "right")+
  #scale_y_continuous(position = "right")+
  scale_fill_manual(name="   ",values = tree.structure.color,labels=tree.structure.label)+
  scale_alpha_manual(name="   ",values = tree.structure.alpha,labels=tree.structure.label)+
  xlab("Lineage tracing editing rate")+ylab("Frequency")+
  coord_cartesian(ylim = c(0,0.5),expand = TRUE)+
  guides(fill=guide_legend(nrow = 2,ncol = 2,reverse = TRUE,direction = "vertical"),
         alpha=guide_legend(nrow = 2,ncol = 2,reverse = TRUE),direction = "vertical")+
  theme(legend.position = "top",
        legend.key.size = unit(c(8,8),units="points"),
        legend.text = element_text(size = 8),
        legend.title = element_text(size=10),
        strip.background = element_blank(),
        strip.text = element_blank(),
        panel.background = element_blank(),
        panel.grid = element_line(color="grey",linetype = "dotted"),
        axis.title = element_text(size=10),
        axis.text = element_text(size=8),
        axis.line = element_line(),plot.margin = unit(c(10,10,10,10),units = "points"))

# generate node-to-tip ratio and root balance data for panel D and E
fig2de.data = do.call(rbind,lapply(tree.props[file.table$size==1250],function(x){
  xx = x$tree
  data.frame(model=xx$model[1],edit_rate=xx$edit_rate[1],
             n2t.mean = mean(xx$node2tip),
             n2t.upper=quantile(xx$node2tip,probs=0.75),
             n2t.lower=quantile(xx$node2tip,probs=0.25),
             rtb.mean = mean(log(xx$root_balance)),
             rtb.upper=quantile(log(xx$root_balance),probs=0.75),
             rtb.lower=quantile(log(xx$root_balance),probs=0.25))
}))
fig2de.data$edit_rate = factor(fig2de.data$edit_rate,levels = c(0,0.01,0.05,0.1,0.5,1))

# generate panel D of node-to-tip ratio
fig2d = ggplot(data=fig2de.data)+
  geom_point(mapping=aes(x=edit_rate,y=n2t.mean,color=model,group=model))+
  geom_errorbar(mapping=aes(x=edit_rate,y=n2t.mean,ymin=n2t.lower,ymax=n2t.upper,color=model),
                position = position_dodge(width = 0.1))+
  geom_line(data=fig2de.data[fig2de.data$edit_rate!=0,],
            mapping=aes(x=edit_rate,y=n2t.mean,color=model,group=model),linewidth=1)+
  scale_color_manual(name="Model",values = model.color,labels=names(model.color))+
  scale_x_discrete(breaks=c(0,0.01,0.05,0.10,0.50,1.00),
                   labels=c("True\ntree","0.01","0.05","0.1","0.5","1"))+
  xlab("Lineage tracing editing rate")+ylab("Node-to-tip ratio")+
  coord_cartesian(ylim = c(0,1),expand = TRUE)+
  theme(legend.position = "none",
        legend.key.size = unit(c(8,8),units="points"),
        legend.text = element_text(size = 8),
        legend.title = element_text(size=10),
        strip.background = element_blank(),
        strip.text = element_blank(),
        panel.background = element_blank(),
        panel.grid = element_line(color="grey",linetype = "dotted"),
        axis.title.y = element_text(size=10),
        axis.title.x = element_blank(),
        axis.text = element_text(size=8),
        axis.line = element_line(),plot.margin = unit(c(10,10,10,10),units = "points"))

# generate panel E of root balance
fig2e = ggplot(fig2de.data)+
  geom_point(mapping=aes(x=edit_rate,y=rtb.mean,color=model,group=model))+
  geom_errorbar(mapping=aes(x=edit_rate,y=rtb.mean,ymin=rtb.lower,ymax=rtb.upper,color=model),
                position = position_dodge(width = 0.1))+
  geom_line(data=fig2de.data[fig2de.data$edit_rate!=0,],
            mapping=aes(x=edit_rate,y=rtb.mean,color=model,group=model),linewidth=1)+
  scale_color_manual(name="Model",values = model.color,labels=names(model.color))+
  scale_x_discrete(breaks=c(0,0.01,0.05,0.10,0.50,1.00),
                   labels=c("True\ntree","0.01","0.05","0.1","0.5","1"))+
  xlab("Lineage tracing editing rate")+ylab("Sister size ratio")+
  #coord_cartesian(ylim = c(0,4),expand = TRUE)+
  theme(legend.position = "none",
        legend.key.size = unit(c(8,8),units="points"),
        legend.text = element_text(size = 8),
        legend.title = element_text(size=10),
        strip.background = element_blank(),
        strip.text = element_blank(),
        panel.background = element_blank(),
        panel.grid = element_line(color="grey",linetype = "dotted"),
        axis.title = element_text(size=10),
        axis.text = element_text(size=8),
        axis.line = element_line(),plot.margin = unit(c(10,10,10,10),units = "points"))
# combine panel D and E for alignment issues
fig2de = ggarrange(plotlist=list(fig2d,NULL,fig2e),ncol = 1,nrow = 3,align = "v",
                   heights = c(0.975,-0.075,1))

# read in J1 and Sackin across models for panel F
J1_Sackin_res = readRDS("Simulated_data/stats/J1_Sackin_data.RDS")

# organize data table, generate pseudo-Sackin for plotting
fig2f.data = J1_Sackin_res[J1_Sackin_res$n==1250&J1_Sackin_res$hetero%in%c(0,0.624,1),]
fig2f.data = split(fig2f.data,fig2f.data$model)
fig2f.data = do.call(rbind,lapply(c("CRH","DRH"),function(mdl){
  this.data = do.call(rbind,fig2f.data[c(mdl,"EBR")])
  this.data$facet_model=mdl
  return(this.data)
}))
fig2f.data$model = factor(fig2f.data$model,levels = names(model.color))
fig2f.data$LT = factor(fig2f.data$LT,levels = sort(unique(fig2f.data$LT)))
Sackin.range = range(fig2f.data$stat[fig2f.data$method=="Sackin"])
J1.range = range(fig2f.data$stat[fig2f.data$method=="J1"])
fig2f.data$pseudo = mapply(FUN = function(stat,method){
  if(method=="Sackin"){
    stat = (stat-Sackin.range[1])/(Sackin.range[2]-Sackin.range[1])*
      (J1.range[2]-J1.range[1])+J1.range[1]
  }
  return(stat)
},fig2f.data$stat,fig2f.data$method,SIMPLIFY = TRUE)
pseudo.Sackin.breaks = sapply(c(10000,20000,30000,40000),function(stat){
  (stat-Sackin.range[1])/(Sackin.range[2]-Sackin.range[1])*
    (J1.range[2]-J1.range[1])+J1.range[1]
})
pseudo.Sackin.labels = sapply(1:4,function(x){parse(text=sprintf("%d%s",x,"%*%10^4"))})
fig2f.data$group = paste0(fig2f.data$model,"_",fig2f.data$LT)

# generate panel F
fig2f= ggplot(data=fig2f.data)+
  geom_violin(mapping=aes(x=LT,y=stat,group=group,fill=model,color=model))+
  scale_color_manual(values=model.color,name="Model",drop=FALSE)+
  scale_fill_manual(values=model.color,name="Model",drop=FALSE)+
  ggh4x::facet_grid2(rows=vars(facet_model),cols = vars(method),labeller = label_parsed,
                     independent = "y",scales = "free_y",axes = "all")+
  scale_x_discrete(breaks=c(0,0.01,0.05,0.10,0.50,1.00),
                   labels=c("True\ntree","0.01","0.05","0.1","0.5","1"))+
  scale_y_continuous(breaks = c(seq(0,1,0.2),1e4*(1:4)),
                     labels = c(sapply(seq(0,1,0.2),function(x){
                       parse(text=sprintf("%.1f",x))
                     }),pseudo.Sackin.labels))+
  xlab("Lineage tracing editing rate")+ylab(parse(text="J^1"))+
  theme(legend.position = "none",
        axis.title = element_text(size=10),
        axis.text = element_text(size=8),
        legend.text = element_text(size = 8),
        legend.title = element_text(size=10),
        panel.background = element_blank(),
        panel.spacing.x = unit(30,units = "points"),
        strip.background = element_blank(),
        strip.text = element_blank(),
        panel.grid = element_line(color="grey",linetype = "dotted"),
        axis.line = element_line(),plot.margin = unit(c(10,10,10,10),units = "points"))

# create the master canvas and add arrows and annotations
fig2 = ggplot()+
  coord_cartesian(xlim = c(-1,181),ylim = c(-1,235),expand = FALSE)+
  theme(panel.background = element_blank(),
        axis.line = element_blank(),axis.ticks = element_blank(),
        axis.text = element_blank(),axis.title = element_blank(),
        legend.position = "none",plot.margin = unit(c(0,0,0,0),units = "points"))
# add panel A to F
fig2 = fig2+
  # smallest sub-panel within each panel roughly size 30 by 30
  # use this standard to unify font sizes: 8 pt sub-panel ~ 3 pt in main canvas
  annotation_custom(grob = cowplot::as_grob(fig2a),xmin = 5,xmax = 125,ymin = 195,ymax = 225)+
  annotation_custom(grob = cowplot::as_grob(fig2b),xmin = 5,xmax = 125,ymin = 100,ymax = 190)+
  annotation_custom(grob = cowplot::as_grob(fig2c),xmin = 125,xmax = 180,ymin = 90,ymax = 210)+
  annotation_custom(grob = cowplot::as_grob(fig2de),xmin = 0,xmax = 62.5,ymin = 0,ymax = 90)+
  annotation_custom(grob = cowplot::as_grob(fig2f),xmin = 62.5,xmax = 180,ymin = 0,ymax = 90)

# add missing Y-axis label in panel F
fig2 = fig2+
  annotate(geom = "text",x = 124.5,y = 50,label="Sackin index",
           size=3.5,hjust=0.5,angle=90,lineheight=0.8)

# add panel labels 
fig2 = fig2+
  annotate(geom = "text", label ="A",x = 2.5,y = 232.5,size=4,fontface="bold")+
  annotate(geom = "text", label ="B",x = 2.5,y = 190,size=4,fontface="bold")+
  annotate(geom = "text", label ="C",x = 127.5,y = 187.5,size=4,fontface="bold")+
  annotate(geom = "text", label ="D",x = 2.5,y = 87.5,size=4,fontface="bold")+
  annotate(geom = "text", label ="E",x = 2.5,y = 45,size=4,fontface="bold")+
  annotate(geom = "text", label ="F",x = 65,y = 87.5,size=4,fontface="bold")

# add legend annotations
fig2 = fig2+
  annotation_custom(grob = cowplot::as_grob(legend.CM.plot),
                    xmin = 135,xmax = 150,ymin = 202,ymax = 217)+
  annotate(geom="tile", x = 137,y = 221,height=2,width=2,fill=model.color[1])+
  annotate(geom="tile", x = 152,y = 221,height=2,width=2,fill=model.color[2])+
  annotate(geom="tile", x = 167,y = 221,height=2,width=2,fill=model.color[3])+
  annotate(geom = "text",x = 135,y = 230,label="Legends",size=4,hjust=0,fontface="bold")+
  annotate(geom = "text",x = 135,y = 225,label="Growth model (all panels)",size=3,hjust=0)+
  annotate(geom = "text",x = 139,y = 221,label="EBR",size=3,hjust=0)+
  annotate(geom = "text",x = 154,y = 221,label="CRH",size=3,hjust=0)+
  annotate(geom = "text",x = 169,y = 221,label="DRH",size=3,hjust=0)+
  annotate(geom = "text",x = 135,y = 217,label="Editing states (Panel A)",size=3,hjust=0)+
  annotate(geom = "text",x = 135,y = 202,label="Polytomy (Panel B and C)",size=3,hjust=0)+
  annotate(geom = "text",x = 150,y = 213,label="Unedited",size=3,hjust=0)+
  annotate(geom = "text",x = 150,y = 207.25,label="Edited",size=3,hjust=0)

# add Panel A and B texts
headers=c("True~~genealogical~~trees","Character~~matrices", "Reconstructed~~lineage~~tracing~~trees",
          "mu[edit]==0.01","mu[edit]==0.10","mu[edit]==1.00")
full.title = c(EBR="Equal branching\nrate",
               CRH="Continuous rate\nheterogeneity",
               DRH="Discrete rate\nheterogeneity")
fig2 = fig2+
  annotate(geom = "text",x = 17,y = 225,label="Cas-9 simulator",size=3,hjust=0,angle=19.3)+
  annotate(geom = "text",x = 80,y = 232.75,label=headers[2],size=3,hjust=0.5,parse=TRUE)+
  annotate(geom = "text",x = 52.5,y = 227.5,label=headers[4],size=4,hjust=0,parse=TRUE)+
  annotate(geom = "text",x = 82.5,y = 227.5,label=headers[5],size=4,hjust=0,parse=TRUE)+
  annotate(geom = "text",x = 112.5,y = 227.5,label=headers[6],size=4,hjust=0,parse=TRUE)+
  annotate(geom = "text",x = 81,y = 193.5,label="Reconstruct by\nMaxCut solver",size=3,hjust=0,lineheight=0.75)+
  annotate(geom = "text",x = 21,y = 193.5,label="Extract\ntopology",size=3,hjust=0,lineheight=0.75)+
  annotate(geom = "text",x = 38,y = 217,label="Cells",angle=90,size=3,hjust=0)+
  annotate(geom = "text",x = 40,y = 225.5,label="Sites",size=3,hjust=0)+
  annotate(geom = "text",x = 20,y = 100,label=headers[1],size=3,hjust=0.5,parse=TRUE)+
  annotate(geom = "text",x = 80,y = 100,label=headers[3],size=3,hjust=0.5,parse=TRUE)+
  annotate(geom="tile", x = 3.5,y = 115,height=25,width=7.5,fill=model.color[3])+
  annotate(geom="tile", x = 3.5,y = 145,height=25,width=7.5,fill=model.color[2])+
  annotate(geom="tile", x = 3.5,y = 175,height=25,width=7.5,fill=model.color[1])+
  annotate(geom="text", x = 3.5,y = 115,label=full.title[3],angle=90,size=3,hjust=0.5,lineheight=0.75,color="white",fontface="bold")+
  annotate(geom="text", x = 3.5,y = 145,label=full.title[2],angle=90,size=3,hjust=0.5,lineheight=0.75,color="white",fontface="bold")+
  annotate(geom="text", x = 3.5,y = 175,label=full.title[1],angle=90,size=3,hjust=0.5,lineheight=0.75,color="white",fontface="bold")+
  annotate(geom="rect", xmin = 176,xmax=180,ymin = 104,ymax=129,fill=model.color[3])+
  annotate(geom="rect", xmin = 176,xmax=180,ymin = 133,ymax=158,fill=model.color[2])+
  annotate(geom="rect", xmin = 176,xmax=180,ymin = 162,ymax=187,fill=model.color[1])+
  annotate(geom="text", x = 178,y = 116.5,label="DRH",angle=270,size=3,hjust=0.5,lineheight=0.75,color="white",fontface="bold")+
  annotate(geom="text", x = 178,y = 145.5,label="CRH",angle=270,size=3,hjust=0.5,lineheight=0.75,color="white",fontface="bold")+
  annotate(geom="text", x = 178,y = 174.5,label="EBR",angle=270,size=3,hjust=0.5,lineheight=0.75,color="white",fontface="bold")
# add arrows
# set up direction arrows and other annotations
arrows.df = data.frame(
  x=c(20,50,80,110,50,80,110),
  xend=c(20,50,80,110,50,80,110),
  y=c(198,198,198,198,231,231,231),
  yend=c(188,188,188,188,225,225,225))
segment.df = data.frame(
  x=c(20,40),
  xend=c(40,110),
  y=c(224,231),
  yend=c(231,231))

fig2 = fig2+
  geom_segment(data=arrows.df,mapping = aes(x=x,xend=xend,y=y,yend=yend),linewidth=0.75,
               arrow = ggplot2::arrow(angle = 45,length = ggplot2::unit(6,units = "points")))+
  geom_segment(data=segment.df,mapping = aes(x=x,xend=xend,y=y,yend=yend),linewidth=0.75)

# save to file
# for online preview
ggsave(plot = fig2,path = "Figures_and_Tables/",filename = "Figure_2.png",device = "png",
       height = 234,width = 185,scale = 1,units = "mm",dpi = 300)
# for submission
ggsave(plot = fig2,path = "Figures_and_Tables/",filename = "Figure_2.pdf",device = "pdf",
       height = 234,width = 185,scale = 1,units = "mm",colormodel = "cmyk")
#
```
