## Supplementary Data and Scripts for "Detecting branching rate heterogeneity with tree balance statistics in lineage tracing trees": Fig_3_S4_Performance_under_LT.R

```
#
library(ggplot2)
library(ggnewscale)
library(ggpubr)
source("Scripts/Fig_Color_Palettes.R")

#
full.data = readRDS("Simulated_data/stats/J1_Sackin_power.RDS")

# pre-compute CI for all possible p^ in the experimental design
CI.data = do.call(rbind,lapply(0:1000,function(x){
  this.test = prop.test(x = x,n = 1000,p = 0.05)
  data.frame(phat=x/1000,pup=this.test$conf.int[2],plow=this.test$conf.int[1])
}))

#
full.data$method = factor(full.data$method,levels = c("J1","Sackin"))
full.data$upper = CI.data$pup[match(full.data$power,CI.data$phat)]
full.data$lower = CI.data$plow[match(full.data$power,CI.data$phat)]
full.data$method_label = factor(c("J^1","\"Sackin index\"")[as.numeric(full.data$method)],
                                levels = c("J^1","\"Sackin index\""))
full.data$LT = factor(full.data$LT,sort(unique(full.data$LT)))
full.data$hetero = factor(full.data$hetero,levels=c(0,0.062,0.1,0.126,0.312,0.5,0.624,1,5,10))
plot.data = split(full.data,full.data$model)

#
fig3a.panels = lapply(c("CRH","DRH"),function(mdl){
  selected.EBR = (plot.data$EBR$n==1250)&(plot.data$EBR$LT==0.1|plot.data$EBR$type=="TGT")
  selected.mdl = (plot.data[[mdl]]$n==1250)&(plot.data[[mdl]]$LT==0.1|plot.data[[mdl]]$type=="TGT")
  this.data = rbind(plot.data[[mdl]][selected.mdl,],plot.data$EBR[selected.EBR,])
  this.panel = ggplot(data=this.data)+
    geom_point(mapping=aes(x=hetero,y=power,color=type,shape=type))+
    geom_line(mapping=aes(x=hetero,y=power,color=type,group=type))+
    geom_errorbar(mapping=aes(x=hetero,ymin=lower,ymax=upper,color=type,group=type),width=0.2)+
    geom_hline(yintercept = 0.05,color="black",linetype="dashed")+
    scale_color_manual(values = tree.type.colors,labels=tree.type.labels)+
    scale_shape_discrete(labels=tree.type.labels)+
    coord_cartesian(ylim=c(0,1))+
    xlab("Strength of rate heterogeneity")+ylab("Power")+
    facet_wrap(facet= vars(method_label),nrow = 1,labeller = label_parsed)+
    guides(color=guide_legend(title = "Tree type"),shape=guide_legend(title="Tree type"))+
    theme(legend.position = "none",
          strip.background = element_rect(fill = model.color[mdl]),
          strip.text = element_text(color="white",face="bold",size=8,margin=unit(c(0,0,0,0),units = "points")),
          panel.background = element_blank(),
          panel.grid = element_line(color="grey",linetype = "dotted"),
          axis.line = element_line(),plot.margin=unit(c(1,1,1,1),units = "points"),
          axis.title = element_text(size=10),
          axis.text = element_text(size=8),
          axis.text.x = element_text(angle=30))
  if(mdl=="CRH") this.panel = this.panel+theme(axis.title.x = element_blank())
  return(this.panel)
})

fig3b.panels = lapply(c("CRH","DRH"),function(mdl){
  selected.EBR = (plot.data$EBR$n==1250)&(plot.data$EBR$hetero==1)
  selected.mdl = (plot.data[[mdl]]$n==1250)&(plot.data[[mdl]]$hetero%in%c(0.624,1))
  this.data = rbind(plot.data[[mdl]][selected.mdl,],plot.data$EBR[selected.EBR,])
  this.plot = ggplot(data = this.data)+
    geom_point(mapping=aes(x=LT,y=power,color=type,shape=type))+
    geom_line(mapping=aes(x=LT,y=power,color=type,group=type))+
    geom_errorbar(mapping=aes(x=LT,ymin=lower,ymax=upper,color=type,group=type),width=0.2)+
    coord_cartesian(ylim=c(0,1))+
    scale_color_manual(values = tree.type.colors,labels=tree.type.labels)+
    scale_shape_discrete(labels=tree.type.labels)+
    scale_x_discrete(breaks=c(0,0.01,0.05,0.10,0.50,1.00),
                     labels=c("True\ntree","0.01","0.05","0.1","0.5","1"))+
    facet_wrap(facet= vars(method_label),nrow = 1,labeller = label_parsed)+
    xlab("Lineage tracing editing rate")+ylab("Power")+
      theme(legend.position = "none",
            strip.background = element_rect(fill = model.color[mdl]),
            strip.text = element_text(color="white",face="bold",size=8,margin=unit(c(0,0,0,0),units = "points")),
            panel.background = element_blank(),
            panel.grid = element_line(color="grey",linetype = "dotted"),
            axis.line = element_line(),plot.margin=unit(c(1,1,1,1),units = "points"),
            axis.title = element_text(size=10),
            axis.text = element_text(size=8))
    if(mdl=="CRH") this.plot = this.plot+theme(axis.title.x = element_blank())
    return(this.plot)
})

dummy.plot = ggplot(data=data.frame(x=1:3,y=1:3,color=c("TGT","LTT","TGT"),
                                    model=factor(c("CRH","CRH","DRH"),levels=names(model.color))))+
  geom_line(mapping = aes(x=x,y=y,color=color,group=color))+
  geom_point(mapping = aes(x=x,y=y,color=color,shape=color))+
  geom_col(mapping = aes(x=x,y=y,fill=model))+
  scale_color_manual(name="Tree type", values = tree.type.colors,labels=tree.type.labels)+
  scale_shape_discrete(name="Tree type",labels=tree.type.labels)+
  scale_fill_manual(name="Growth model",values=model.color)+
  guides(color=guide_legend(direction = "horizontal"),
         fill=guide_legend(direction = "horizontal"))+
  theme(legend.text = element_text(size=8),
        legend.title = element_text(size=10),
        legend.key = element_blank(),
        legend.direction = "horizontal",
        legend.margin = margin(t = 3),
        legend.box.margin = margin(),
        legend.spacing = unit(12,units="points"))

fig3 = ggarrange(plotlist=do.call(mapply,list(FUN=list,SIMPLIFY=TRUE,fig3a.panels,fig3b.panels)),
                                  ncol = 2,nrow = 2,heights = c(35,35),
                 align = "hv",common.legend = TRUE,legend = "bottom",labels = c("A","B","",""),
                 label.x = -0.01,label.y = 1.01,
                 legend.grob = get_legend(dummy.plot,position = "bottom"))
#
ggsave(plot = fig3,filename = "Figure_3.png",path ="Figures_and_Tables/",device = "png",
       width = 185,height = 105,scale = 1,units = "mm",dpi = 300)
ggsave(plot = fig3,filename = "Figure_3.pdf",path ="Figures_and_Tables/",device = "pdf",
       width = 185,height = 105,scale = 1,units = "mm",colormodel="cmyk")

figS4.panels = lapply(c("CRH","DRH"),function(mdl){
  selected.mdl =plot.data[[mdl]]$method=="J1"
  selected.EBR = plot.data$EBR$method=="J1"
  this.data = rbind(plot.data[[mdl]][selected.mdl,],plot.data$EBR[selected.EBR,])
  this.data$hetero_by_type = paste0(this.data$type,"_",this.data$hetero)
  this.hetero = sort(unique(this.data$hetero))
  this.hetero = c(this.hetero[-1],this.hetero[1])
  this.data$hetero = factor(this.data$hetero,levels = this.hetero)
  this.plot = ggplot(data = this.data)+
    geom_point(mapping=aes(x=LT,y=power,color=hetero,shape=hetero))+
    geom_line(mapping=aes(x=LT,y=power,color=hetero,group=hetero_by_type))+
    geom_errorbar(mapping=aes(x=LT,ymin=lower,ymax=upper,color=hetero,group=hetero_by_type),width=0.2)+
    geom_hline(yintercept = 0.05,color="black",linetype="dashed")+
    scale_x_discrete(breaks=c(0,0.01,0.05,0.10,0.50,1.00),
                     labels=c("True\ntree","0.01","0.05","0.1","0.5","1"))+
    scale_color_manual(values = hetero.color,drop=FALSE)+
    scale_shape_manual(values = hetero.shape,drop=FALSE)+
    coord_cartesian(ylim=c(0,1))+
    xlab("Lineage tracing editing rate")+ylab("Power")+
    guides(color=guide_legend(title = "Strength of rate heterogeneity",nrow = 1))+
    facet_wrap(facets = vars(n),nrow = 1,
               labeller = labeller(n=c("50"="n=50","250"="n=250","1250"="n=1250")))+
    theme(legend.position = "none",
          strip.background = element_rect(fill = model.color[mdl]),
          strip.text = element_text(color="white",face="bold",size=8,margin=unit(c(0,0,0,0),units = "points")),
          panel.background = element_blank(),
          panel.grid = element_line(color="grey",linetype = "dotted"),
          axis.line = element_line(),plot.margin=unit(c(1,1,1,1),units = "points"),
          axis.title = element_text(size=10),
          axis.text = element_text(size=8))
  if(mdl=="CRH") this.plot = this.plot+theme(axis.title.x = element_blank())
  return(this.plot)
})
#
dummyS.plot = ggplot(
  data=data.frame(x=1:10,y=1:10,
                  color=factor(as.numeric(names(hetero.color)),
                               levels = as.numeric(names(hetero.color))[c(2,7,3,8,4,9,5,10,6,1)]),
                  model=factor(rep(c("CRH","DRH"),5),
                               levels=names(model.color))))+
  geom_line(mapping = aes(x=x,y=y,color=color,group=color))+
  geom_point(mapping = aes(x=x,y=y,color=color,shape=color))+
  geom_col(mapping = aes(x=x,y=y,fill=model))+
  scale_color_manual(name="Strength of rate heterogeneity", values = hetero.color,
                     labels=hetero.label)+
  scale_shape_manual(name="Strength of rate heterogeneity",values = hetero.shape,
                     labels=hetero.label)+
  scale_fill_manual(name="Growth model",values=model.color)+
  guides(color=guide_legend(title = "Strength of rate heterogeneity",nrow = 2,order = 2,direction = "vertical"),
         shape=guide_legend(title = "Strength of rate heterogeneity",nrow = 2,order = 2,direction = "vertical"),
         fill=guide_legend(order = 1,direction="vertical"))+
  theme(legend.text = element_text(size=8),
        legend.title = element_text(size=10),
        legend.key = element_blank(),
        legend.direction = "horizontal",
        legend.margin = margin(t = 3),
        legend.box.margin = margin(),
        legend.spacing = unit(12,units="points"))
#
figS4 = ggarrange(plotlist = figS4.panels,ncol = 1,nrow = 2,
                 align = "hv",common.legend = TRUE,legend = "bottom",
                 legend.grob = get_legend(dummyS.plot,position="bottom"))+
  theme(plot.margin = margin(t = 2,r = 2,b = 2,l = 12,unit = "pt"))
#
ggsave(plot = figS4,filename = "Figure_S4.png",path ="Figures_and_Tables/",device = "png",
       width = 185,height = 120,scale = 1,units = "mm",dpi = 300)
ggsave(plot = figS4,filename = "Figure_S4.pdf",path ="Figures_and_Tables/",device = "pdf",
       width = 185,height = 120,scale = 1,units = "mm",colormodel="cmyk")
```
