## Supplementary Data and Scripts for "Detecting branching rate heterogeneity with tree balance statistics in lineage tracing trees": Fig_4_S7_Testing_empirical_trees.R

```
# load packages
library(ape)
library(ggplot2)
library(ggpubr)
library(ggtree)
source("Scripts/Fig_Color_Palettes.R")

# read in test outcomes
empirical.test = readRDS("Empirical_data/empirical_res.RDS")
empirical.tree = mapply(FUN = function(id,group){
  ff = sprintf("Empirical_data/%s/%s__MaxCut.tre",group,id)
  read.tree(ff)
},
SIMPLIFY = FALSE,empirical.test$tree_id,empirical.test$group)

# trim trees to 100 tips for display
set.seed(20250226)
plot.tree = lapply(empirical.tree,function(phy){
  if(Ntip(phy)>100){
    phy = keep.tip(phy = phy,tip = sample(phy$tip.label,size = 100,replace = FALSE))
  }
  phy$edge.length = phy$edge.length*0+1
  phy$root.edge = 1
  return(phy)
})
full.plot.tree = lapply(empirical.tree,function(phy){
  phy$edge.length = phy$edge.length*0+1
  phy$root.edge = 1
  return(phy)
})
ph.tree = rtree(n = 3)
plot.tree$placeholder=ph.tree
class(plot.tree) = "multiPhylo"
class(full.plot.tree) = "multiPhylo"

# set up tree colors
empirical.test$label = paste0(empirical.test$group,"\n",empirical.test$tree_id)
names(plot.tree) = empirical.test$label
names(full.plot.tree) = empirical.test$label
phy.color = significance.color[as.numeric(empirical.test$p.value<=0.05)+1]
names(phy.color) = empirical.test$label
phy.color = c(phy.color,c(placeholder="white"))

# ggtree objects for empirical trees
tree.combined= ggtree(tr = plot.tree[
  c(empirical.test$label[empirical.test$group=="Lung_auto"],
    empirical.test$label[empirical.test$group=="Lung_xeno"],
    empirical.test$label[empirical.test$group=="PDAC"&empirical.test$estimated_LT<0.5])
  ],
  mapping=aes(color=.id),layout = "fan",open.angle = 0)+
  scale_color_manual(values = phy.color)+
  facet_wrap(facets = ~.id,ncol=7,scales = "free")+
  theme(legend.position = "none",
        panel.spacing = unit(1,units = "points"),
        strip.background = element_rect(fill = "transparent",color="black"),
        strip.text = element_text(color="black",face="bold",size=8,margin=unit(c(1,0,2,0),units = "points")),
        plot.margin=unit(c(6,1,1,1),units = "points"))

# save figures
ggsave(plot = tree.combined,filename = "Figures_and_Tables/Figure_4.png",
       device = "png",width = 185,height = 165,dpi = 300,units = "mm")
ggsave(plot = tree.combined,filename = "Figures_and_Tables/Figure_4.pdf",
       device = "pdf",width = 185,height = 165,units = "mm",colormodel="cmyk")

# ggtree objects for full empirical trees
full.tree.combined= 
  lapply(c(empirical.test$label[empirical.test$group=="Lung_auto"],
           empirical.test$label[empirical.test$group=="Lung_xeno"],
           empirical.test$label[empirical.test$group=="PDAC"&empirical.test$estimated_LT<0.5]),
         function(nn){
           ggtree(tr = full.plot.tree[nn],
                  mapping=aes(color=.id),layout = "fan",open.angle = 0)+
             scale_color_manual(values = phy.color)+
             facet_wrap(facets = ~.id,ncol=5,scales = "free",)+
             theme(legend.position = "none",
                   panel.spacing = unit(1,units = "points"),
                   strip.background = element_rect(fill = "transparent",color="black"),
                   strip.text = element_text(color="black",face="bold",size=8,margin=unit(c(1,0,2,0),units = "points")),
                   plot.margin=unit(c(2,0,0,0),units = "points"))
           })
figS7= ggarrange(plotlist = full.tree.combined,ncol = 5,nrow = 7,align = "hv",legend = "none")

# save supplementary figures
ggsave(plot = figS7,filename = "Figures_and_Tables/Figure_S7.png",
       device = "png",width = 185,height = 275,dpi = 300,units = "mm")
ggsave(plot = figS7,filename = "Figures_and_Tables/Figure_S7.pdf",
       device = "pdf",width = 185,height = 275,units = "mm",colormodel="cmyk")
```
