## Supplementary Data and Scripts for "Detecting branching rate heterogeneity with tree balance statistics in lineage tracing trees": Fig_Color_Palettes.R

```
# This file contains the definition of colors so that they are consistent across
# all figures

# color by model
model.color = c(EBR="black",CRH="orange",DRH="#377eb8")
model.order = c("EBR","CRH","DRH")

# color by statistics
tbs.color = c(J1="#e9a3c9",Sackin="#a1d76a")
tbs.label = c(J1=parse(text = "J^1"),Sackin="Sackin index")

# setup color schemes for CM
CM.colors= c("white",c("cyan","darkblue","#4daf4a","#984ea3","#ff7f00","#ffff33","#a65628","#f781bf"))
names(CM.colors) = (1:length(CM.colors))-1

# set up color schemes and labels for polytomy composition
tree.structure.color = c("100"="#bd0026","010"="#7570b3","001"="#1b9e77","000"="#a9a9a9")
tree.highlight.color = c("100"="#bd0026","010"="#7570b3","001"="#1b9e77","000"="#a9a9a9")
tree.structure.alpha = c("100"=1,"010"=1,"001"=1,"000"=0.75)
tree.structure.label = c("100"="Star","010"="Star-like","001"="Other polytomy","000"="Binary split")

# set up color schemes for tree type: genealogical vs lineage tracing
tree.type.colors = c(TGT="#1b9e77",LTT="#d95f02")
tree.type.labels=c(TGT="Genealogical tree",LTT="Lineage tracing tree")

# set up color schemes for branching rate heterogeneity
hetero.color = c("0"="black","0.1"="#5aae61","0.5"="#d9f0d3",
                 "1"="#c2a5cf","5"="#9970ab","10"="#762a83",
                 "0.062"="#5aae61","0.126"="#a6dba0",
                 "0.312"="#d9f0d3","0.624"="#c2a5cf")
hetero.shape = c("0"=19,"0.1"=15,"0.5"=15,"1"=15,"5"=15,"10"=15,
                 "0.062"=17,"0.126"=17,"0.312"=17,"0.624"=17)
hetero.label = c("0"="0 (Type I error)","0.1"="0.1","0.5"="0.5","1"="1","5"="5","10"="10",
                 "0.062"="0.062","0.126"="0.126","0.312"="0.312","0.624"="0.624")

# color scheme for test significance
significance.color = c("black","#d7191c")
```
