## Supplementary Data and Scripts for "Detecting branching rate heterogeneity with tree balance statistics in lineage tracing trees": Fig_S1_Effect_of_tree_size_on_power.R

```
#
# load packages
library(ggplot2)
library(ggpubr)
source("Scripts/Fig_Color_Palettes.R")

# read in J1 and Sackin across models
J1_Sackin_stat = readRDS("Simulated_data/stats/J1_Sackin_power.RDS")
J1_Sackin_stat = J1_Sackin_stat[J1_Sackin_stat$type=="TGT",]

# pre-compute CI for all possible p^ in the experimental design
CI.data = do.call(rbind,lapply(0:1000,function(x){
  this.test = prop.test(x = x,n = 1000,p = 0.05)
  data.frame(phat=x/1000,pup=this.test$conf.int[2],plow=this.test$conf.int[1])
}))

# title, color, and adjustment settings
full.title = c(EBR="Equal branching\nrate",
               CRH="Continuous rate\nheterogeneity",
               DRH="Discrete rate\nheterogeneity")
method.power.adjust = c(J1=+0.004,Sackin=-0.004)

# Adjust data
J1_Sackin_stat$hetero=factor(J1_Sackin_stat$hetero,levels=sort(unique(J1_Sackin_stat$hetero)))
J1_Sackin_stat$upper = CI.data$pup[match(round(J1_Sackin_stat$power,digits = 3),round(CI.data$phat,digits = 3))]
J1_Sackin_stat$lower = CI.data$plow[match(round(J1_Sackin_stat$power,digits = 3),round(CI.data$phat,digits = 3))]
J1_Sackin_stat$power = J1_Sackin_stat$power+method.power.adjust[J1_Sackin_stat$method]
J1_Sackin_stat$upper = J1_Sackin_stat$upper+method.power.adjust[J1_Sackin_stat$method]
J1_Sackin_stat$lower = J1_Sackin_stat$lower+method.power.adjust[J1_Sackin_stat$method]

figS1.data = do.call(rbind,lapply(c("CRH","DRH"),function(mdl){
  this.data = rbind(
    J1_Sackin_stat[J1_Sackin_stat$model==mdl,],
    J1_Sackin_stat[J1_Sackin_stat$model=="EBR",])
  this.data$facet_model=mdl
  return(this.data)
}))

# 
figS1.panels = ggplot(data = figS1.data)+
    geom_line(mapping=aes(x=hetero,y=power,group=method,color=method))+
    geom_point(mapping=aes(x=hetero,y=power,group=method,color=method,shape=method))+
    geom_errorbar(mapping=aes(x=hetero,ymin=lower,ymax=upper,color=method,group=method),width=0.2)+
    geom_hline(yintercept = 0.05, color="black",linetype="dashed")+
    ggh4x::facet_grid2(cols=vars(n),rows=vars(facet_model),scales = "free_x",independent = "x",
               labeller = labeller(n=c("50"="n=50","250"="n=250","1250"="n=1250"),
                                   facet_model=c(CRH="CRH",DRH="DRH")))+
    scale_color_manual(name="Tree balance statistics",values = tbs.color,labels=tbs.label)+
    scale_shape_discrete(name="Tree balance statistics",labels=tbs.label)+
    coord_cartesian(ylim=c(0,1))+
    xlab("Strength of rate heterogeneity")+ylab("Power")+
  theme(legend.position = "none",
        axis.title = element_text(size=10),
        axis.text = element_text(size=8),
        legend.text = element_text(size = 8),
        legend.title = element_text(size=10),
        panel.background = element_blank(),
        panel.spacing.x = unit(4,units = "points"),
        strip.background.y = element_rect(fill="transparent"),
        strip.text.y = element_text(color="white"),
        strip.background.x = element_rect(fill = "transparent",color="black"),
        strip.text.x = element_text(size = 8,face = "bold",margin = margin(1,0,1,0)),
        panel.grid = element_line(color="grey",linetype = "dotted"),
        axis.line = element_line(),plot.margin = unit(c(10,10,10,10),units = "points"))

#
dummy.fig = ggplot(data=data.frame(x=1:2,y=1:2,color=c("J1","Sackin"),model=c("CRH","DRH")))+
  geom_line(mapping = aes(x=x,y=y,color=color,group=color))+
  geom_point(mapping = aes(x=x,y=y,color=color,shape=color))+
  geom_col(mapping = aes(x=x,y=y,fill=model))+
  scale_color_manual(name="Tree balance statistics",values = tbs.color,
                     labels=tbs.label)+
  scale_shape_discrete(name="Tree balance statistics",labels=tbs.label)+
  scale_fill_manual(name="Growth model",values=model.color)+
  guides(color=guide_legend(direction = "horizontal",order = 2),
         fill=guide_legend(direction = "horizontal",order = 1),
         shape=guide_legend(direction = "horizontal",order = 2))+
  theme(legend.direction = "horizontal")

# arrange the plot in a master canvas
figS1 = ggplot()+
  coord_cartesian(xlim = c(-1,186),ylim = c(-1,136),expand = FALSE)+
  theme(panel.background = element_blank(),
        axis.line = element_blank(),axis.ticks = element_blank(),
        axis.text = element_blank(),axis.title = element_blank(),
        legend.position = "none",plot.margin = unit(c(0,0,0,0),units = "points"))
# add panels and legends
figS1 = figS1+
  annotation_custom(grob = cowplot::as_grob(figS1.panels),
                    xmin = 0,xmax = 185,ymin = 15,ymax=135)+
  annotation_custom(grob=cowplot::as_grob(get_legend(dummy.fig,position="bottom")),
                    xmin=0,xmax=185,ymin=0,ymax=15)+
  annotate(geom="rect",xmin=175,xmax=179,ymin=81,ymax = 128,fill=model.color[2])+
  annotate(geom="rect",xmin=175,xmax=179,ymin=27,ymax = 74,fill=model.color[3])+
  annotate(geom="text",x=177,y= 104.5,label=names(model.color)[2],
           color="white",fontface="bold",size=3,angle=270)+
  annotate(geom="text",x=177,y= 50.5,label=names(model.color)[3],
           color="white",fontface="bold",size=3,angle=270)

# save to file
ggsave(plot = figS1,path = "Figures_and_Tables/",filename = "Figure_S1.png",device = "png",
       height = 135,width = 185,scale = 1,units = "mm",dpi = 300)
ggsave(plot = figS1,path = "Figures_and_Tables/",filename = "Figure_S1.pdf",device = "pdf",
       height = 135,width = 185,scale = 1,units = "mm",colormodel="cmyk")
#
```
