## Supplementary Data and Scripts for "Detecting branching rate heterogeneity with tree balance statistics in lineage tracing trees": Fig_S2_More_effects_of_LT_on_trees.R

```
# load packages
library(ggplot2)
library(ggpubr)
source("Scripts/Fig_Color_Palettes.R")

#
tree.props=readRDS("Simulated_data/stats/LT_topology_and_examples.RDS")

# generate the full panels for polytomy compositions
polytomy.counts = do.call(rbind,lapply(tree.props,function(x){
  x$node$freq = x$node$x/sum(x$node$x)
  return(x$node)
}))
polytomy.counts$edit_rate = factor(polytomy.counts$edit_rate,c(0,0.01,0.05,0.10,0.50,1.00))

# generate the full plot for supplementary material
figS2a = ggplot(polytomy.counts[polytomy.counts$edit_rate!=0,])+
  geom_bar(mapping=aes(x=edit_rate,y=freq,fill=type,alpha=type),stat = "identity")+
  facet_grid(cols = vars(size), rows =vars(model),
             labeller = labeller(size=c("50"="n=50","250"="n=250","1250"="n=1250"),
                                 model=c(EBR="EBR",CRH="CRH",DRH="DRH")))+
  scale_fill_manual(name="Polytomy (Panel A)",values = tree.structure.color,labels=tree.structure.label)+
  scale_alpha_manual(name="Polytomy (Panel A)",values = tree.structure.alpha,labels=tree.structure.label)+
  xlab("Lineage tracing editing rate")+ylab("Frequency")+
  coord_cartesian(ylim = c(0,0.5),expand = TRUE)+
  guides(fill=guide_legend(reverse = TRUE),
         alpha=guide_legend(reverse = TRUE))+
  theme(legend.position = "none",
        panel.background = element_blank(),
        panel.grid = element_line(color="grey",linetype = "dotted"),
        strip.background = element_rect(fill="transparent",color="black"),
        strip.background.y = element_rect(fill="transparent",color="transparent"),
        strip.text = element_text(face="bold",size=8,margin=unit(c(1,0,1,0),units = "points")),
        strip.text.y = element_text(color="white"),
        axis.line = element_line(),
        axis.title = element_text(size=10),
        axis.text = element_text(size=8),
        plot.margin = unit(c(4,1,4,1),units = "points"))

# generate node-to-tip ratio and root balance data for panel D and E
figS2bc.data = do.call(rbind,lapply(tree.props,function(x){
  xx = x$tree
  data.frame(model=xx$model[1],edit_rate=xx$edit_rate[1],size=xx$size[1],
             pseudo_facet="PH",
             n2t.mean = mean(xx$node2tip),
             n2t.upper=quantile(xx$node2tip,probs=0.75),
             n2t.lower=quantile(xx$node2tip,probs=0.25),
             rtb.mean = mean(log(xx$root_balance)),
             rtb.upper=quantile(log(xx$root_balance),probs=0.75),
             rtb.lower=quantile(log(xx$root_balance),probs=0.25))
}))
figS2bc.data$edit_rate = factor(figS2bc.data$edit_rate,levels = c(0,0.01,0.05,0.1,0.5,1))

# generate panel D of node-to-tip ratio
figS2b = ggplot(data=figS2bc.data)+
  geom_point(mapping=aes(x=edit_rate,y=n2t.mean,color=model,group=model))+
  geom_errorbar(mapping=aes(x=edit_rate,y=n2t.mean,ymin=n2t.lower,ymax=n2t.upper,color=model),
                position = position_dodge(width = 0.1))+
  geom_line(data=figS2bc.data[figS2bc.data$edit_rate!=0,],
            mapping=aes(x=edit_rate,y=n2t.mean,color=model,group=model),linewidth=1)+
  facet_grid(cols = vars(size), rows =vars(pseudo_facet),
             labeller = labeller(size=c("50"="n=50","250"="n=250","1250"="n=1250"),
                                 pseudo_facet=c(PH="Placeholder")))+
  scale_color_manual(name="Model",values = model.color,labels=names(model.color))+
  scale_x_discrete(breaks=c(0,0.01,0.05,0.10,0.50,1.00),
                   labels=c("True\ntree","0.01","0.05","0.1","0.5","1"))+
  xlab("    ")+ylab("Node-to-tip ratio")+
  coord_cartesian(ylim = c(0,1),expand = TRUE)+
  theme(legend.position = "none",
        legend.key.size = unit(c(8,8),units="points"),
        legend.text = element_text(size = 8),
        legend.title = element_text(size=10),
        panel.background = element_blank(),
        panel.grid = element_line(color="grey",linetype = "dotted"),
        strip.background = element_rect(fill="transparent",color="black"),
        strip.background.y = element_rect(fill="transparent",color="transparent"),
        strip.text = element_text(face="bold",size=8,margin=unit(c(1,0,1,0),units = "points")),
        strip.text.y = element_text(color="transparent"),
        axis.line = element_line(),
        axis.title = element_text(size=10),
        axis.text = element_text(size=8),
        plot.margin = unit(c(4,1,4,1),units = "points"))

# generate panel E of root balance
figS2c = ggplot(data=figS2bc.data)+
  geom_point(mapping=aes(x=edit_rate,y=rtb.mean,color=model,group=model))+
  geom_errorbar(mapping=aes(x=edit_rate,y=rtb.mean,ymin=rtb.lower,ymax=rtb.upper,color=model),
                position = position_dodge(width = 0.1))+
  geom_line(data=figS2bc.data[figS2bc.data$edit_rate!=0,],
            mapping=aes(x=edit_rate,y=rtb.mean,color=model,group=model),linewidth=1)+
  facet_grid(cols = vars(size), rows =vars(pseudo_facet),
             labeller = labeller(size=c("50"="n=50","250"="n=250","1250"="n=1250"),
                                 pseudo_facet=c(PH="Placeholder")))+
  scale_color_manual(name="Model",values = model.color,labels=names(model.color))+
  scale_x_discrete(breaks=c(0,0.01,0.05,0.10,0.50,1.00),
                   labels=c("True\ntree","0.01","0.05","0.1","0.5","1"))+
  xlab("Lineage tracing editing rate")+ylab("Sister size ratio")+
  #coord_cartesian(ylim = c(0,1),expand = TRUE)+
  theme(legend.position = "none",
        legend.key.size = unit(c(8,8),units="points"),
        legend.text = element_text(size = 8),
        legend.title = element_text(size=10),
        panel.background = element_blank(),
        panel.grid = element_line(color="grey",linetype = "dotted"),
        strip.background = element_rect(fill="transparent",color="black"),
        strip.background.y = element_rect(fill="transparent",color="transparent"),
        strip.text = element_text(face="bold",size=8,margin=unit(c(1,0,1,0),units = "points")),
        strip.text.y = element_text(color="transparent"),
        axis.line = element_line(),
        axis.title = element_text(size=10),
        axis.text = element_text(size=8),
        plot.margin = unit(c(4,1,4,1),units = "points"))

figS2abc = ggarrange(plotlist = list(figS2a,figS2b,figS2c),ncol = 1,nrow = 3,
                     align = "v",legend = "none",common.legend = TRUE,
                     heights = c(117,45,45),labels = "AUTO",label.x = -0.01,label.y = 1.02)
#
dummy.plot = ggplot(polytomy.counts)+
  geom_bar(mapping=aes(x=size,y=freq,fill=type,alpha=type),stat="identity")+
  geom_line(mapping=aes(x=size,y=freq,color=model,group=edit_rate))+
  scale_color_manual(name="Growth model",values = model.color,labels=names(model.color))+
  scale_fill_manual(name="Polytomy",values = tree.structure.color,labels=tree.structure.label)+
  scale_alpha_manual(name="Polytomy",values = tree.structure.alpha,labels=tree.structure.label)+
  guides(fill=guide_legend(reverse = TRUE,direction = "horizontal"),
         alpha=guide_legend(reverse = TRUE,direction = "horizontal"),
         color=guide_legend(direction = "horizontal"))+
  theme(legend.position = "bottom",
        legend.box.background = element_rect(fill="transparent",color="transparent"),
        legend.text = element_text(size = 8),
        legend.title = element_text(size=10),
        legend.box = "vertical",
        legend.spacing.y = unit(4,units="points"),
        panel.background = element_blank(),
        panel.grid = element_line(color="grey",linetype = "dotted"),
        plot.margin = unit(c(1,1,1,1),units = "points"))

# arrange full plot in a master canvas
figS2 = ggplot()+
  coord_cartesian(xlim = c(-1,181),ylim = c(-1,235),expand = FALSE)+
  theme(panel.background = element_blank(),
        axis.line = element_blank(),axis.ticks = element_blank(),
        axis.text = element_blank(),axis.title = element_blank(),
        legend.position = "none",plot.margin = unit(c(0,0,0,0),units = "points"))

# add each panel and panel-specific annotations
figS2 = figS2+
  annotation_custom(grob = cowplot::as_grob(figS2abc),xmin = 0,xmax = 180,ymin = 25,ymax = 234)+
  annotate(geom="rect",xmin=177,xmax=180,ymin=197.5,ymax = 229.5,fill=model.color[1])+
  annotate(geom="rect",xmin=177,xmax=180,ymin=162,ymax = 194,fill=model.color[2])+
  annotate(geom="rect",xmin=177,xmax=180,ymin=127,ymax = 159,fill=model.color[3])+
  annotate(geom="text",x=178.5,y= 213.5,label=names(model.color)[1],
           color="white",fontface="bold",size=3,angle=270)+
  annotate(geom="text",x=178.5,y= 178,label=names(model.color)[2],
           color="white",fontface="bold",size=3,angle=270)+
  annotate(geom="text",x=178.5,y= 142,label=names(model.color)[3],
           color="white",fontface="bold",size=3,angle=270)+
  annotation_custom(grob=cowplot::as_grob(get_legend(dummy.plot)),
                    xmin = 20,xmax = 160,ymin = 0,ymax = 25)

# save supplementary figure to file
# for online preview
ggsave(plot = figS2,path = "Figures_and_Tables/",filename = "Figure_S2.png",device = "png",
       height = 234,width = 185,scale = 1,units = "mm",dpi = 300)
# for submission
ggsave(plot = figS2,path = "Figures_and_Tables/",filename = "Figure_S2.pdf",device = "pdf",
       height = 234,width = 185,scale = 1,units = "mm",colormodel = "cmyk")
```
