## Supplementary Data and Scripts for "Detecting branching rate heterogeneity with tree balance statistics in lineage tracing trees": Fig_S3_More_effects_of_LT_on_tree_balance.R

```
# load packages
library(ggplot2)
library(ggpubr)
source("Scripts/Fig_Color_Palettes.R")

# read in J1 and Sackin across models for panel F
J1_Sackin_res = readRDS("Simulated_data/stats/J1_Sackin_data.RDS")
axis.labels = c(J1="J^1",Sackin="Sackin~~index")

# organize data table, generate pseudo-Sackin for plotting
figS3.data = J1_Sackin_res[J1_Sackin_res$hetero%in%c(0,1),]
figS3.data = split(figS3.data,figS3.data$model)
figS3.data = do.call(rbind,lapply(c("CRH","DRH"),function(mdl){
  this.data = do.call(rbind,figS3.data[c(mdl,"EBR")])
  this.data$facet_model=mdl
  return(this.data)
}))
figS3.data$model = factor(figS3.data$model,levels = names(model.color))
figS3.data$LT = factor(figS3.data$LT,levels = sort(unique(figS3.data$LT)))
figS3.data$group = paste0(figS3.data$model,"_",figS3.data$LT)
figS3.data = split(figS3.data,figS3.data$method)

# generate Figure S3
figS3.panels = lapply(names(figS3.data),function(mm){
  ggplot(data=figS3.data[[mm]])+
    geom_violin(mapping=aes(x=LT,y=stat,group=group,fill=model,color=model))+
    scale_color_manual(values=model.color,name="Growth model",drop=FALSE)+
    scale_fill_manual(values=model.color,name="Growth model",drop=FALSE)+
    ggh4x::facet_grid2(cols=vars(n),rows = vars(facet_model),scales = "free_y",independent = "y",
                       labeller = labeller(facet_model=c(CRH="CRH",DRH="DRH"),
                                           n=c("50"="n=50","250"="n=250","1250"="n=1250")))+
    scale_x_discrete(breaks=c(0,0.01,0.05,0.10,0.50,1.00),
                     labels=c("True\ntree","0.01","0.05","0.1","0.5","1"))+
    xlab("Lineage tracing editing rate")+ylab(parse(text=axis.labels[mm]))+
    theme(legend.position = "bottom",
          axis.title = element_text(size=10),
          axis.text = element_text(size=8),
          legend.text = element_text(size = 8),
          legend.title = element_text(size=10),
          panel.background = element_blank(),
          panel.spacing.x = unit(4,units = "points"),
          strip.background.y = element_blank(),
          strip.text.y = element_blank(),
          strip.background.x = element_rect(fill = "transparent",color="black"),
          strip.text.x = element_text(size = 8,face = "bold",margin = margin(1,0,1,0)),
          panel.grid = element_line(color="grey",linetype = "dotted"),
          axis.line = element_line(),plot.margin = unit(c(10,10,10,10),units = "points"))
})
figS3 = ggarrange(plotlist=figS3.panels,ncol = 1,nrow = 2,
                  labels = "AUTO",label.x = -0.01,label.y = 1.01,
                  align = "hv",legend = "bottom",common.legend = TRUE)

# save supplementary figure to file
# for online preview
ggsave(plot = figS3,path = "Figures_and_Tables/",filename = "Figure_S3.png",device = "png",
       height = 200,width = 185,scale = 1,units = "mm",dpi = 300)
# for submission
ggsave(plot = figS3,path = "Figures_and_Tables/",filename = "Figure_S3.pdf",device = "pdf",
       height = 200,width = 185,scale = 1,units = "mm",colormodel = "cmyk")
```
