## Supplementary Data and Scripts for "Detecting branching rate heterogeneity with tree balance statistics in lineage tracing trees": Fig_S5_Robustness_of_J1_against_misspecification.R

```
#
# load packages
library(ggplot2)
library(ggpubr)
source("Scripts/Fig_Color_Palettes.R")

# structure required data
J1_robustness_stat = readRDS("Simulated_data/stats/J1_robustness_power.RDS")
plot.data = J1_robustness_stat[J1_robustness_stat$type=="LTT"&J1_robustness_stat$null_type=="LTT",]
plot.data$hetero = factor(plot.data$hetero,levels = sort(unique(plot.data$hetero)))
null_N.range = sort(unique(plot.data$null_N))
null_N.label = unname(sapply(c("1250","6250","31250","10^6"),function(x){parse(text=x)}))
null_edit_rate.range = sort(unique(plot.data$null_edit_rate))

# pre-compute CI for all possible p^ in the experimental design
CI.data = do.call(rbind,lapply(0:1000,function(x){
  this.test = prop.test(x = x,n = 1000,p = 0.05)
  data.frame(phat=x/1000,pup=this.test$conf.int[2],plow=this.test$conf.int[1])
}))
plot.data$upper = CI.data$pup[match(plot.data$power,CI.data$phat)]
plot.data$lower = CI.data$plow[match(plot.data$power,CI.data$phat)]

# label for each editing rate
rate.label = c("0.01"="mu[edit]==0.01",
               "0.05"="mu[edit]==0.05",
               "0.1"="mu[edit]==0.10",
               "0.5"="mu[edit]==0.50",
               "1"="mu[edit]==1.00")

# Dummy plot for legend
dummy.plot = ggplot(
  data=data.frame(x=1:10,y=1:10,
                  color=factor(as.numeric(names(hetero.color)),
                               levels = as.numeric(names(hetero.color))[c(2,7,3,8,4,9,5,10,6,1)]),
                  model=factor(rep(c("CRH","DRH"),5),
                               levels=names(model.color))))+
  geom_line(mapping = aes(x=x,y=y,color=color,group=color))+
  geom_point(mapping = aes(x=x,y=y,color=color,shape=color))+
  geom_col(mapping = aes(x=x,y=y,fill=model))+
  scale_color_manual(name="Strength of rate heterogeneity", values = hetero.color,
                     labels=hetero.label)+
  scale_shape_manual(name="Strength of rate heterogeneity",values = hetero.shape,
                     labels=hetero.label)+
  scale_fill_manual(name="Growth model",values=model.color)+
  guides(color=guide_legend(title = "Strength of rate heterogeneity",nrow = 2,order = 2,direction = "vertical"),
         shape=guide_legend(title = "Strength of rate heterogeneity",nrow = 2,order = 2,direction = "vertical"),
         fill=guide_legend(order = 1,direction="vertical"))+
  theme(legend.text = element_text(size=8),
        legend.title = element_text(size=10),
        legend.key = element_blank(),
        legend.direction = "horizontal",
        legend.margin = margin(t = 3),
        legend.box.margin = margin(),
        legend.spacing = unit(12,units="points"))

# generate panel A-specific data
figS5a.data = do.call(rbind,lapply(c("CRH","DRH"),function(mdl){
  selected.rows = plot.data$model%in%c(mdl,"EBR")&plot.data$LT%in%c(0.01,0.1,1)&
    plot.data$n==1250&plot.data$null_N==6250&plot.data$null_n==1250
  df = plot.data[selected.rows,]
  df$pseudo_facet = mdl
  df$label = rate.label[as.character(df$LT)]
  df$null_edit_rate = factor(df$null_edit_rate,levels = null_edit_rate.range)
  df = df[df$LT%in%c(0.01,0.1,1),]
  df$LT = factor(df$LT,levels = null_edit_rate.range)
  return(df)
}))
# make panel A
figS5a = ggplot(figS5a.data)+
  geom_point(mapping=aes(x=null_edit_rate,y=power,color=hetero,shape=hetero))+
  geom_line(mapping=aes(x=null_edit_rate,y=power,color=hetero,group=hetero))+
  geom_errorbar(mapping=aes(x=null_edit_rate,ymin=lower,ymax=upper,color=hetero,group=hetero),width=0.2)+
  geom_vline(data = figS5a.data[figS5a.data$hetero==0&figS5a.data$null_edit_rate==0.1,],
             mapping = aes(xintercept=LT),color="black",linetype="dashed")+
  geom_hline(yintercept = 0.05,linetype="dashed",color="black")+
  facet_grid(rows=vars(pseudo_facet),cols=vars(label),labeller = label_parsed)+
  scale_color_manual(name = "Strength of rate\nheterogeneity",values = hetero.color,drop=FALSE)+
  scale_shape_manual(name = "Strength of rate\nheterogeneity",values = hetero.shape,drop=FALSE)+
  xlab("Editing rate for generating null distribution")+ylab("Power")+
  theme(legend.position = "none",
        axis.title = element_text(size=10),
        axis.text = element_text(size=8),
        legend.text = element_text(size = 8),
        legend.title = element_text(size=10),
        panel.background = element_blank(),
        panel.spacing.x = unit(12,units = "points"),
        strip.background.x = element_rect(fill="transparent",color="black"),
        strip.text.x = element_text(size=8,face = "bold",margin = margin(1,0,1,0)),
        strip.background.y = element_blank(),
        strip.text.y = element_blank(),
        panel.grid = element_line(color="grey",linetype = "dotted"),
        axis.line = element_line(),plot.margin = unit(c(4,2,4,2),units = "points"),
        axis.text.x = element_text(angle=35))

# generate panel B-specific data
figS5b.data = do.call(rbind,lapply(c("CRH","DRH"),function(mdl){
  selected.rows = plot.data$model%in%c(mdl,"EBR")&plot.data$LT%in%c(0.01,0.1,1)&
    plot.data$n==1250&plot.data$null_edit_rate==plot.data$LT&plot.data$null_n==1250
  df = plot.data[selected.rows,]
  df$pseudo_facet = mdl
  df$label = rate.label[as.character(df$LT)]
  df$null_N = factor(df$null_N,levels = null_N.range)
  df$N = factor(df$N,levels=null_N.range)
  df = df[df$LT%in%c(0.01,0.1,1),]
  return(df)
}))
# make panel A
figS5b = ggplot(figS5b.data)+
  geom_point(mapping=aes(x=null_N,y=power,color=hetero,shape=hetero))+
  geom_line(mapping=aes(x=null_N,y=power,color=hetero,group=hetero))+
  geom_errorbar(mapping=aes(x=null_N,ymin=lower,ymax=upper,color=hetero,group=hetero),width=0.2)+
  geom_vline(xintercept=2,color="black",linetype="dashed")+
  geom_hline(yintercept = 0.05,linetype="dashed",color="black")+
  facet_grid(rows=vars(pseudo_facet),cols=vars(label),labeller = label_parsed)+
  scale_x_discrete(labels=function(x){
    sapply(x,function(xx){null_N.label[which(xx==null_N.range)]})})+
  scale_color_manual(name = "Strength of rate\nheterogeneity",values = hetero.color,drop=FALSE)+
  scale_shape_manual(name = "Strength of rate\nheterogeneity",values = hetero.shape,drop=FALSE)+
  xlab("Population size for generating null distribution")+ylab("Power")+
  theme(legend.position = "none",
        axis.title = element_text(size=10),
        axis.text = element_text(size=8),
        legend.text = element_text(size = 8),
        legend.title = element_text(size=10),
        panel.background = element_blank(),
        panel.spacing.x = unit(12,units = "points"),
        strip.background.x = element_rect(fill="transparent",color="black"),
        strip.text.x = element_text(size=8,face = "bold",margin = margin(1,0,1,0)),
        strip.background.y = element_blank(),
        strip.text.y = element_blank(),
        panel.grid = element_line(color="grey",linetype = "dotted"),
        axis.line = element_line(),plot.margin = unit(c(4,2,4,2),units = "points"))

# aggregate panels
figS5ab= ggarrange(plotlist=list(figS5a,figS5b),labels = "AUTO",
                    ncol = 1,nrow = 2,align = "hv",
                    common.legend = TRUE,legend = "bottom",
                    legend.grob = get_legend(dummy.plot,position = "bottom"))
# arrange figure in the master canvas and add annotations
figS5 = ggplot()+
  annotation_custom(grob = cowplot::as_grob(figS5ab),xmin = 0,xmax = 180,ymin=0,ymax=185)+
  annotate(geom="tile",x=rep(182,4),y=c(48,81,130,164),height=31,width=4,fill=model.color[c(3,2,3,2)])+
  annotate(geom="text",x=rep(182,4),y=c(48,81,130,164),label=names(model.color)[c(3,2,3,2)],
           fontface="bold",color="white",size=3,angle=270)+
  coord_cartesian(xlim = c(-1,186),ylim = c(-1,186),expand = FALSE)+
  theme(panel.background = element_blank(),
        axis.line = element_blank(),axis.ticks = element_blank(),
        axis.text = element_blank(),axis.title = element_blank(),
        legend.position = "none",plot.margin = unit(c(0,0,0,0),units = "points"))

# save to file
ggsave(plot = figS5, path = "Figures_and_Tables/",filename = "Figure_S5.png",device = "png",
       height = 185,width = 185,scale = 1,units = "mm",dpi = 300)
ggsave(plot = figS5, path = "Figures_and_Tables/",filename = "Figure_S5.pdf",device = "pdf",
       height = 185,width = 185,scale = 1,units = "mm",colormodel="cmyk")
```
