## Supplementary Data and Scripts for "Detecting branching rate heterogeneity with tree balance statistics in lineage tracing trees": Fig_S6_Variability_of_J1_over_population_size.R

```
#
# load packages
library(ggplot2)
library(ggpubr)
source("Scripts/Fig_Color_Palettes.R")

# structure required data
empirical.nulls = readRDS("Simulated_data/stats/J1_Sackin_nulls.RDS")
null_edit_rate.range = c(0.01,0.1,1)
null_N.range = c(1250,6250,31250,1e6)
null_N.label = unname(sapply(c("1250","6250","31250","10^6"),function(x){parse(text=x)}))
params2use = which(
  empirical.nulls$meta$N%in%null_N.range&
    empirical.nulls$meta$n==1250&
    empirical.nulls$meta$type=="LTT"&
    empirical.nulls$meta$edit_rate%in%null_edit_rate.range)
plot.data = do.call(rbind,lapply(params2use,function(i){
  cbind(empirical.nulls$meta[i,],data.frame(J1 = empirical.nulls$null[[i]]$J1))
}))

# label for each editing rate
rate.label = c("0.01"="mu[edit]==0.01",
               "0.05"="mu[edit]==0.05",
               "0.1"="mu[edit]==0.10",
               "0.5"="mu[edit]==0.50",
               "1"="mu[edit]==1.00")
plot.data$label = rate.label[as.character(plot.data$edit_rate)]
plot.data$N = factor(plot.data$N,levels = null_N.range)

# make figure
figS6 = ggplot(plot.data)+
  geom_violin(mapping=aes(x=N,y=J1,group=N),fill="black",color="black")+
  facet_wrap(facets = vars(label),labeller = label_parsed,nrow = 1,ncol = 3)+
  xlab("Population size")+ylab(parse(text="J^1"))+
  scale_x_discrete(labels=function(x){
    sapply(x,function(xx){null_N.label[which(xx==null_N.range)]})})+
  theme(legend.position = "none",
        axis.title = element_text(size=10),
        axis.text = element_text(size=8),
        legend.text = element_text(size = 8),
        legend.title = element_text(size=10),
        panel.background = element_blank(),
        panel.spacing.x = unit(12,units = "points"),
        strip.background = element_rect(fill="transparent",color="black"),
        strip.text = element_text(size=8,face = "bold",margin = margin(1,0,1,0)),
        panel.grid = element_line(color="grey",linetype = "dotted"),
        axis.line = element_line(),plot.margin = unit(c(10,10,10,10),units = "points"))

# save to file
ggsave(plot = figS6, path = "Figures_and_Tables/",filename = "Figure_S6.png",device = "png",
       height = 75,width = 185,scale = 1,units = "mm",dpi = 300)
ggsave(plot = figS6, path = "Figures_and_Tables/",filename = "Figure_S6.pdf",device = "pdf",
       height = 75,width = 185,scale = 1,units = "mm",colormodel="cmyk")
```
