## Supplementary Data and Scripts for "Detecting branching rate heterogeneity with tree balance statistics in lineage tracing trees": infer_LT_tree_from_CM.py

```
# infer_LT_tree_from_CM.py
import sys
import argparse
import numpy as np
import pandas as pd
import cassiopeia as cas
import pickle

def save_object(obj, filename, mode):
    outp = open(filename, mode) # Overwrites any existing file.
    pickle.dump(obj, outp, pickle.HIGHEST_PROTOCOL)
    outp.close()

def pickle_loader(filename):
    """ Deserialize a file of pickled objects. """
    with open(filename, "rb") as f:
        while True:
            try:
                yield pickle.load(f)
            except EOFError:
                break

def main():
    # setting up parser
    parser = argparse.ArgumentParser()
    parser.add_argument('-o', '--output', required=True,help="Full directory of output files")
    parser.add_argument('-i', '--infile', required=True,help="File that contains the character matrix")
    parser.add_argument('--test',action='store_true',help="Test parser by printing parsed arguments")
    # parse arguments
    args = parser.parse_args()
    # print parsed arguments and exit if test mode is enabled
    if args.test:
        print(args)
        sys.exit(0)
    
    # read trees from Newick files:
    f = open(args.infile,mode="r")
    CM_text = pd.read_csv(f,sep="\t",header=0)
    f.close()
    CM_text = CM_text.replace("-",-1)
    CM_text = CM_text.set_index('cellBC')
    # then simulate lineage tracing trees
    reconstructed_tree = cas.data.CassiopeiaTree(
        character_matrix = CM_text,
        missing_state_indicator = -1
    )
    maxcut_solver = cas.solver.MaxCutSolver()
    maxcut_solver.solve(reconstructed_tree)
    f = open(args.output,"w")
    f.write(reconstructed_tree.get_newick(record_branch_lengths=True)+"\n")
    f.close()
    save_object(reconstructed_tree,args.output+".pkl",'wb')
    
if __name__ == "__main__":
    main()
```
