## Supplementary Data and Scripts for "Detecting branching rate heterogeneity with tree balance statistics in lineage tracing trees": simulate_genealogical_trees.py

```
# simulate_genealogical_trees.py
import sys
import argparse
import numpy as np
import pandas as pd
import cassiopeia as cas

def main():
    # setting up parser
    parser = argparse.ArgumentParser()
    parser.add_argument('-o', '--output', required=True,help="Full directory + prefix of output files")
    parser.add_argument('-N', '--pop_size', type=int,default=6250,help="Population size of the true trees")
    parser.add_argument('-n', '--sample_size', type=int,nargs='*',help="Tree size after subsampling")
    parser.add_argument('-t', '--tree',help="File that contains true trees as an alternative option; overides other simulator parameters")
    parser.add_argument('-i', '--replicate',type=int,default=1,help="Number of replicates")
    parser.add_argument('-b', '--birth',type=float,default=1.0,help="Expected time until a birth event at root")
    parser.add_argument('-u', '--mutation',type=float,nargs=2,help="Probability of a mutation (and it being beneficial) at birth event")
    parser.add_argument('-s', '--fitness',type=float,nargs=2,help="Fitness effect of beneficial and deleterious mutations")
    parser.add_argument('-d', '--death',type=float,help="Expected time until a death event")
    parser.add_argument('--test',action='store_true',help="Test parser by printing parsed arguments")
    # parse arguments
    args = parser.parse_args()
    # print parsed arguments and exit if test mode is enabled
    if args.test:
        print(args)
        sys.exit(0)
    
    # check if trees are supplied as Newick files:
    if args.tree is not None:
        f = open(args.tree,mode="r")
        newick_text = f.read()
        newick_text = newick_text.replace("\n","")
        tree_texts = [tt+";" for tt in newick_text.split(";") if tt]
        ground_truth_tree = [cas.data.CassiopeiaTree(tree=tree_text) for tree_text in tree_texts]
        f.close()
        
    # otherwise parse tree model
    else:
        # parse death rate (as expected time until death)
        if args.death is None:
            d_func = lambda:np.inf
        else:
            d_func = lambda:np.random.exponential(args.death)
        # parse mutation rate and fitness effect
        if args.mutation is None:
            mu_func = None
            s_func = None
        else:
            mu_func = lambda: 1 if np.random.uniform() < args.mutation[0] else 0
            if args.fitness is None:
                s_func = lambda: 0
            else:
                s_func = lambda: args.fitness[0] if np.random.uniform() < args.mutation[1] else args.fitness[1]
        # and build simulator
        bd_sim = cas.sim.BirthDeathFitnessSimulator(
            birth_waiting_distribution = lambda scale: np.random.exponential(scale),
            initial_birth_scale = args.birth,
            death_waiting_distribution = d_func,
            mutation_distribution = mu_func,
            fitness_distribution = s_func,
            num_extant = args.pop_size
        )
        # then simulate trees
        ground_truth_tree = [bd_sim.simulate_tree() for i in range(0,args.replicate)]
        print('{} full trees with {} tips were simulated.\n'.format(len(ground_truth_tree),args.pop_size))
        
        # save full genealogical trees
        f = open(args.output+'_N{}_n{}.nwk'.format(args.pop_size,args.pop_size),"w")
        for this_tree in ground_truth_tree:
            f.write(this_tree.get_newick(record_branch_lengths=True)+"\n")
        f.close()
    
    # subsample trees
    if args.sample_size is not None:
        for this_size in args.sample_size:
            sub_sim = cas.sim.UniformLeafSubsampler(number_of_leaves=this_size)
            subsample_tree = [sub_sim.subsample_leaves(tree=tree) for tree in ground_truth_tree]
            # save subsampled trees
            f = open(args.output+'_N{}_n{}.nwk'.format(args.pop_size,this_size),"w")
            for this_tree in subsample_tree:
                f.write(this_tree.get_newick(record_branch_lengths=True)+"\n")
            f.close()
            print('{} subtrees with {} tips sampled.\n'.format(len(subsample_tree),this_size))
    # print summary messages
    print("Simulation finished!\n")
    
if __name__ == "__main__":
    main()
```
