## Supplementary Data and Scripts for "Detecting branching rate heterogeneity with tree balance statistics in lineage tracing trees": simulate_lineage_tracing_trees.py

```
# simulate_genealogical_trees.py
import sys
import argparse
import numpy as np
import pandas as pd
import cassiopeia as cas
import pickle

def save_object(obj, filename, mode):
    outp = open(filename, mode) # Overwrites any existing file.
    pickle.dump(obj, outp, pickle.HIGHEST_PROTOCOL)
    outp.close()

def pickle_loader(filename):
    """ Deserialize a file of pickled objects. """
    with open(filename, "rb") as f:
        while True:
            try:
                yield pickle.load(f)
            except EOFError:
                break

def main():
    # setting up parser
    parser = argparse.ArgumentParser()
    parser.add_argument('-o', '--output', required=True,help="Full directory of output files")
    parser.add_argument('-t', '--tree', required=True,help="File that contains true genealogical trees")
    parser.add_argument('-n', '--site',type=int,default=5,help="Number of editable sites per cassette")
    parser.add_argument('-c', '--cassette',type=int,default=10,help="Number of cassettes")
    parser.add_argument('-u', '--mutation',type=float,default=0.01,help="Rate of mutation (editing)")
    parser.add_argument('-k', '--state',type=int,default=50,help="Number of states")
    parser.add_argument('-s', '--s_coef',type=float,default=1e-5,help="Exponent scaling coefficient of state distribution")
    parser.add_argument('--test',action='store_true',help="Test parser by printing parsed arguments")
    # parse arguments
    args = parser.parse_args()
    # print parsed arguments and exit if test mode is enabled
    if args.test:
        print(args)
        sys.exit(0)
    
    # read trees from Newick files:
    f = open(args.tree,mode="r")
    newick_text = f.read()
    f.close()
    newick_text = newick_text.replace("\n","")
    tree_texts = [tt+";" for tt in newick_text.split(";") if tt]
    ground_truth_tree = [cas.data.CassiopeiaTree(tree=tree_text) for tree_text in tree_texts]
    
    # and build simulator
    lt_sim = cas.sim.Cas9LineageTracingDataSimulator(
        number_of_cassettes = args.cassette,
        size_of_cassette = args.site,
        mutation_rate = args.mutation,
        state_generating_distribution = lambda: np.random.exponential(args.s_coef),
        number_of_states = args.state,
        state_priors = None,
	collapse_sites_on_cassette=False
    )
    # then simulate lineage tracing trees
    np.random.seed(seed=None)
    maxcut_solver = cas.solver.MaxCutSolver()
    f = open(args.output,"w")
    f.close()
    for this_tree in ground_truth_tree:
        lt_sim.overlay_data(this_tree)
        save_object(this_tree,args.output+"_true.pkl",'ab')
        reconstructed_tree = cas.data.CassiopeiaTree(
            character_matrix = this_tree.character_matrix,
            missing_state_indicator = -1
        )
        maxcut_solver.solve(reconstructed_tree)
        save_object(reconstructed_tree,args.output+".pkl",'ab')
        f = open(args.output,"a")
        f.write(reconstructed_tree.get_newick(record_branch_lengths=True)+"\n")
        f.close()
        print('### lineage tracing tree simualted and written to file.\n')
    print('{} lineage tracing trees were simulated.\n'.format(len(ground_truth_tree)))
    # print summary messages
    print("Simulation finished!\n")
    
if __name__ == "__main__":
    main()
```
