## Supplementary Data and Scripts for "Detecting branching rate heterogeneity with tree balance statistics in lineage tracing trees": Jq_tree_balance.R

```
#' @title  Calculate J^q tree balance statistics
#'
#' @description  Calculate J^q tree balance statistics for a `phylo` class object
#'
#' @param phy a `phylo` class object containing the tree
#' @param weight the weight of each node and tip
#' @param q,nonrootdomfactor same as the original implementation of J^q
#' @param binary.only whether multifurcating nodes should be excluded; default to `FALSE`
#' @details The function `calculate_Jq_tree_balance_metric` is a copy of the original J^q implementation that uses a `phylo` class object as the input. 
#' @return A numeric value of the J^q statistic
#' @export
#' @rdname calculate_Jq_tree_balance_metric
calculate_Jq_tree_balance_metric = 
  function(phy,weight=c(rep(1,Ntip(phy)),rep(0,Nnode(phy))),
           q=1,nonrootdomfactor=FALSE,binary.only=FALSE){
  n = Ntip(phy)+Nnode(phy)
  if (n<=1) return(0)
  # initialize key variables
  Adj = split(phy$edge[,2],phy$edge[,1]) # adjacency list
  Cumul = get_clade_size_for_each_node(phy,weight = weight) # subtree sizes, including the root
  Star = Cumul - weight # subtree sizes, excluding the root
  eff_int_nodes = Ntip(phy)+(1:Nnode(phy)) # vector of internal nodes
  binary.idx = which(sapply(Adj[as.character(eff_int_nodes)],length)==2)
  if(binary.only&(length(binary.idx)<1)){
    warning("No binary node found. Returning 0.",immediate. = TRUE)
    return(0)
  }
  # non-root dominance factor:
  h_factor = (Star/Cumul)^as.numeric(nonrootdomfactor)
  # calculate J
  J = sapply(eff_int_nodes,function(i){
    if(Star[i]<1) return(0)
    eff_children = Cumul[Adj[[as.character(i)]]]>0
    K = sum(sapply(Adj[[as.character(i)]],function(j){
      p = Cumul[j]/Star[i]
      if(q == 1) {
        return(-p*log(p))
      } else {
        return(p^q)
      }
    })[eff_children])
    eff_children = sum(eff_children)
    # normalize the sum of balance scores
    if(q==1){
      return(Star[i] * K / log(eff_children))
    }else{
      return(Star[i] * (1 - K) * eff_children^(q - 1) / (eff_children^(q - 1) - 1))
    }
  })
  if(binary.only){
    J = sum(h_factor[eff_int_nodes[binary.idx]]*J[binary.idx],na.rm = TRUE)/sum(Star[eff_int_nodes[binary.idx]])
  }else{
    J = sum(h_factor[eff_int_nodes]*J,na.rm = TRUE)/sum(Star[eff_int_nodes])
  }
  return(J)
}
    
#' @title  Calculate the Sackin index
#'
#' @description  Calculate the Sackin index for a `phylo` class object
#'
#' @param phy a `phylo` class object containing the tree to be tested
#' @param normalize whether the output value should be normalized by the tree size
#' @details The function `calculate_Sackin_index` calculates the Sackin index for a `phylo` class object with the option to normalize it by the tree size. 
#' @return A numeric value of the Sackin index
#' @export
#' @rdname calculate_Sackin_index
calculate_Sackin_index = function(phy,normalize=FALSE){
  N = Ntip(phy)
  phy$edge.length = rep(1,dim(phy$edge)[1])
  this.sackin = sum(node.depth.edgelength(phy)[1:N])
  if(normalize) this.sackin = (this.sackin-N)/(N*(N-1)/2-1)
  return(this.sackin)
}
```
