## Supplementary Data and Scripts for "Detecting branching rate heterogeneity with tree balance statistics in lineage tracing trees": phylo_ops.R

```
#
phyloapply = function(x,phy,func,
                      use.branch.length=FALSE,use.self=FALSE,postorder=TRUE){
  # reorder edges in the phylo object
  phy = reorder.phylo(phy,order = "postorder")
  # initialize result list for each tip and node
  res = lapply(1:(Ntip(phy)+Nnode(phy)),function(i){list()})
  if(length(x)>1){
    res[1:length(x)] = as.list(x)
  }else{
   res[[Ntip(phy)+1]] = x
  }
  # determine the order of application
  if(postorder){
    index.apply = split(1:dim(phy$edge)[1],factor(phy$edge[,1],levels=unique(phy$edge[,1])))
    nnidx = 1
  }else{
    index.apply = rev(1:dim(phy$edge)[1])
    nnidx = 2
  }
  # apply functions over the phylo object
  if(use.branch.length|use.self){
    for(nn in index.apply){
      this.arg = list(res[[unique(phy$edge[nn,nnidx])]],#value of itself
                      res[phy$edge[nn,3-nnidx]],#value of parent/children
                      phy$edge.length[nn])#branch length
      res[[unique(phy$edge[nn,nnidx])]] = 
        do.call(what = func,args = this.arg[c(use.self,TRUE,use.branch.length)])
    }
  }else{
    for(nn in index.apply){
      res[[unique(phy$edge[nn,nnidx])]] = 
        do.call(what = func,args = res[phy$edge[nn,3-nnidx]])
    }
  }
  return(res)
}

#
get_clade_size_for_each_node = 
  function(phy,weight=c(rep(1,Ntip(phy)),rep(0,Nnode(phy)))){
  res = phyloapply(x = weight,phy = phy,
                   func = function(x,y){do.call("sum",c(as.list(x),y))},
                   use.branch.length = FALSE,use.self = TRUE,postorder = TRUE)
  return(do.call(c,res))
}
```
