## Supplementary Data and Scripts for "Detecting branching rate heterogeneity with tree balance statistics in lineage tracing trees": test_of_constant_rate.R

```
#' @title  Test if the branching rate of a tree is constant through user-specified tree statistics
#'
#' @description  A wrapper function to test the hypothesis of a constant branching rate for a tree through tree statistics
#'
#' @param phy a `phylo` class object containing the tree to be tested
#' @param func,params the function and its arguments to calculate the tree statistics, see `Details` 
#' @param null.phy the baseline trees under the null model, see `Details`
#' @param nrep the number of replicates for baseline trees
#' @param ncore the number of cores to use in parallel computation
#' @param alternative the alternative hypothesis
#' @details The function `test_of_constant_branching_rate_by_summary_statistic` tests the hypothesis that the focal tree emerges under a constant rate birth-death model.
#' @details The `func` argument can be a function or the name of that function, or a list of function and its arguments (and the `params` argument will be ignored).
#' @details The `null.phy` argument can be a numeric vector of the empirical null distribution or a list of `phylo` class objects, or a list of function and its arguments to generate the previous two types.
#' @return A list of four elements containing the observed statistic ($statistic), null distribution ($base), p-value of the null hypothesis ($p.value), and the alternative hypothesis ($alternative)
#' @export
#' @rdname test_of_constant_rate_by_summary_statistic
test_of_constant_branching_rate_by_summary_statistic = 
  function(phy,func="calculate_Jq_tree_balance_metric",params=list(),
           null.phy=NULL,nrep=1000,ncore=1,alternative="less"){
    this.stat = do.call(func,c(list(phy),params))
    n = Ntip(phy)
    if(is.null(null.phy)){
      cat("No baseline trees supplied, simulating from scratch...\n")
      null.phy = mclapply(1:nrep,function(i){
        diversitree::tree.yule(pars = c(1),max.taxa = n)
      },mc.cores = ncore)
    }
    if(!is.numeric(null.phy)){
      if(!is.null(null.phy$func)&!is.null(null.phy$params)){
        cat("Generating baseline from user-supplied function...\n")
        null.phy = mclapply(1:nrep,function(i){
          do.call(null.phy$func,null.phy$params)
        },mc.cores = ncore)
      }
    }
    #
    if(is.numeric(null.phy)){
      baseline.stat = null.phy
    }else{
      if(is.list(func)){
        params = func$params
        func = func$func
      }
      baseline.stat = do.call(c,mclapply(null.phy,function(base.tree){
        base.stat = do.call(func,c(list(base.tree),params))
        return(base.stat)
      },mc.cores = ncore))
    }
    #
    left.p = mean(sum(baseline.stat<this.stat),sum(baseline.stat<=this.stat))
    right.p = mean(sum(baseline.stat>this.stat),sum(baseline.stat>=this.stat))
    this.pval = c(less=left.p,greater=right.p,two.sided=min(left.p,right.p)*2)/nrep
    if(!alternative%in%names(this.pval)){
      warning("Invalid alternative hypothesis, enforcing to two-sided.",immediate. = TRUE)
      alternative="two.sided"
    }
    return(list(statistic=this.stat,base=baseline.stat,p.value=unname(this.pval[alternative]),alternative=alternative))
  }
```
