## Supplementary Data and Scripts for "Detecting branching rate heterogeneity with tree balance statistics in lineage tracing trees": reproduce_figures_and_tables.sh

```
#!/bin/bash
# Description: This script documents the examples codes to reproduce the figures and tables 
# Authors: Yingnan Gao, Alison F. Feder
# DOI: https://doi.org/10.1101/2024.06.27.601073

# The directory from which the following commands are run should be the same as
# that of README.txt
# To reproduce the figures, make sure that null_trees.tar.gz and test_trees.tar.gz
# are extracted:
tar -xvzf Simulated_data/null_trees.tar.gz -C Simulated_data/
tar -xvzf Simulated_data/test_trees.tar.gz -C Simulated_data/

# Then reproduce the figures through:
Rscript Scripts/Fig_1_Signal_of_branching_rate_heterogeneity.R
Rscript Scripts/Fig_2_Effect_of_lineage_tracing_on_tree_balance.R
Rscript Scripts/Fig_3_S4_Performance_under_LT.R
Rscript Scripts/Fig_4_S7_Testing_empirical_trees.R
Rscript Scripts/Fig_S1_Effect_of_tree_size_on_power.R
Rscript Scripts/Fig_S2_More_effects_of_LT_on_trees.R
Rscript Scripts/Fig_S3_More_effects_of_LT_on_tree_balance.R
Rscript Scripts/Fig_S5_Robustness_of_J1_against_misspecification.R
Rscript Scripts/Fig_S6_Variability_of_J1_over_population_size.R

# For tables, run the two analysis scripts by:
# Table S1
Rscript Scripts/Analysis_benchmarking_LT_estimation.R
# Table S2
Rscript Scripts/Analysis_test_EBR_in_empirical_data.R

# Alternatively, these scripts can be sourced in Rstudio with working directory 
# set to the same directory as README.txt.
```
