## Supplementary Data and Scripts for "Detecting branching rate heterogeneity with tree balance statistics in lineage tracing trees": reproduce_simulation.sh

```
#!/bin/bash
# Description: This script documents the examples codes to reproduce a small fraction
# of simulations in the manuscript titled 'Detecting branching rate heterogeneity with
# tree balance statistics in lineage tracing trees'. The simulations were run on the
# high performance computing clusters of UW Genome Sciences. This script only shows the
# core functional commands for the simulations, and only yields 10 out of 1000 replicates. 
# Authors: Yingnan Gao, Alison F. Feder
# DOI: https://doi.org/10.1101/2024.06.27.601073

# 0. Setting up directory and environments
# Scripts to reproduce the simulations are stored in separate folders by their language used.
# For reproduction, make a new directory and copy all the scripts under Scripts/Python and
# Scripts/Rscript to the new directory.
# Make sure the python software Cassiopeia (https://github.com/YosefLab/Cassiopeia) is
# available in your environment.
# Also make sure the following R package is installed:
# ape, diversitree, parallel
# From the newly created directory, run the commands in the following sections.

# 1. Simulating genealogical trees
# EBR trees can be simulated through the commands:
python simulate_genealogical_trees.py -N 6250 -n 50 250 1250 -i 10 -o EBR

# DRH trees can be simulated through the commands:
python simulate_genealogical_trees.py -N 6250 -n 50 250 1250  -u 0.007 0.100 -s 0.700 -0.078 -o DRH_S062 -i 10  
python simulate_genealogical_trees.py -N 6250 -n 50 250 1250  -u 0.029 0.100 -s 0.700 -0.078 -o DRH_S126 -i 10
python simulate_genealogical_trees.py -N 6250 -n 50 250 1250  -u 0.179 0.100 -s 0.700 -0.078 -o DRH_S312 -i 10
python simulate_genealogical_trees.py -N 6250 -n 50 250 1250  -u 0.714 0.100 -s 0.700 -0.078 -o DRH_S624 -i 10

# Unsampled CRH trees can be simulated through R::diversitree
Rscript simulate_genealogical_trees.R CRH_S01_N6250_n6250 1 10 6250 0.1
Rscript simulate_genealogical_trees.R CRH_S05_N6250_n6250 1 10 6250 0.5
Rscript simulate_genealogical_trees.R CRH_S1_N6250_n6250 1 10 6250 1
Rscript simulate_genealogical_trees.R CRH_S5_N6250_n6250 1 10 6250 5
Rscript simulate_genealogical_trees.R CRH_S10_N6250_n6250 1 10 6250 10
# Then sampled by running the commands:
python simulate_genealogical_trees.py -N 6250 -n 50 250 1250 -t CRH_S01_N6250_n6250.nwk -o CRH_S01
python simulate_genealogical_trees.py -N 6250 -n 50 250 1250 -t CRH_S05_N6250_n6250.nwk -o CRH_S05
python simulate_genealogical_trees.py -N 6250 -n 50 250 1250 -t CRH_S1_N6250_n6250.nwk -o CRH_S1
python simulate_genealogical_trees.py -N 6250 -n 50 250 1250 -t CRH_S5_N6250_n6250.nwk -o CRH_S5
python simulate_genealogical_trees.py -N 6250 -n 50 250 1250 -t CRH_S10_N6250_n6250.nwk -o CRH_S10

# We may store the unsampled trees in a separate folder as we do not use them:
mkdir full_trees_not_in_use
mv *N6250_n6250*.nwk full_trees_not_in_use

# 2. Simulating lineage tracing trees
# For a single Newick tree file, we may simulate lineage tracing trees with a certain
# editing rate by the command:
# python simulate_lineage_tracing_trees.py -t EBR_N6250_N50.nwk -o EBR_N6250_N50_3_10_001.tre -n 3 -c 10 -u 0.01

# To simulate lineage tracing trees in batch, we may use the R script that automates the
# call of the above command for each .nwk file in a specified directory (replace 1 with
# the number of cores you intend to use):
Rscript run_LT_simulation_for_each_nwk_file.R ./ nwk 1 0.01
Rscript run_LT_simulation_for_each_nwk_file.R ./ nwk 1 0.05
Rscript run_LT_simulation_for_each_nwk_file.R ./ nwk 1 0.10
Rscript run_LT_simulation_for_each_nwk_file.R ./ nwk 1 0.50
Rscript run_LT_simulation_for_each_nwk_file.R ./ nwk 1 1.00

# The python script simulate_lineage_tracing_trees.py generates intermediate files
# that contains the character matrices. We may store them for future use and clean
# up the folder for later simulations
mkdir pickle
mv *.pkl pickle
mkdir test_trees
mv *nwk test_trees
mv *.tre test_trees

# Check the directory. There should be no .nwk file left outside the subfolders.
ls ./

# 3. Simulating null EBR trees
# Simulation of null EBR trees is similar to that of EBR trees used as focal trees
# in tests. However, we tried more parameter combinations in these trees to test
# the impact of parameter misspecification. Make sure you have checked the folder
# that no .nwk file is left outside subfolders.

# First, we start with genealogical trees:
python simulate_genealogical_trees.py -N 1250 -n 50 250 -i 10 -o NULL
python simulate_genealogical_trees.py -N 6250 -n 50 250 1250 -i 10 -o NULL
python simulate_genealogical_trees.py -N 31250 -n 50 250 1250 -i 10 -o NULL
python simulate_genealogical_trees.py -N 1000000 -n 50 250 1250 -i 10 -o NULL
mv NULL_N6250_n6250.nwk NULL_N31250_n31250.nwk NULL_N1000000_n1000000.nwk full_trees_not_in_use
# Note that NULL_N1000000_n1000000.nwk can be huge, especially with more replicates.

# Then we simulate lineage tracing trees:
Rscript run_LT_simulation_for_each_nwk_file.R ./ nwk 1 0.01
Rscript run_LT_simulation_for_each_nwk_file.R ./ nwk 1 0.05
Rscript run_LT_simulation_for_each_nwk_file.R ./ nwk 1 0.10
Rscript run_LT_simulation_for_each_nwk_file.R ./ nwk 1 0.20
Rscript run_LT_simulation_for_each_nwk_file.R ./ nwk 1 0.30
Rscript run_LT_simulation_for_each_nwk_file.R ./ nwk 1 0.40
Rscript run_LT_simulation_for_each_nwk_file.R ./ nwk 1 0.50
Rscript run_LT_simulation_for_each_nwk_file.R ./ nwk 1 0.60
Rscript run_LT_simulation_for_each_nwk_file.R ./ nwk 1 0.70
Rscript run_LT_simulation_for_each_nwk_file.R ./ nwk 1 0.80
Rscript run_LT_simulation_for_each_nwk_file.R ./ nwk 1 0.90
Rscript run_LT_simulation_for_each_nwk_file.R ./ nwk 1 1.00

# After that we clean up the folder:
mv *.pkl pickle
mkdir null_trees
mv *nwk null_trees
mv *.tre null_trees

# The subfolder test_trees/ and null_trees/ should correspond to the subfolders
# extracted from test_trees.tar.gz and null_trees.tar.gz, with fewer replicates
```
