## Supplementary Data and Scripts for "Detecting branching rate heterogeneity with tree balance statistics in lineage tracing trees": calculate_editing_proportion.R

```
# calculate_editing_proportion
arg=commandArgs(TRUE)
indir =arg[1]
inpattern = arg[2]
outfile = arg[3]
barcode.size=as.numeric(arg[4])
#
infiles = list.files(indir,inpattern,full.names = TRUE)
instat = lapply(infiles,function(f){
  print(f)
  CM = read.table(f,sep=",",row.names = 1,header = FALSE)
  this.p = unname(apply(CM,1,function(x){
    n.all = length(x[1:barcode.size])
    if(is.numeric(x)){
      n.miss = sum(x[1:barcode.size]<0)
      n.none = sum(x[1:barcode.size]==0)
    }else{
      n.miss = sum(x[1:barcode.size]=="-")
      n.none = sum(x[1:barcode.size]=="0")
    }
    return(data.frame(n.miss=n.miss,n.none=n.none,n.all=n.all,
                      pp=(n.all-n.none-n.miss)/(n.all-n.miss)))
  },simplify = FALSE))
  this.meta = unname(sapply(rownames(CM),function(xx){
    rev(strsplit(xx,split = "_")[[1]])[2]
  }))
  return(lapply(split(x=this.p,f=this.meta),function(x){
    do.call(rbind,x)
  }))
})
names(instat) = infiles
saveRDS(object = instat,file = outfile)
```
