## Supplementary Data and Scripts for "Detecting branching rate heterogeneity with tree balance statistics in lineage tracing trees": infer_for_each_CM_file.R

```
library(parallel)
args = commandArgs(TRUE)
CM.files = list.files(path = args[1],pattern = args[2],full.names = TRUE)
commands = sapply(CM.files,function(f){
  newfile = gsub(pattern = args[2],replacement = "_MaxCut.tre",x = f)
  this.command = sprintf("python infer_LT_tree_from_CM.py -i %s -o %s",f,newfile)
  return(this.command)
})

out = mclapply(commands,function(x){
  cat(sprintf("Running command:%s\n",x))
  system(command = x,intern = TRUE)
},mc.cores = as.numeric(args[3])-1)
```
