## Supplementary Data and Scripts for "Detecting branching rate heterogeneity with tree balance statistics in lineage tracing trees": run_LT_simulation_for_each_nwk_file.R

```
library(parallel)
args = commandArgs(TRUE)
nwk.files = list.files(path = args[1],pattern = args[2],full.names = TRUE)
edit_rate = as.numeric(args[4])
commands = sapply(nwk.files,function(f){
  newfile = gsub(pattern = args[2],replacement = sprintf("_3_10_%03d.tre",round(edit_rate*100)),x = f)
  this.command = sprintf("python simulate_lineage_tracing_trees.py -t %s -o %s -n 3 -c 10 -u %.2f",f,newfile,edit_rate)
  return(this.command)
})

out = mclapply(commands,function(x){
  cat(sprintf("Running command:%s\n",x))
  system(command = x,intern = TRUE)
},mc.cores = max(1,as.numeric(args[3])-1))
```
