## Supplementary Data and Scripts for "Detecting branching rate heterogeneity with tree balance statistics in lineage tracing trees": simulate_genealogical_trees.R

```
#
library(ape)
library(diversitree)
library(parallel)
#
arg = commandArgs(TRUE)
savename = arg[1] # output file name with suffix
numCores = as.numeric(arg[2]) # number of cores to use
N = as.numeric(arg[3]) # number of replicates
Ntip = as.numeric(arg[4]) # number of tips in the tree
s = as.numeric(arg[5]) # strength of rate heterogeneity
#
trees = mclapply(1:N,function(i){
  #cat(sprintf("Simulation\t\t%d\t\t\r",i))
  exp.x = function(x,s,c){c*exp(s*x)}
  lambda <- function(x) exp.x(x, s, 1)
  mu = function(x) constant.x(x, 0)
  char <- make.brownian.with.drift(0, 0.1)
  phy <- tree.quasse(c(lambda, mu, char), max.taxa=Ntip, x0=0, single.lineage=FALSE, verbose=FALSE)
  return(phy)
},mc.cores = numCores)
cat(sprintf("\nSimluation completed\n"))
saveRDS(trees,paste0(savename,".RDS"))
write.tree(trees,paste0(savename,".nwk"))
#
```
