## Supplementary figures and images for "Detecting branching rate heterogeneity with tree balance statistics in lineage tracing trees"

### Figure_1.pdf

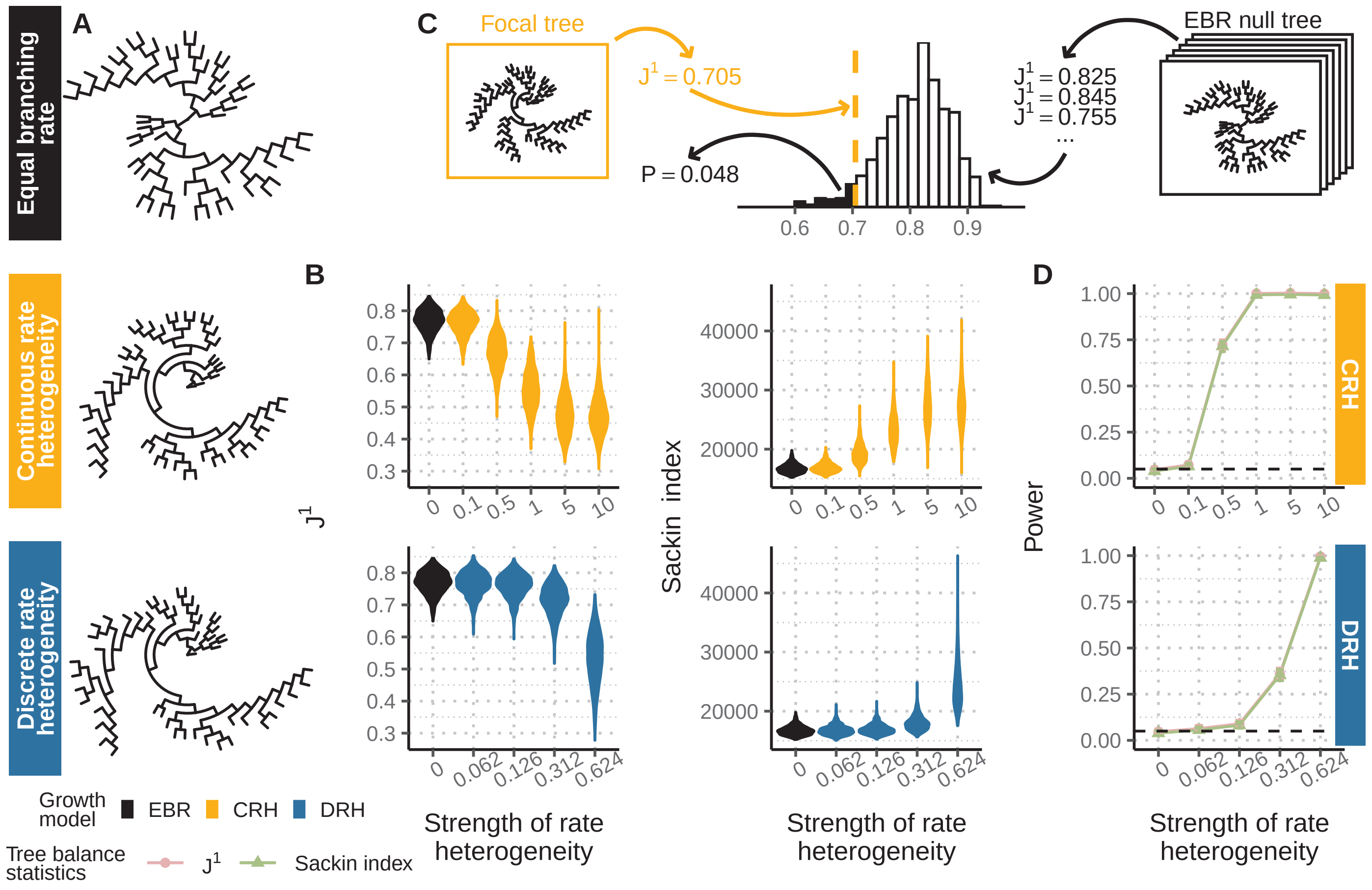

### Figure_1.png

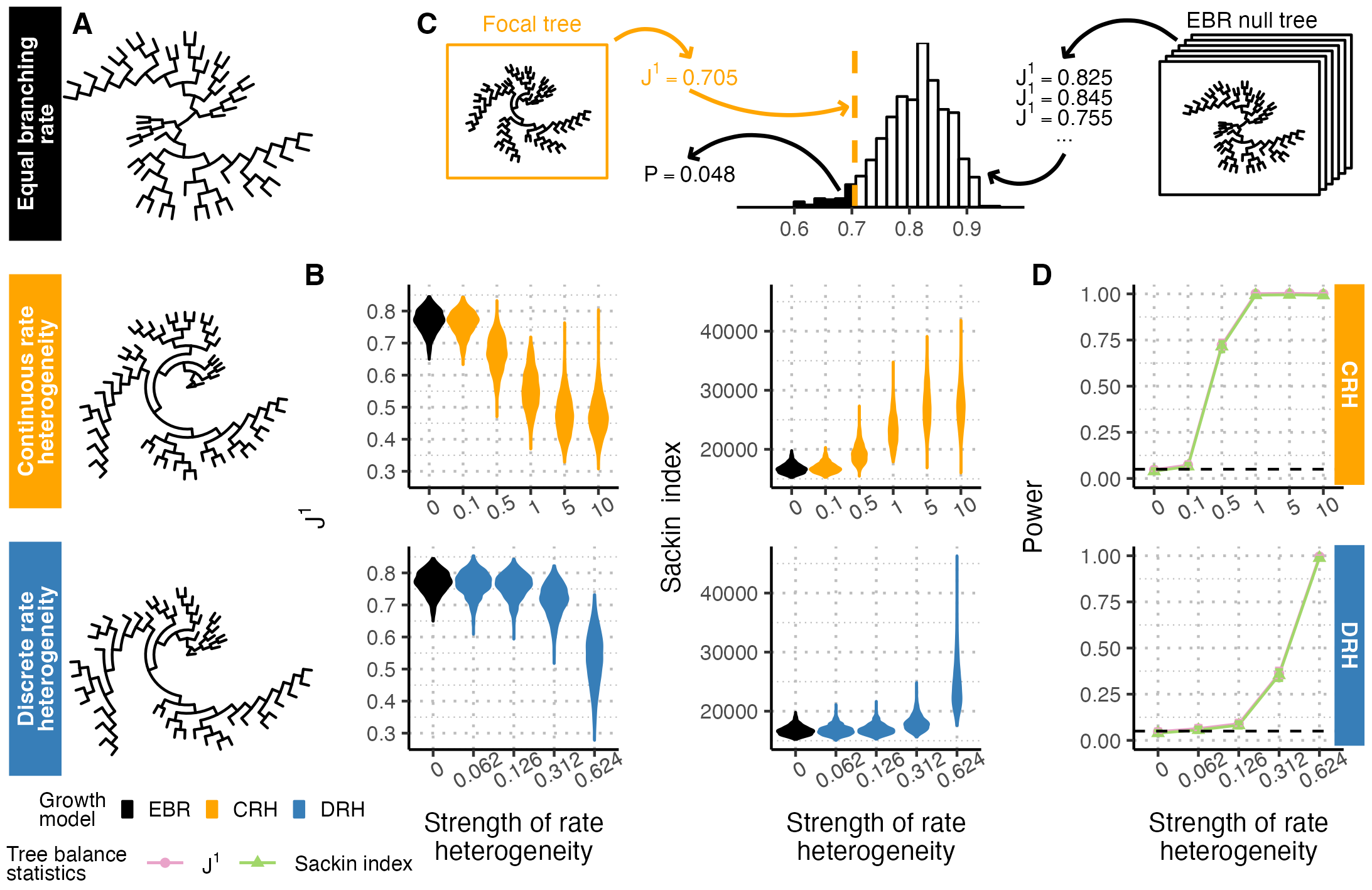

### Figure_2.pdf

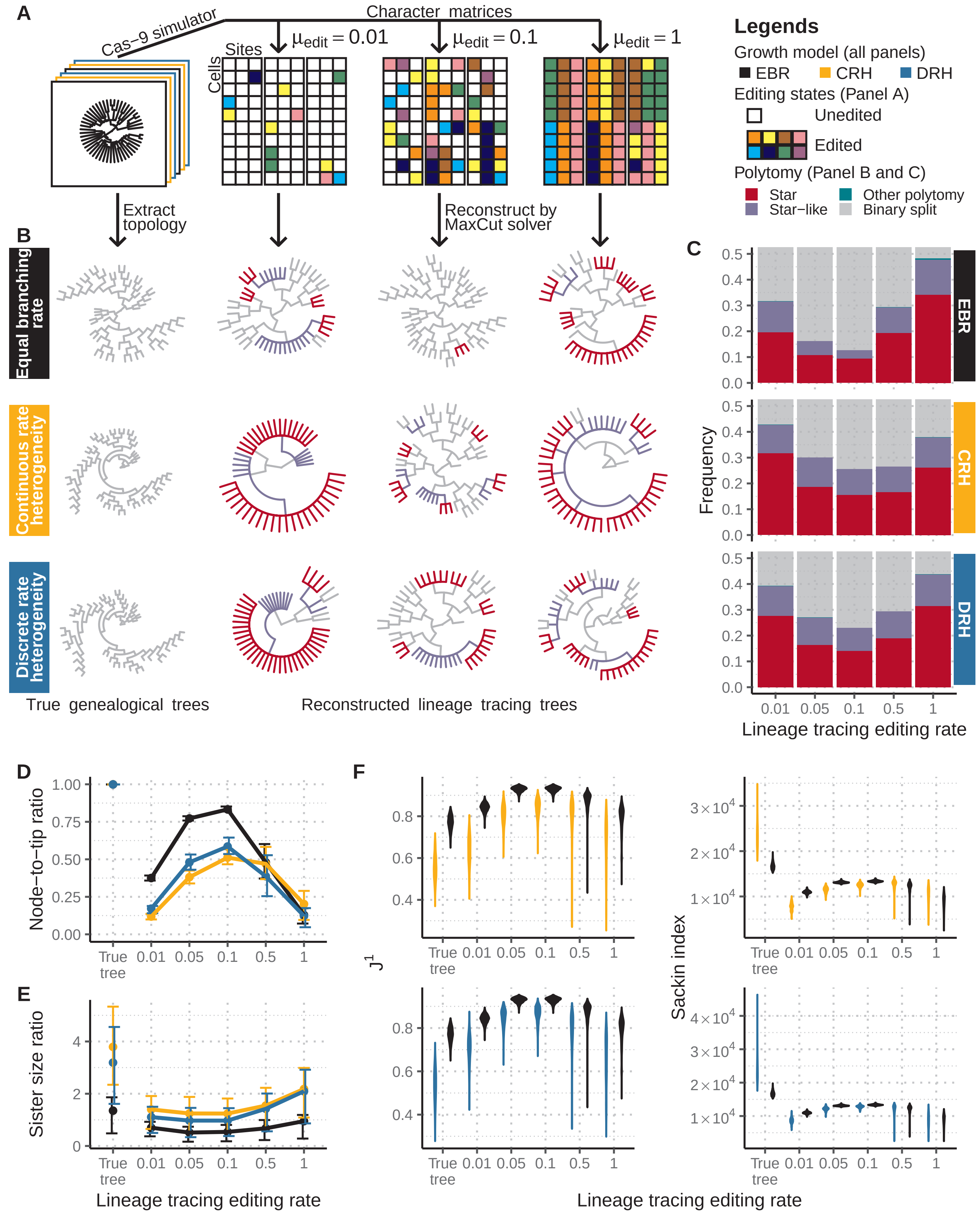

### Figure_2.png

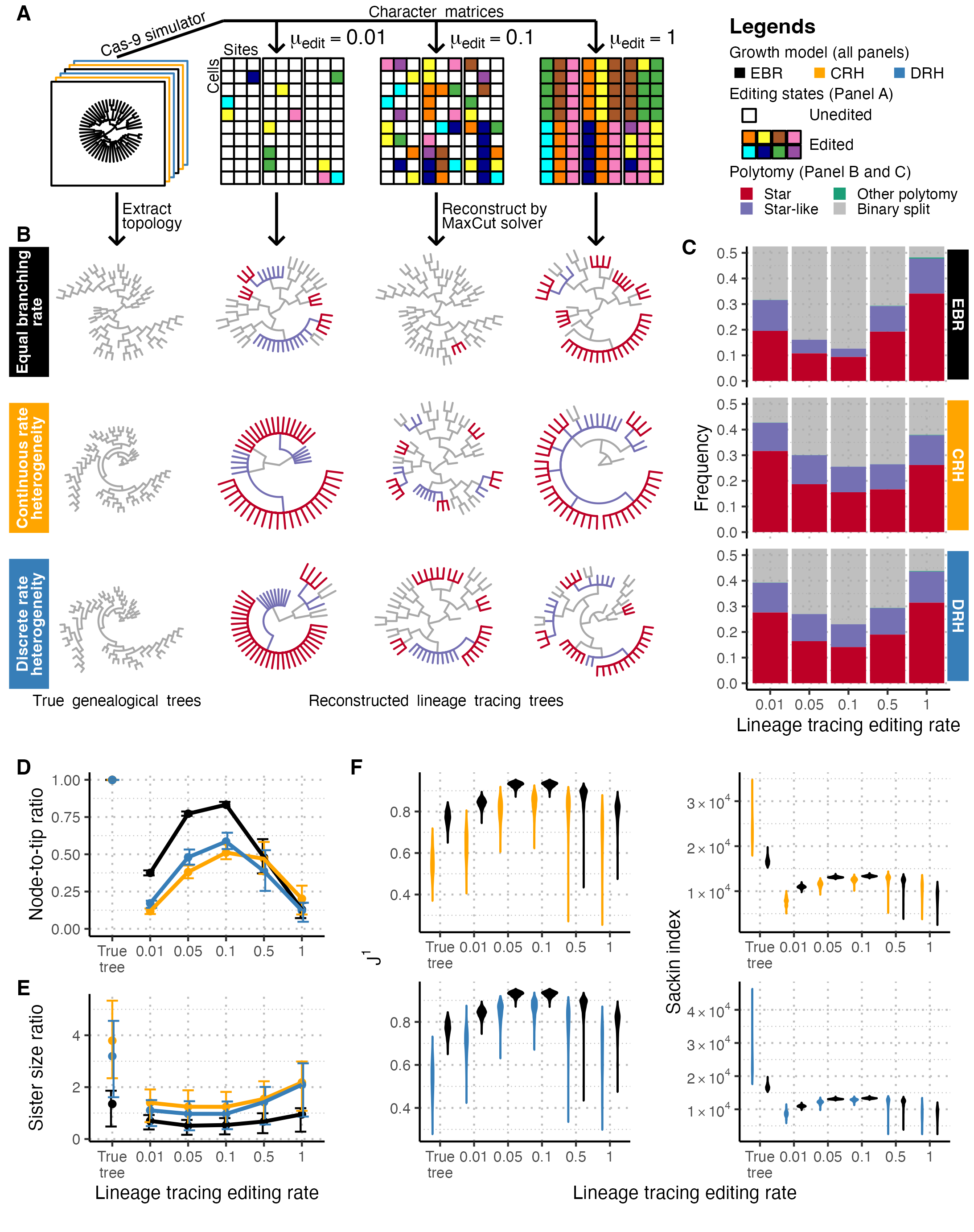

### Figure_3.pdf

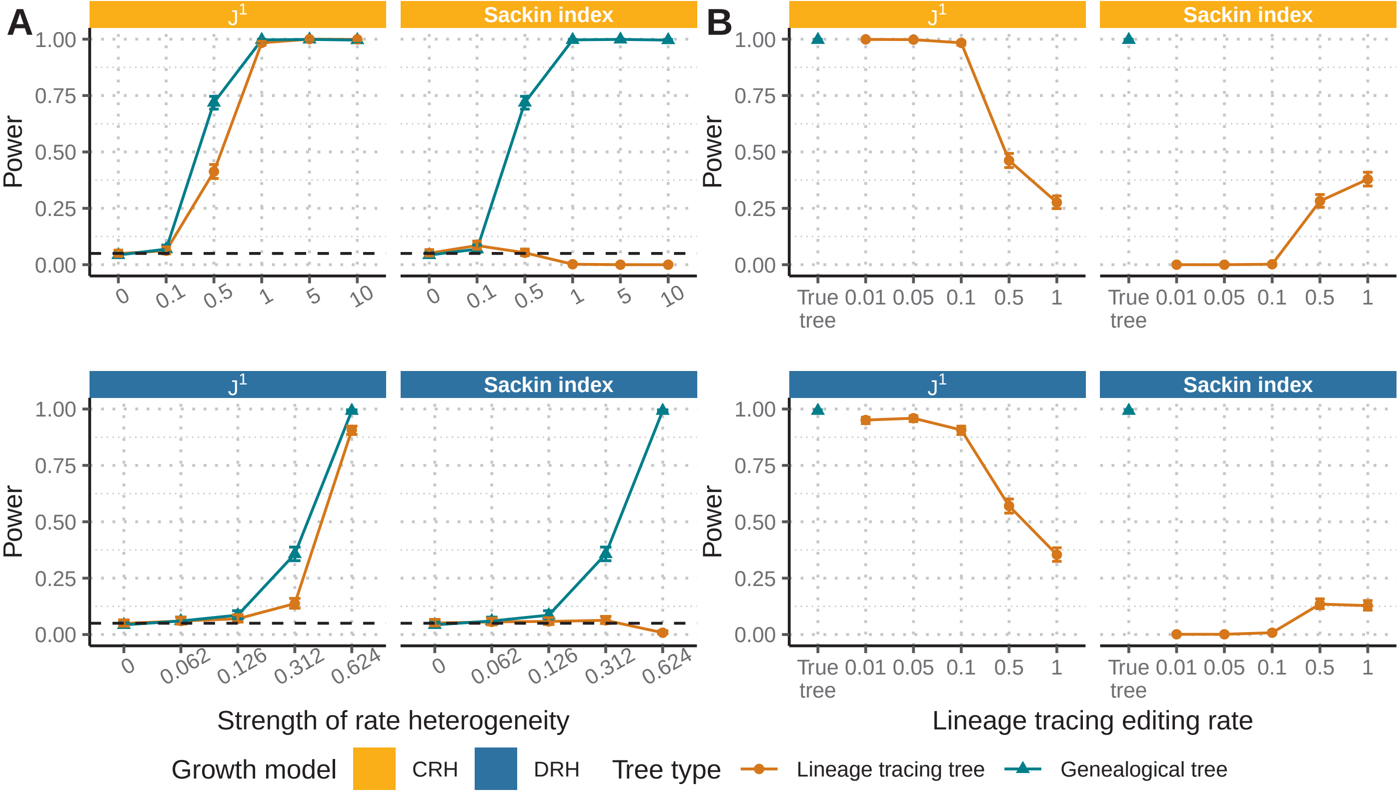

### Figure_3.png

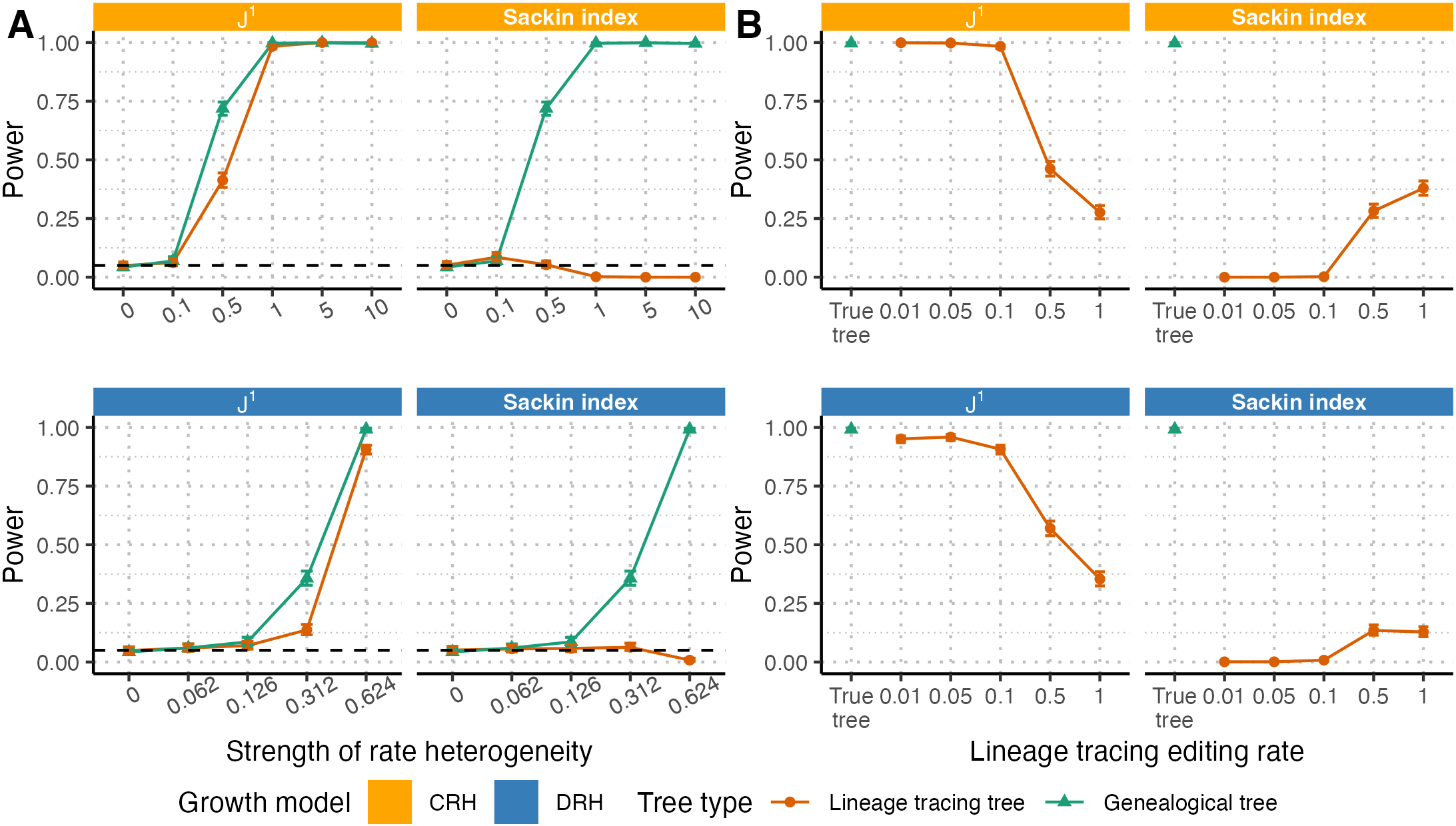

### Figure_4.pdf

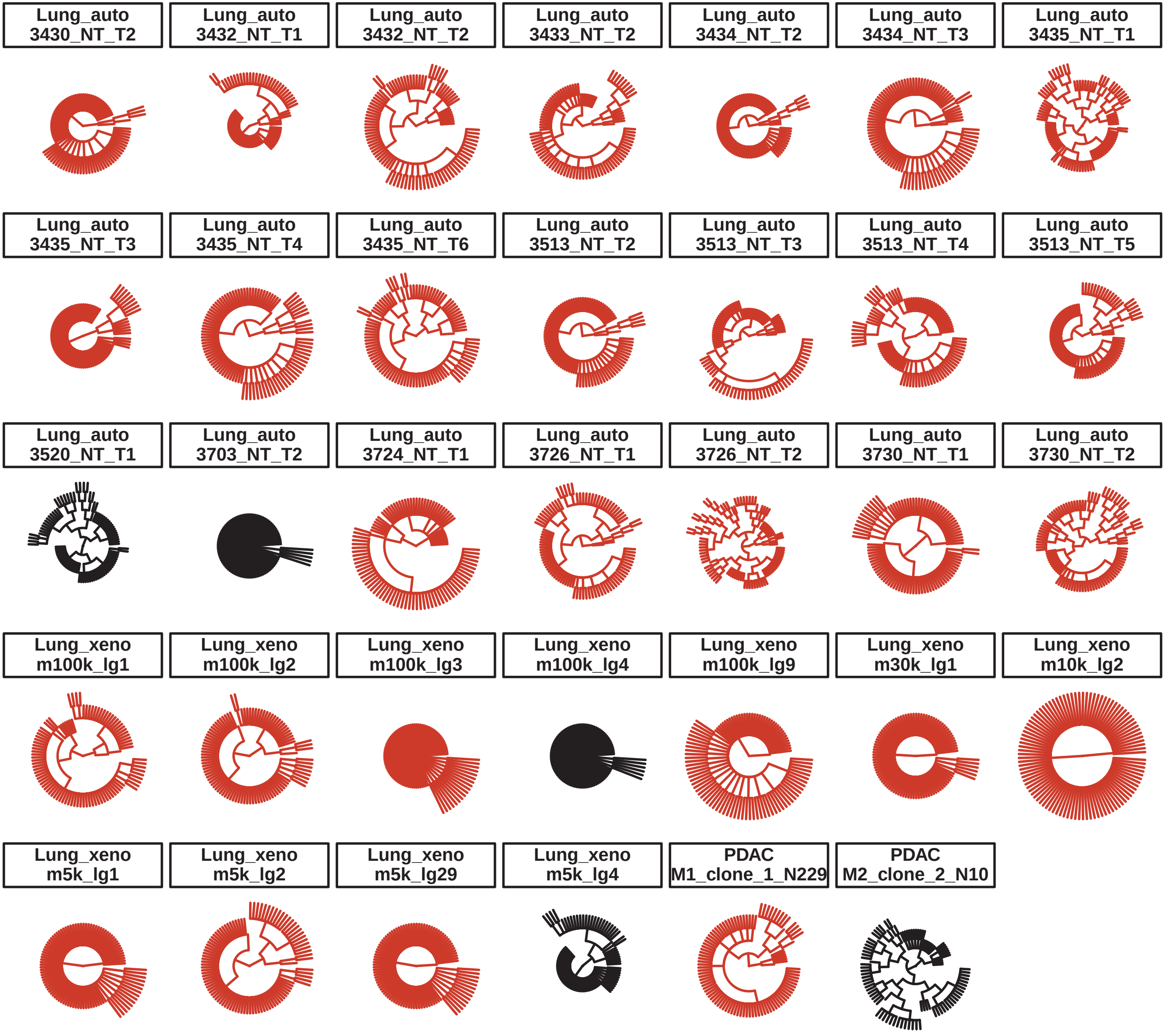

### Figure_4.png

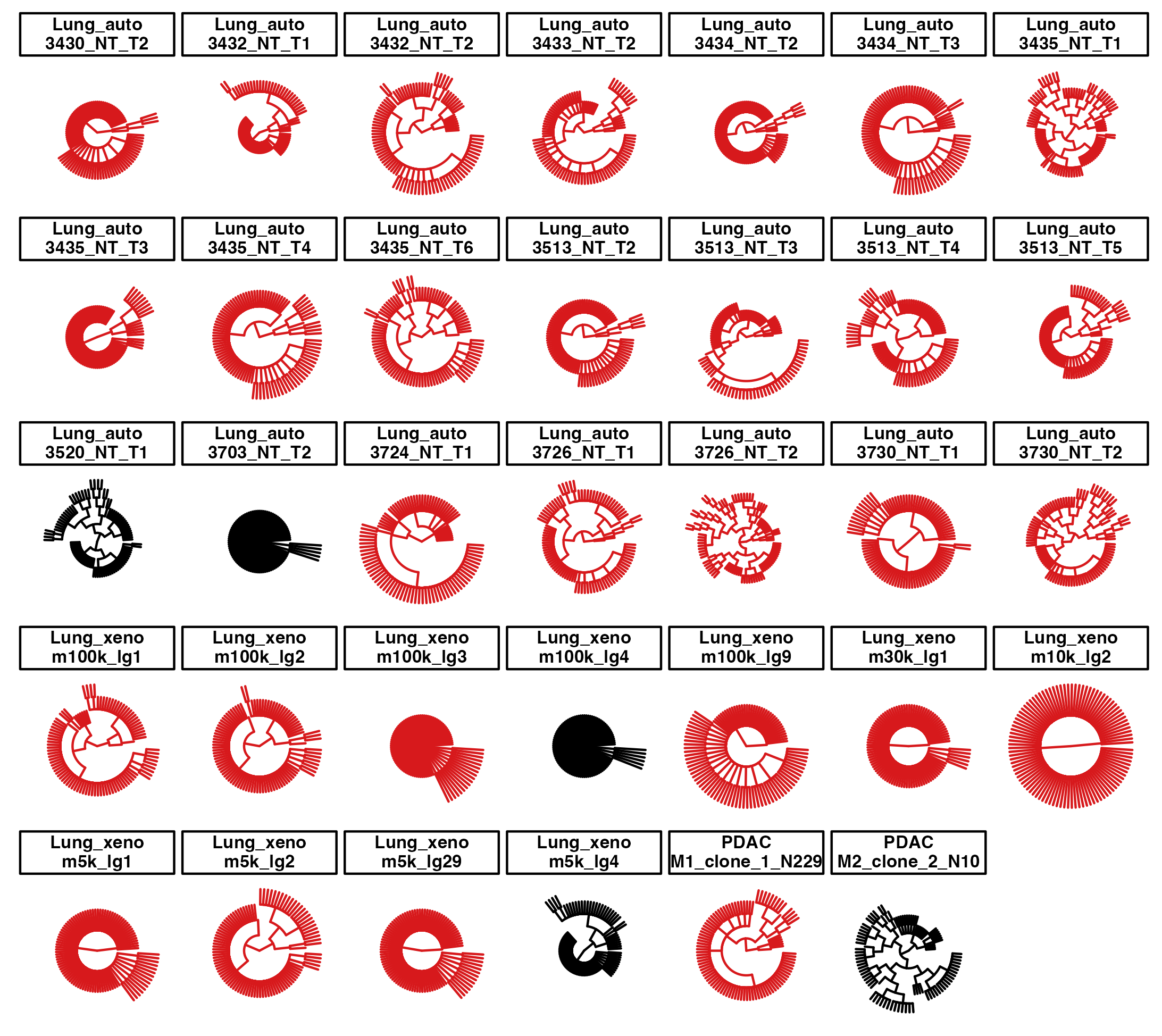

### Figure_S1.pdf

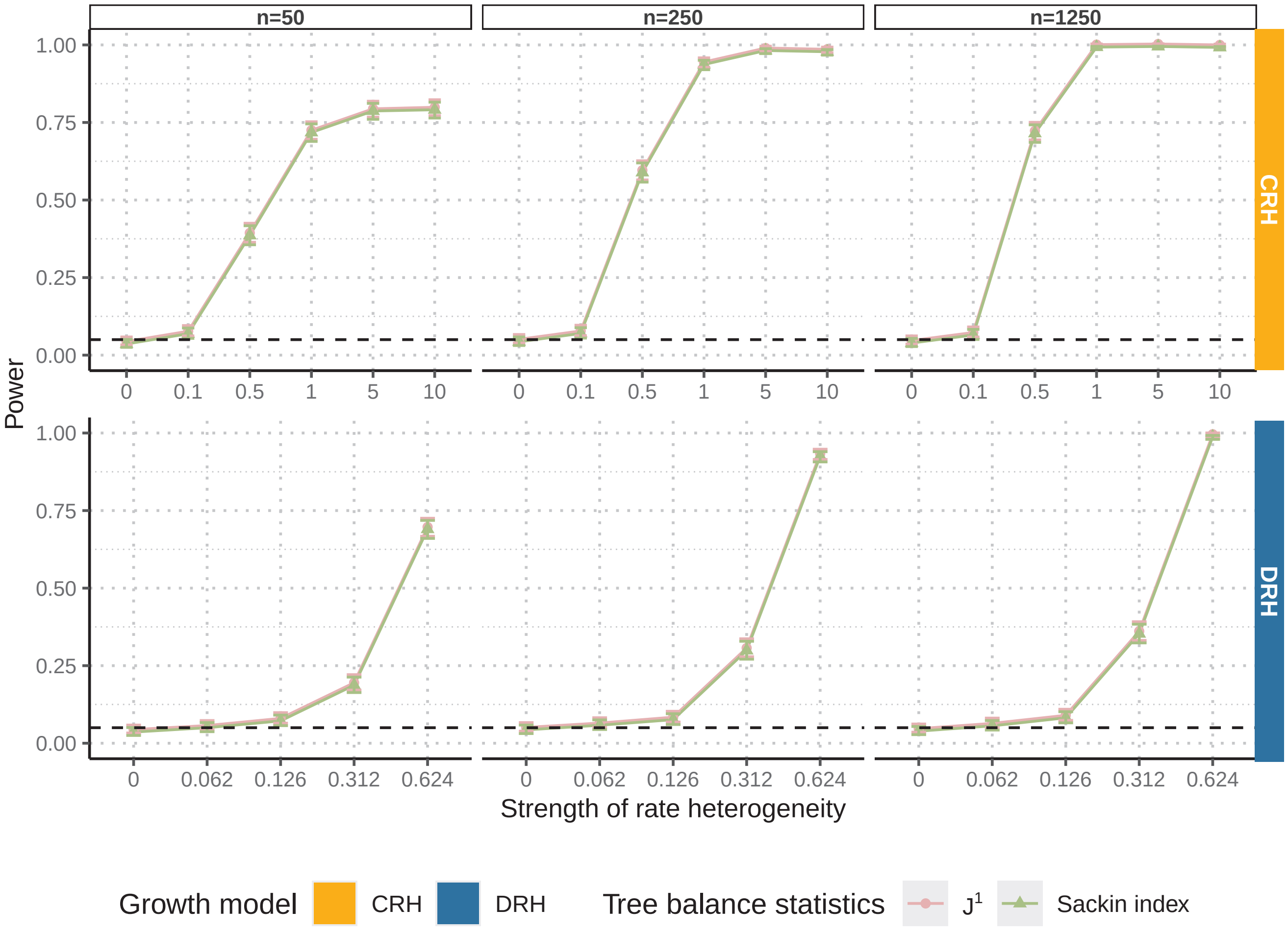

### Figure_S1.png

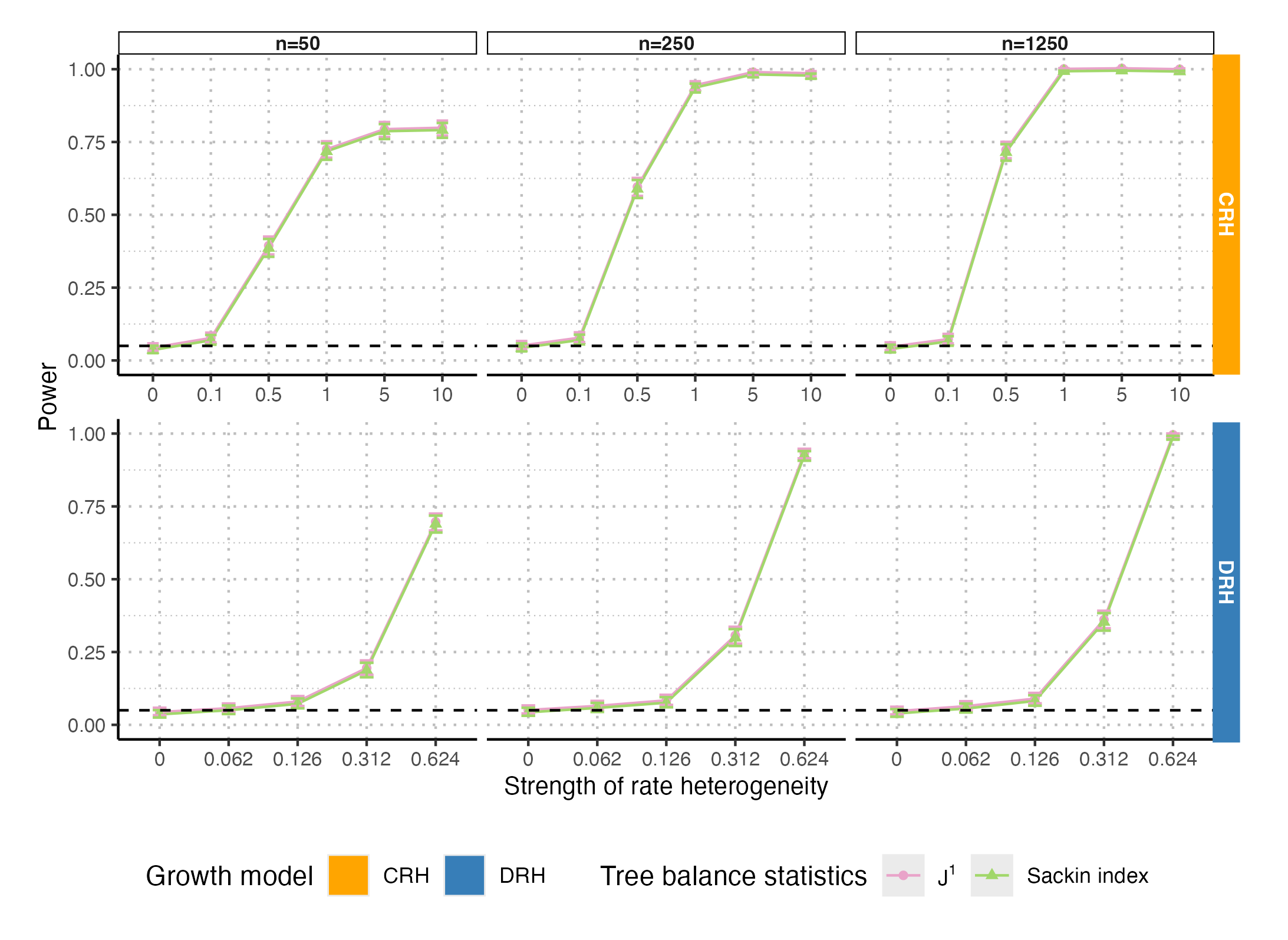

### Figure_S2.pdf

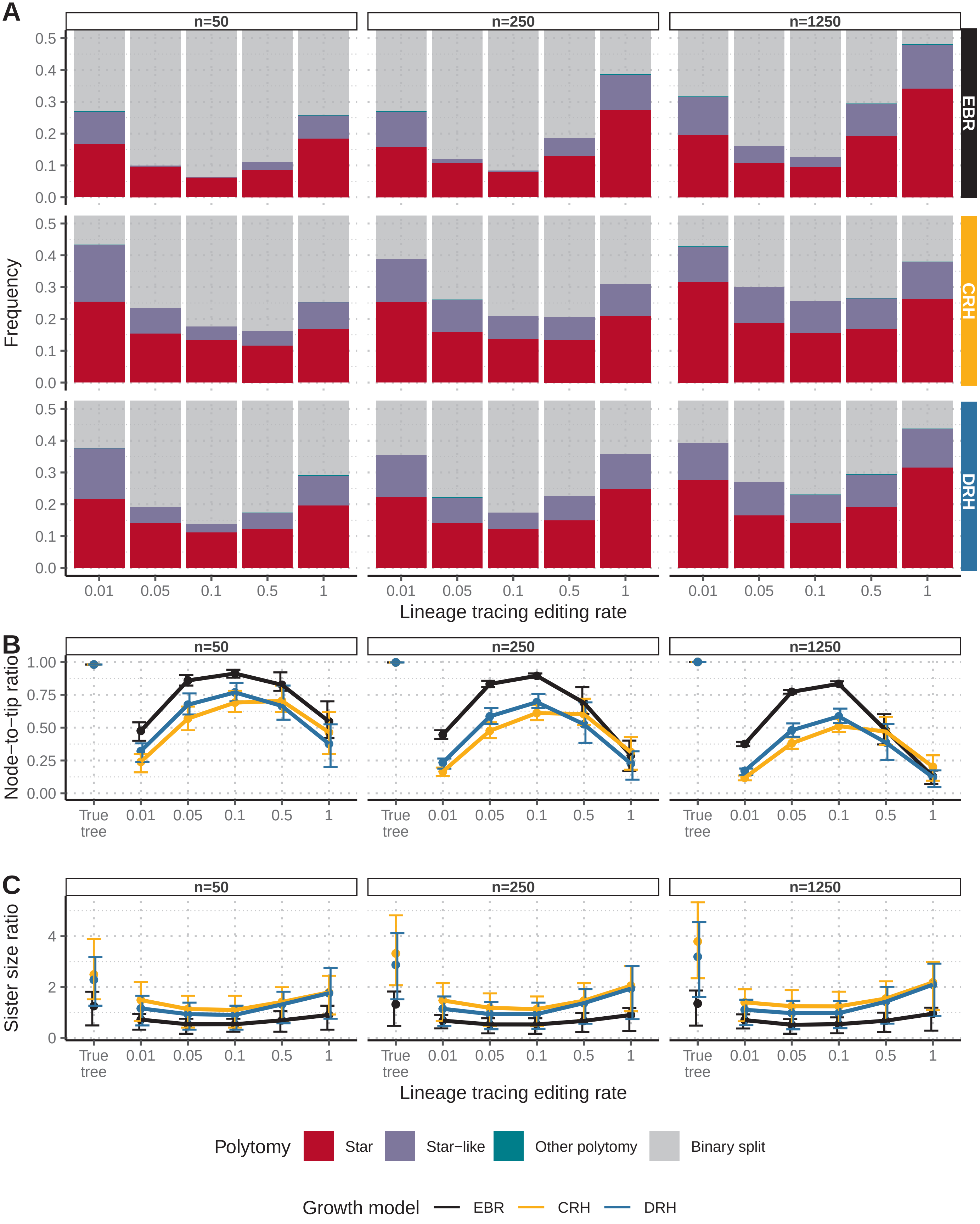

### Figure_S2.png

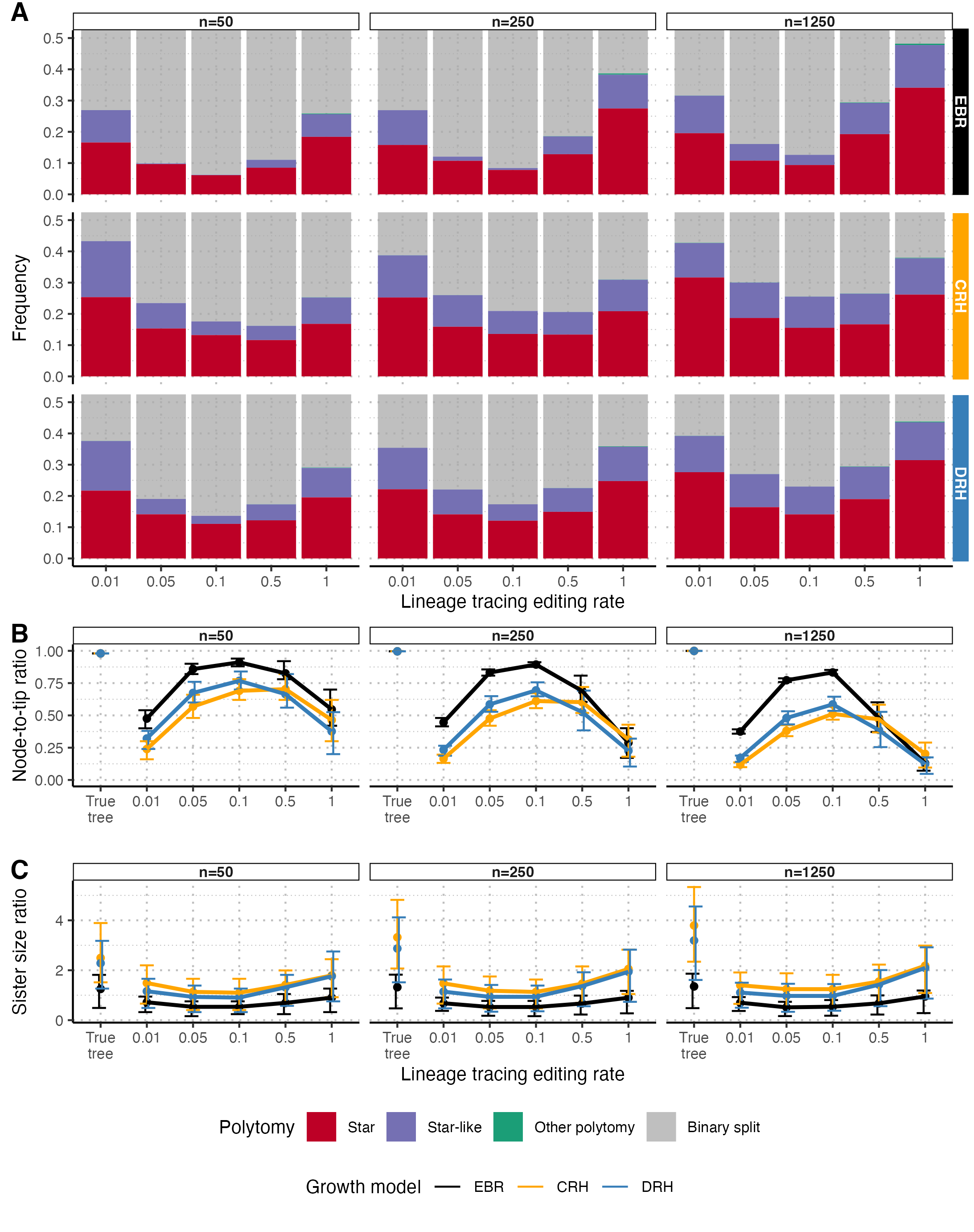

### Figure_S3.pdf

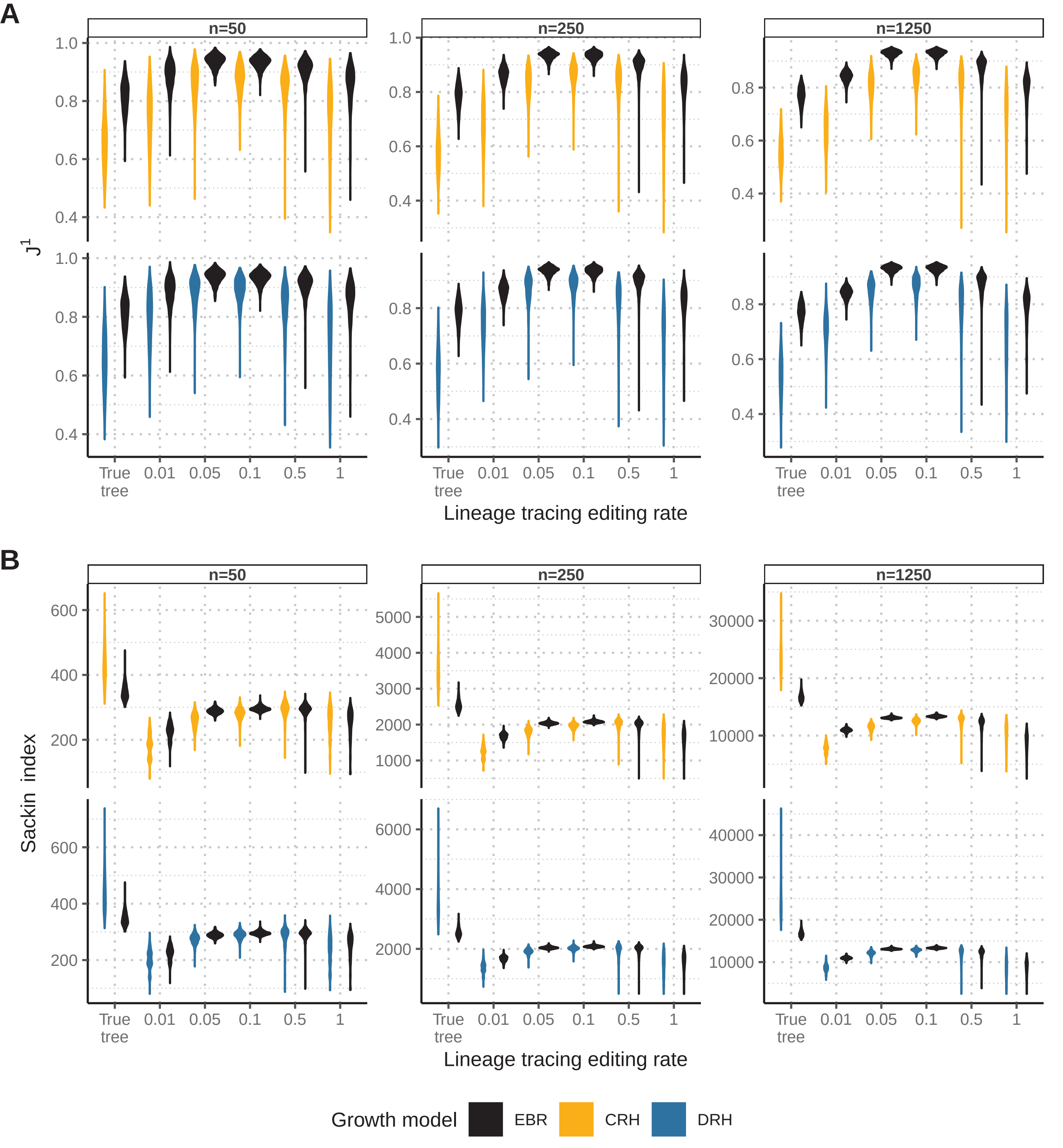

### Figure_S3.png

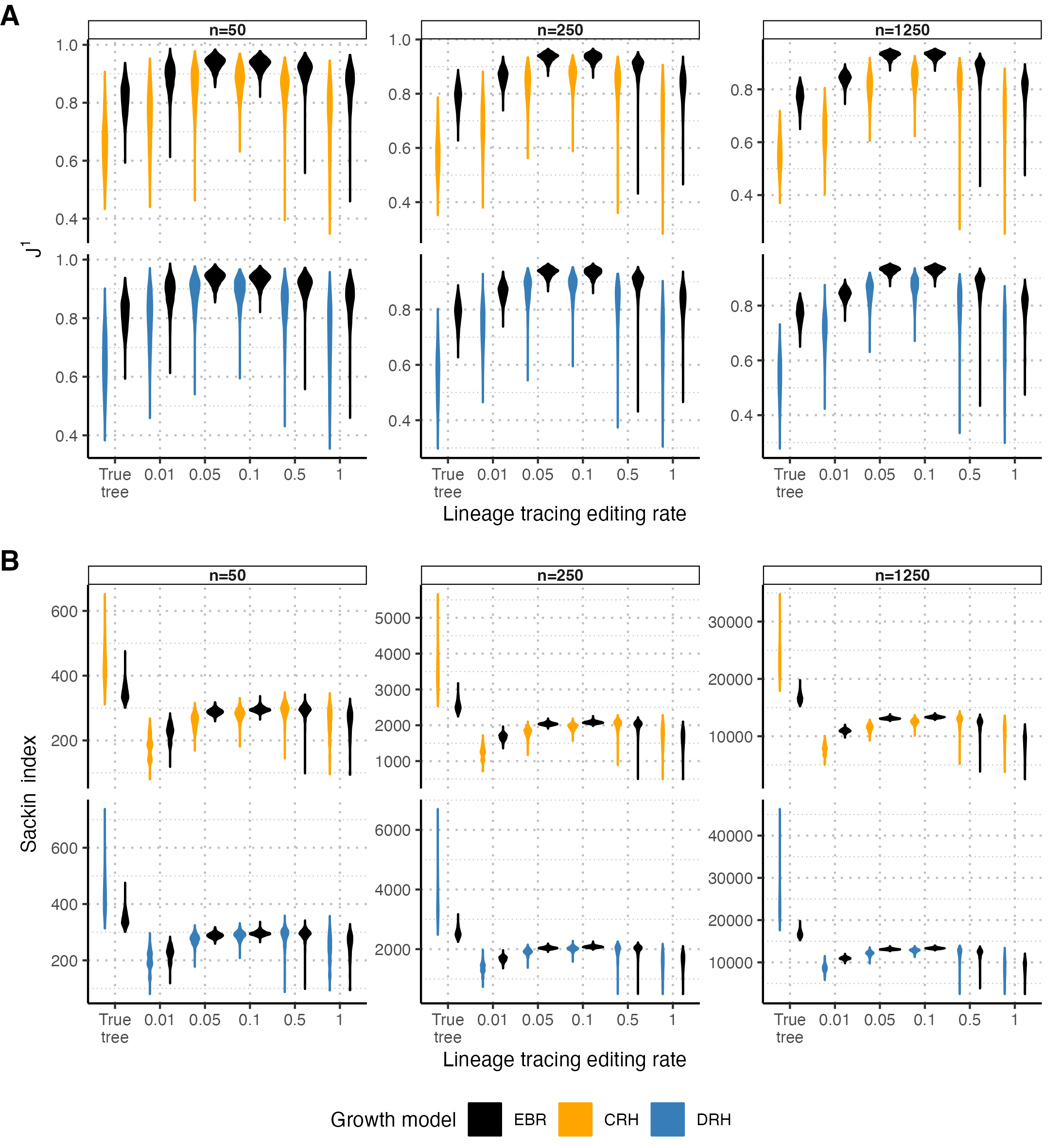

### Figure_S4.pdf

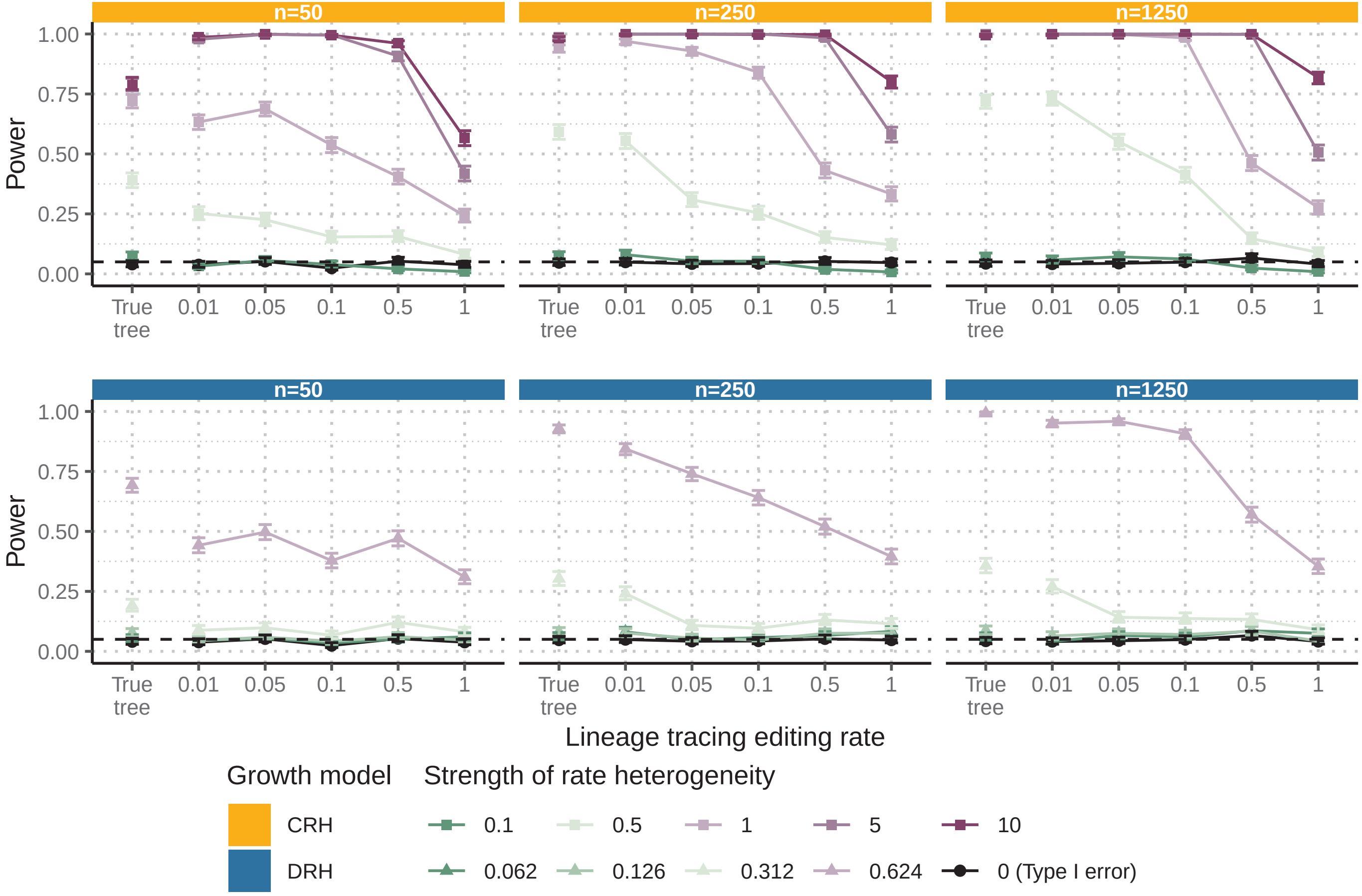

### Figure_S4.png

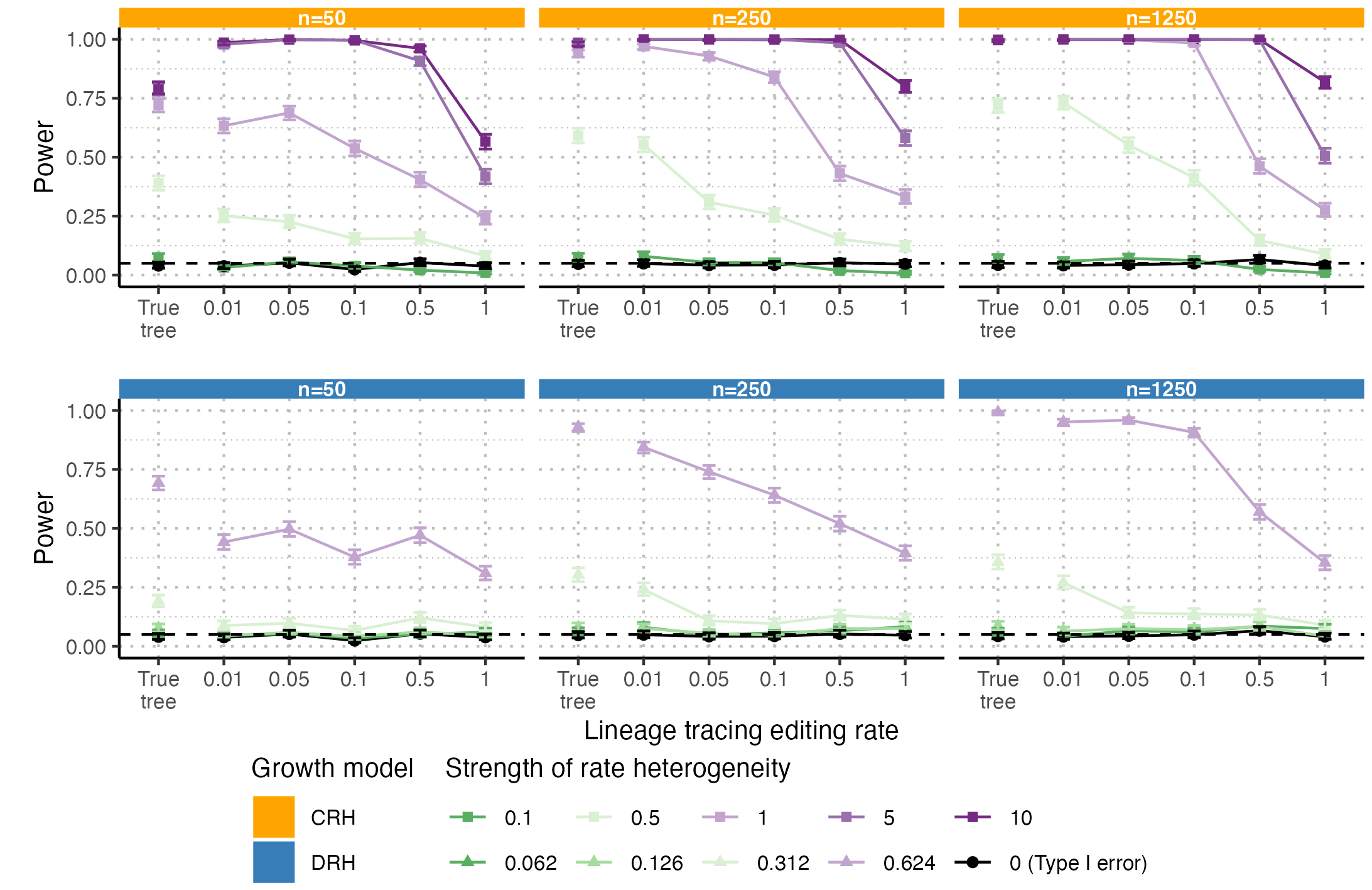

### Figure_S5.pdf

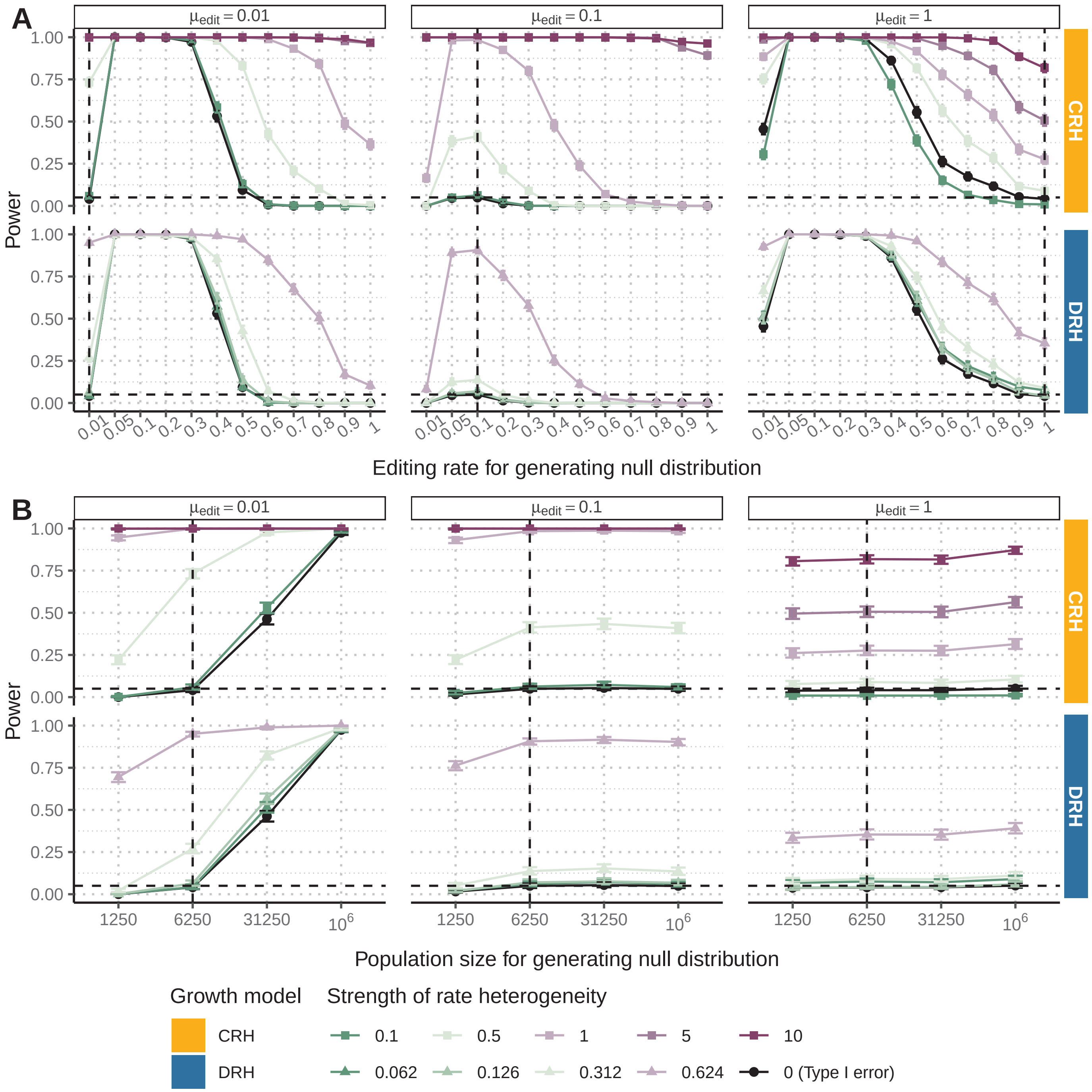

### Figure_S5.png

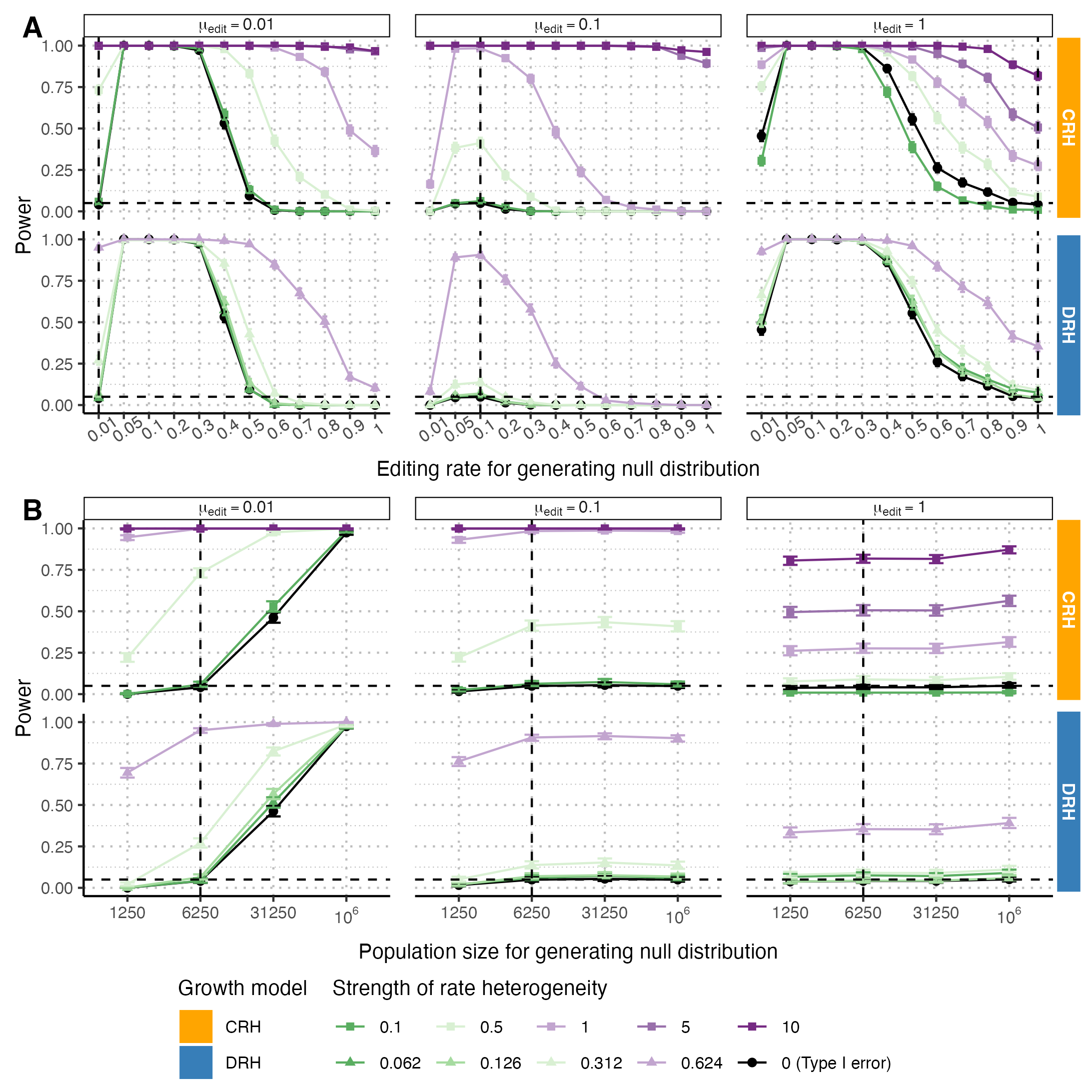

### Figure_S6.pdf

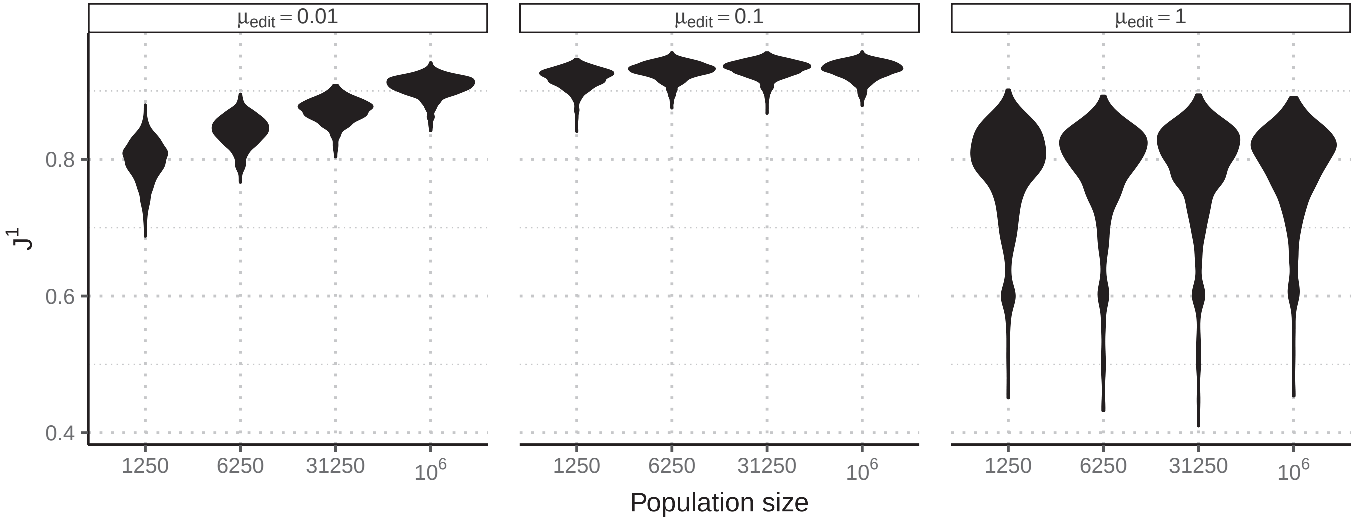

### Figure_S6.png

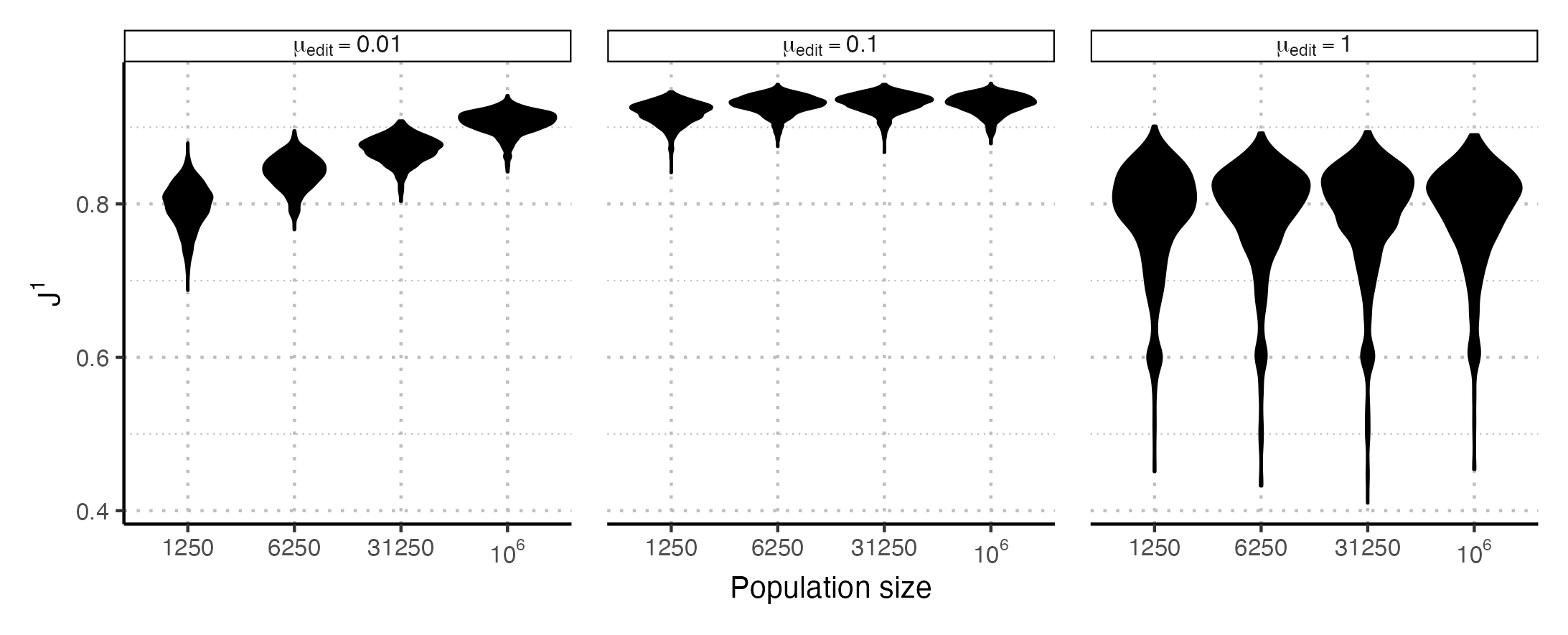

### Figure_S7.pdf

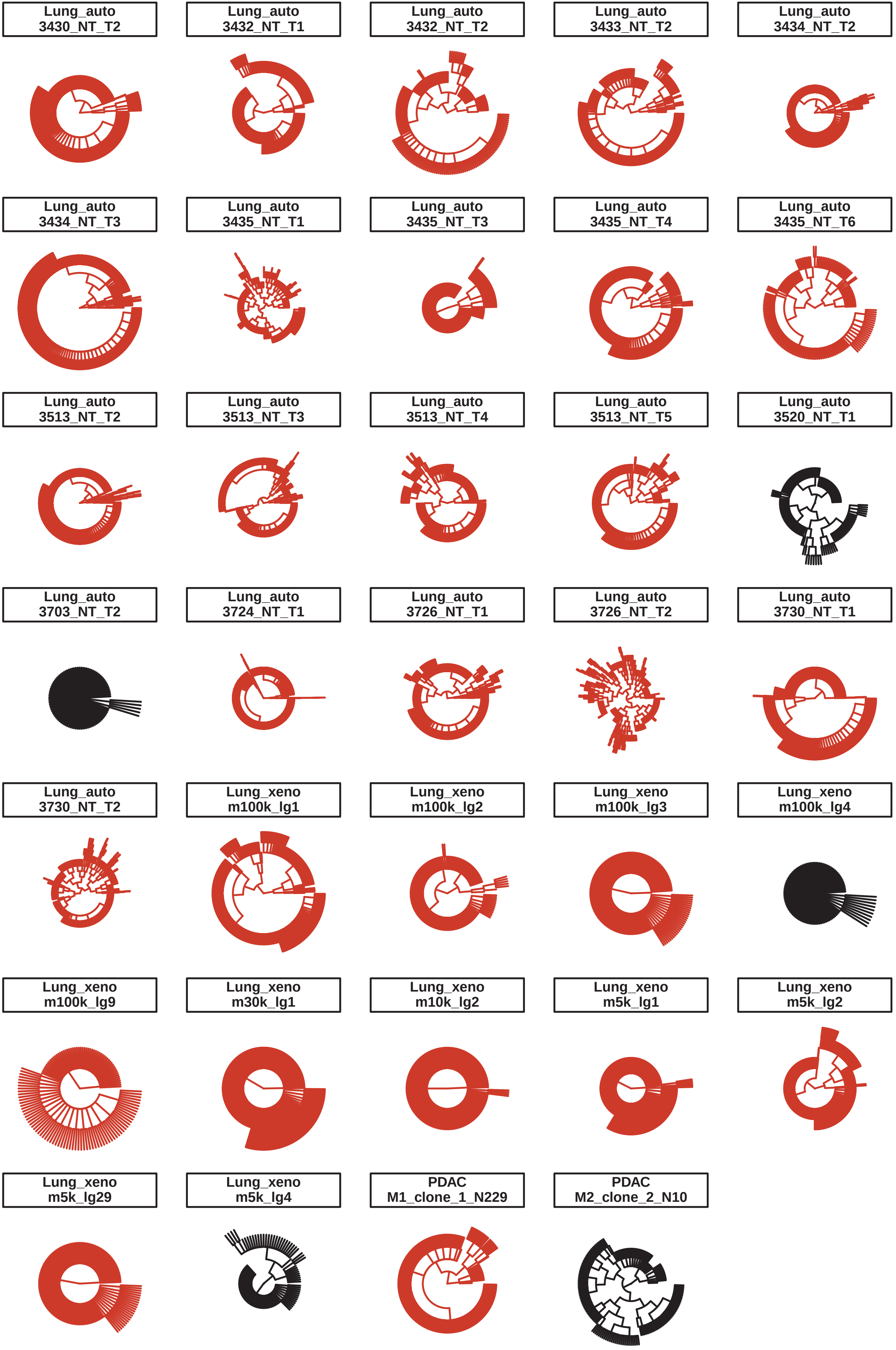

### Figure_S7.png

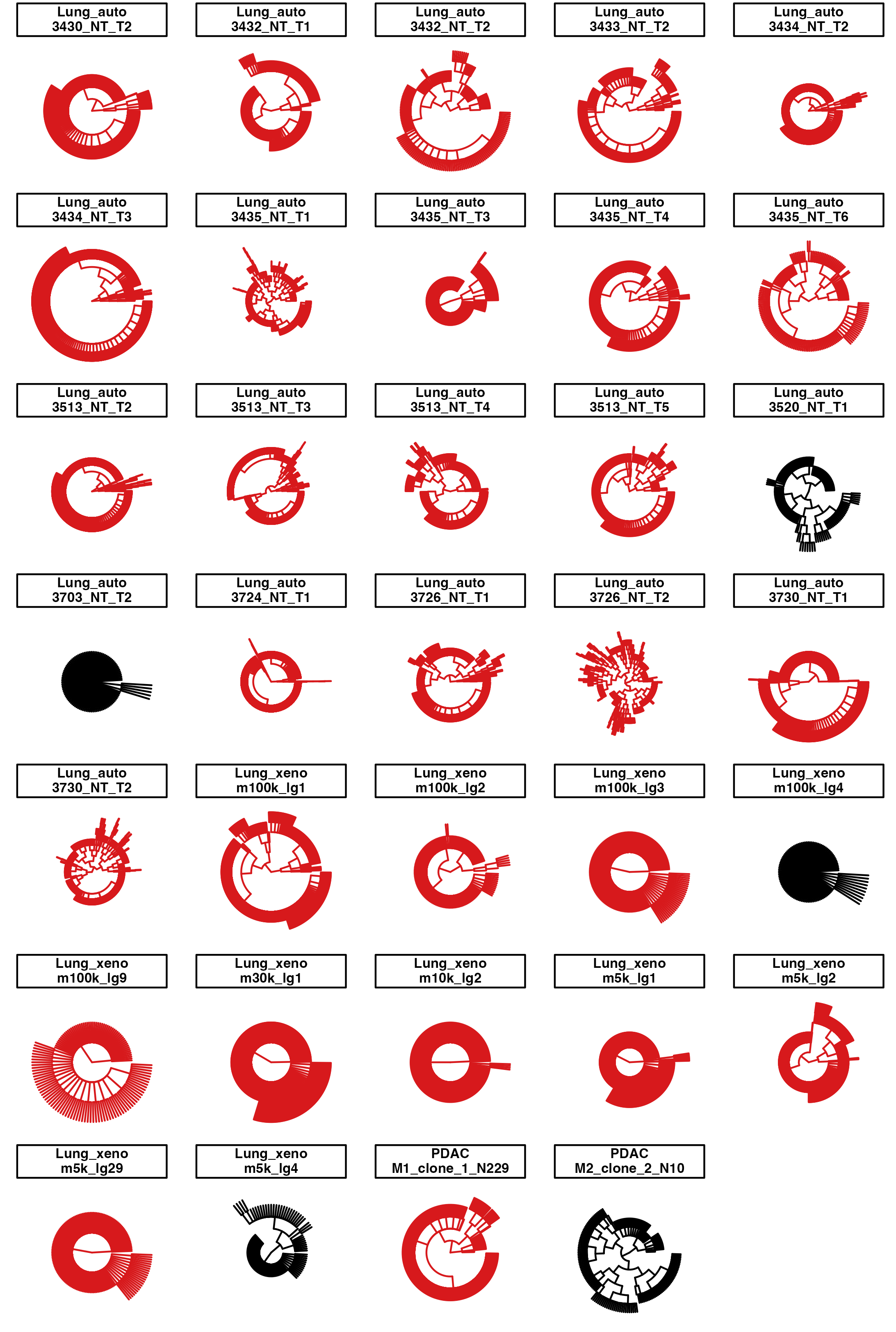
